# Analysing pneumococcal invasiveness using Bayesian models of pathogen progression rates

**DOI:** 10.1101/2021.09.01.458483

**Authors:** Alessandra Løchen, James E. Truscott, Nicholas J. Croucher

## Abstract

The disease burden attributable to opportunistic pathogens depends on their prevalence in asymptomatic colonisation and the rate at which they progress to cause symptomatic disease. Increases in infections caused by commensals can result from the emergence of “hyperinvasive” strains. Such pathogens can be identified through quantifying progression rates using matched samples of typed microbes from disease cases and healthy carriers. This study describes Bayesian models for analysing such datasets, implemented in an RStan package (https://github.com/nickjcroucher/progressionEstimation). The models converged on stable fits that accurately reproduced observations from meta-analyses of *Streptococcus pneumoniae* datasets. The estimates of invasiveness, the progression rate from carriage to invasive disease, in cases per carrier per year correlated strongly with the dimensionless values from meta-analysis of odds ratios when sample sizes were large. At smaller sample sizes, the Bayesian models produced more informative estimates. This identified historically rare but high-risk *S. pneumoniae* serotypes that could be problematic following vaccine-associated disruption of the bacterial population. The package allows for hypothesis testing through model comparisons with Bayes factors. Application to datasets in which strain and serotype information were available for *S. pneumoniae* found significant evidence for within-strain and within-serotype variation in invasiveness. The heterogeneous geographical distribution of these genotypes is therefore likely to contribute to differences in the impact of vaccination in between locations. Hence genomic surveillance of opportunistic pathogens is crucial for quantifying the effectiveness of public health interventions, and enabling ongoing meta-analyses that can identify new, highly invasive variants.

**Author summary:** Opportunistic pathogens are microbes that are commonly carried by healthy hosts, but can occasionally cause severe disease. The progression rate quantifies the risk of such a pathogen transitioning from a harmless commensal to causing a symptomatic infection. The incidence of infections caused by opportunistic pathogens can rise with the emergence of “hyperinvasive” strains, which have high progression rates. Therefore methods for calculating progression rates of different pathogen strains using surveillance data are crucial for rapidly identifying emerging infectious disease threats. Existing methods typically measure progression rates relative to the overall mix of microbes in the population, but these populations can vary substantially between locations and times, making the outputs challenging to combine across studies. This work presents a new method for estimating progression rates from surveillance data that generates values useful for modelling pathogen populations, even from relatively small sample sizes.

## Introduction

Opportunistic pathogens are commonly found in the environment, or microbiota of healthy individuals, but have the capacity to cause disease in some host organisms [1]. The frequency with which they transition from asymptomatic colonisation to symptomatic infection can be quantified as a progression rate [2]. Many opportunistic pathogens are diverse, and can be typed by a variety of phenotypic methods [3], or divided into strains using genotyping or whole genome sequencing [4, 5].

Understanding the variation in progression rates across such species is crucial for understanding changes in the incidence of diseases caused by opportunistic pathogens. Recently, strains with high progression rates, sometimes referred to as “hypervirulent” [6] or “hyperinvasive” [7], have been ascribed as the primary cause of rising case numbers of disease caused by *Neisseria meningitidis* [8], *Streptococcus pyogenes* [9] and *Klebsiella pneumoniae* [10], among others. However, understanding whether such strains drive elevated disease burdens through more rapid transmission, or heightened progression rates, typically requires additional information on the carried population of the opportunistic pathogen.

Progression rates have been extensively studied in the nasopharyngeal commensal and respiratory pathogen *Streptococcus pneumoniae* (the pneumococcus). Over one hundred serotypes have been identified in this species, each corresponding to a structurally-distinct polysaccharide capsule [11, 12]. These are clustered into 48 serogroups, based on serological cross-reactivity [13]. This capsule inhibits pneumococci being cleared by the immune system, and is crucial to their ability to cause invasive pneumococcal disease (IPD) [14, 15], infections of normally sterile anatomical sites. Hence the rate of progression from carriage to IPD is referred to as invasiveness [16]. It has long been assumed the capsule is an important determinant of pneumococcal invasiveness, based on differences in serotypes’ ability to cause disease in animal experiments [17]. Epidemiological differences are also apparent in human disease. Paediatric serotypes (serotype 14 and serotypes within serogroups 6, 9, 19, 23) were identified as being common in sporadic infant disease [18, 19].

Epidemic serotypes (such as 1, 2, 5 and 12F) were found to be capable of causing IPD outbreaks in adults [19], and therefore are likely to represent hyperinvasive types of *S. pneumoniae*.

Understanding these differences between serotypes has become crucial for *S. pneumoniae* epidemiology, owing to the introduction of polysaccharide conjugate vaccines (PCVs) for infant immunisation [20]. A seven valent PCV (PCV7) was introduced in the USA in 2000, which has been supplanted by ten-and thirteen-valent formulations (PCV10 and PCV13, respectively). The next generation of higher-valency vaccines (PCV15 and PCV20) will soon be introduced [21, 22]. PCVs induce immune responses against a specific subset of serotypes, usually resulting in their elimination from carriage [23]. This contrasts with the 23-valent polysaccharide vaccine (PPV23) administered to older adults, which only protects against symptomatic disease [20]. However, the overall frequency of *S. pneumoniae* colonisation typically remains stable after any PCV’s introduction, owing to the replacement of vaccine serotypes by the plethora of non-vaccine serotypes [24, 25]. Nevertheless, PCVs have usually proved highly effective at reducing IPD through facilitating the replacement of the pre-PCV carried population with pneumococcal serotypes associated with lower invasiveness [26, 27]. Hence optimal PCVs are those which minimise the overall invasiveness of the carried pneumococcal population [28, 29]. Therefore estimation of invasiveness across vaccine-targeted serotypes, and those which may replace them post-PCV, is vital for reducing the incidence of IPD. Such quantification of invasiveness is typically achieved through paired studies of serotypes’ prevalence in geographically and temporally matched surveys of pneumococcal carriage, and surveillance of IPD.

Multiple methods have been used to estimate invasiveness using such paired case and carrier data. The most common approaches use odds ratios: the ratio of isolates of a given serotype recovered from disease against those recovered from carriage, divided by the same ratio calculated across all other serotypes in the population [30, 31]. However, this statistic is intrinsically imperfect, because even if a serotype causes disease at a consistent rate across populations, the mix of serotypes against which it is compared will differ between locations [32, 33]. This geographical heterogeneity has been exacerbated by post-PCV serotype replacement, which has driven increasing divergence between countries’ serotype compositions [34]. The interpretability of the odds ratios may be improved by standardising them relative to the geometric mean across all odds ratios [35], but this does not resolve the underlying problem, which likely contributes to the substantial heterogeneity observed for some serotypes when odds ratios are combined in meta-analyses [36]. This means carefully-conducted odds ratio analyses can be hampered by having to subsample data, and employ both fixed and random effects analyses, within a single analysis [37]. One solution has been to estimate invasiveness relative to a standard serotype that is common in both carriage and IPD across studied locations [36, 38]. However, these properties mean such serotypes are likely to be targeted by PCVs. Correspondingly, the original standard, serotype 14 [31], has been eliminated by PCV7 in many settings. Similarly, PCV13 has removed the replacement standard serotype used in post-PCV7 studies, 19A [36].

Invasiveness has been more directly characterised as the ratio of disease cases relative to the estimated prevalence in carriage [27]. However, this approach did not account for the stochasticity of disease cases being observed, and assumed the uncertainty associated with the estimated case-to-carrier ratio derived only from the colonisation survey data. More comprehensive quantification of the uncertainty in invasiveness can be achieved with Bayesian frameworks, which have been applied to individual pneumococcal populations [2, 39]. Invasiveness has been modelled by jointly fitting distributions to carriage and disease data, then using random effects models to estimate invasiveness in different age groups [39]. An alternative approach, using Poisson regression, was used to estimate serotypes’ progression rates for causing otitis media [2].

In this work, we apply Bayesian modelling to estimate progression rates from meta-analyses of multiple studies. The ability to synthesise data across populations is important for maximising the available information on each type, which may be infrequently observed in individual studies. This is particularly important in *S. pneumoniae*, as many genotypes that have emerged post-PCV were previously rare, and vary geographically in their prevalence [34, 40]. The model enables the estimation of progression rates as an opportunistic pathogen’s hazard of progressing from carriage to disease in a specified population. Such absolute values avoid a measure that is relative either to the rest of the local microbial population, or to one standard type. These properties are crucial for quantifying and modelling changes in disease incidence, such as following vaccine introduction, or the emergence of a hyperinvasive strain. They can be particularly valuable in settings where disease surveillance is not comprehensive, and only carriage data are available [39, 41].

Using a Bayesian approach also enables hypothesis testing through statistical comparison of alternative model structures [2, 39]. This is crucial for understanding the host and pathogen factors that affect progression rates. For instance, the correlation between pneumococcal serotypes and invasiveness has not been indisputably established as a causal link. In *Haemophilus influenzae*, changes to an isolate’s serotype altered its virulence in an animal model of disease in such a manner that reflected the epidemiology of human disease [42]. While some equivalent experiments in *S. pneumoniae* have replicated observations from human IPD [43, 44], others have found changes in an isolate’s serotype did not change its invasiveness in an animal model [45, 46], suggesting serotype-independent factors may contribute to this phenotype. Studies of isolates from disease and carriage using comparative genomic hybridisation [47, 48] and whole genome sequencing [49, 50] have both supported links between non-capsular loci and invasiveness. An unambiguous association cannot even be made between the distinctive phenotypes of the epidemic serotypes and their capsules, as these pneumococci tend to have low genetic diversity [37, 51], and therefore their capsule polysaccharide synthesis locus is in linkage disequilibrium with many other variable genomic loci [52, 53].

Epidemiological studies using genotyped and serotyped isolates have also found examples of within-serotype differences in invasiveness [35,54,55], although other studies found evidence of serotype being the primary determinant of invasiveness [31]. Genomic analysis by the Global Pneumococcal Sequencing project also identified differences in the invasiveness of strains of the same serotype [33]. However, given the multiple pairwise comparisons conducted in such analyses, it is not clear whether such observations might be expected, even if the capsule is the primary determinant of invasiveness. Such questions can be addressed through comparison of different model fits to the data, thereby improving our understanding of the determinants of progression rates. This is essential for deciding on the most effective methods for ongoing pathogen surveillance, and enabling the most efficient use of such data. Using appropriately-structured models, capable of combining information from multiple studies at the most informative level of resolution, will help identify emerging highly invasive types at the earliest possible opportunity. Therefore the methods, models and data described in this study are made available as an R package (https://github.com/nickjcroucher/progressionEstimation), to enable the continual aggregation of case and carrier studies for any opportunistic pathogens.

## Methods

### Model definitions and assumptions

All models were designed to be applied to a meta-population of a multi-strain opportunistic pathogen. Each population *i* corresponded to a matched sample of microbes from asymptomatic carriers, or the environment, and a sample from disease. It was assumed all cases of disease emerged independently from the carried or environmental population. Across the meta-population, the microbes were classified according to their type *j*, strain *k*, or the combination of both, *j*,*k*. The “type” may be defined by any phenotypic categorisation; analyses of *S. pneumoniae* typically use serotyping. The “strain” can represent any coherent genetic subdivision of the population; for instance, analyses of *S. pneumoniae* can use the Global Pneumococcal Sequence Clusters (GPSCs) [37], but alternatively-defined “clades” or “lineages” could be employed instead. If *x* denotes any one of these classifications, then the carriage rate in population *i*, ρ*_i,x_*, was estimated based on the total number of samples (including negative results), η*_i_*, and the number of samples positive for isolates of category *x*, *c_i,x_*:

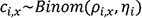

Modelling the number of positive samples using a Binomial distribution follows the precedent of Weinberger *et al* [39], and assumes all carriage samples are independently drawn from the circulating population.

A set of related statistical models were used to estimate the progression rate [2], ν, the hazard of developing a disease, given carriage of a microbe per unit time. As ν was assumed to be constant for each type *x* in each study *i*, the number of observed cases of disease caused by *x* in *i*, *d_i,x_*, was modelled as the product of the number of potential hosts in *i*, *N_i_*; the proportion carrying *x* in *i*, ρ*_i,x_*; the per unit time probability of *x* causing disease, ν*_x_*; and the duration of the study, *t_i_*. . Hence, the expected number of isolates from disease, IE[*d_i,x_*], in all models was proportional to all these quantities:

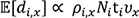

Some models also adjusted the expected *d_i,x_* between *i* through a scaling parameter, γ_i_, that was relative to a reference population. This corrects for differences between studies both in the actual hazard of developing a disease, due to host population variation, and differences in the comprehensiveness of surveillance across the study populations. Hence, the expected number of isolates from disease, IE[*d_i,x_*], in such models was proportional the quantities:

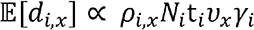

All these parameters were assumed to be constant within a study over *t_i_*. All disease cases were assumed to develop independently, and therefore were modelled as a Poisson process. However, relevant unmodelled variation, or oversampling of local transmission chains in either carriage or disease surveillance [56], could cause heterogeneity in the observed isolate counts in either component of the paired samples. Hence *d_i,x_* was additionally modelled as following a negative binomial distribution, in which overdispersion was quantified by the precision parameter, 0, relative to the mean, μ Lower 0 values indicate greater overdispersion, as the variance of the distribution was calculated as:

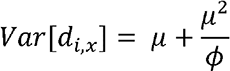

### Models of disease prevalence

Four different structures were used to model IE[*d_i,x_*]. For each, two different versions were fitted to the data. In the simpler version, the observed disease case counts were assumed to follow a Poisson distribution:

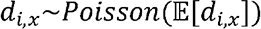

The more complex version assumed a negative binomial distribution of the observed disease case counts:

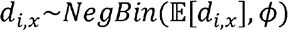

### Null model

In this simplest model, the progression rate was independent of the pathogen’s type (ν*_x_* = ν for all *x*) and host population (γ*_i_* = 1 for all *i*). Therefore *d_i,x_* is expected to be correlated with *c_i,x_*:

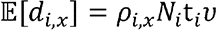

### Type-specific model

This model, similar to that Weinberger *et al* applied to a single population [39], allowed for variation in progression rate between types, but still assumed host populations to be homogeneous (γ*_i_* = 1 for all *i*). Therefore *d_i,x_* depends on both the carriage prevalence and progression rate of *x*:

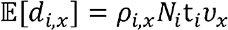

### Study-adjusted model

In this model, variation in disease prevalence between studies reflected differences in host population and surveillance, rather than differences between types. Hence the progression rate was independent of the pathogen’s type (ν*_x_* = ν for all *x*), but varied between *i* due to γ*_i_*:

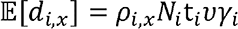

### Study-adjusted type-specific model

This model allowed for both variation in the progression rate between types *x* and study populations *i*:

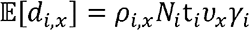

### Joint modelling of strain and type progression rates

For datasets in which information was available on both type (for *S. pneumoniae*, serotype) *j* and strain *k*, ν*_x_* was modelled in five different ways:

### Serotype-determined progression rate

The progression rate was entirely determined by an isolate’s type, and independent of its strain background (*v_x_* = *v_j_*).

### Strain-determined progression rate

The progression rate was entirely determined by the strain to which an isolate belonged, and independent of its type (*v_x_* = *v_k_*).

### Strain-modified type-determined progression rate

Two approaches were taken to modelling progression rates as being independently affected by type and genetic background. The first modelled the progression rate as being primarily determined by an isolate’s type, which was modified by its strain background. This was calculated as *v_x_* = *v_j_v_k_*, where the model priors meant *v_j_* was less constrained than *v_k_*. Hence most variation in *v_x_* should be attributed to the type *j*.

### Type-modified strain-determined progression rate

The second approach to analysing the independent effects of type and genetic background modelled progression rate as being primarily determined by the strain to which an isolate belonged, which was modified by its type. This was calculated as *v_x_* = *v_j_v_k_*, where the model priors meant *v_k_* was less constrained than *v_j_*. Hence most variation in *v_x_* should be attributed to the strain *k*.

### Type and strain-determined progression rate

Each combination of type *j* and strain *k* was modelled as having a unique progression rate (*v_x_* = *v_j,k_*), suggesting non-multiplicative interactions between strain background and type on an isolate’s progression rate.

### Model priors and implementation

The characteristics of each study population (η*_i_*, *N_i_*, *t_i_*) and observed counts of type *x* in carriage (*c_i,x_*) and disease (*d_i,x_*) were assumed to be accurate and without error. As each recovery of a type *x* isolate from carriage contributes to the estimation of ρ*_i,x_*, even if an individual carries multiple types, the prior distribution was:

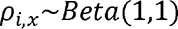

This means each type’s carriage was independently estimated, and the sum of carriage prevalences (∑) may be greater than one, if there is high level of multiple carriage in a population. The beta distribution was used, as it is the conjugate prior of the binomial distribution, although setting the scale and shape parameters to one make it equivalent to a uniform distribution bounded by zero and one.

Invasiveness has been estimated to vary over orders of magnitude in *S. pneumoniae* [29], and therefore a uniform prior was placed on the logarithm of *v_x_* in all models. The lower bound was set at one IPD case per million carriers per unit time, as the largest studies used surveillance of host populations consisting of tens of millions of individuals (Table S1). The upper bound was ten cases per carrier per unit time, which would effectively correspond to an obligate pathogen for a bacterium with the carriage duration of *S. pneumoniae* [57]:

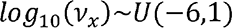

**Table 1.**
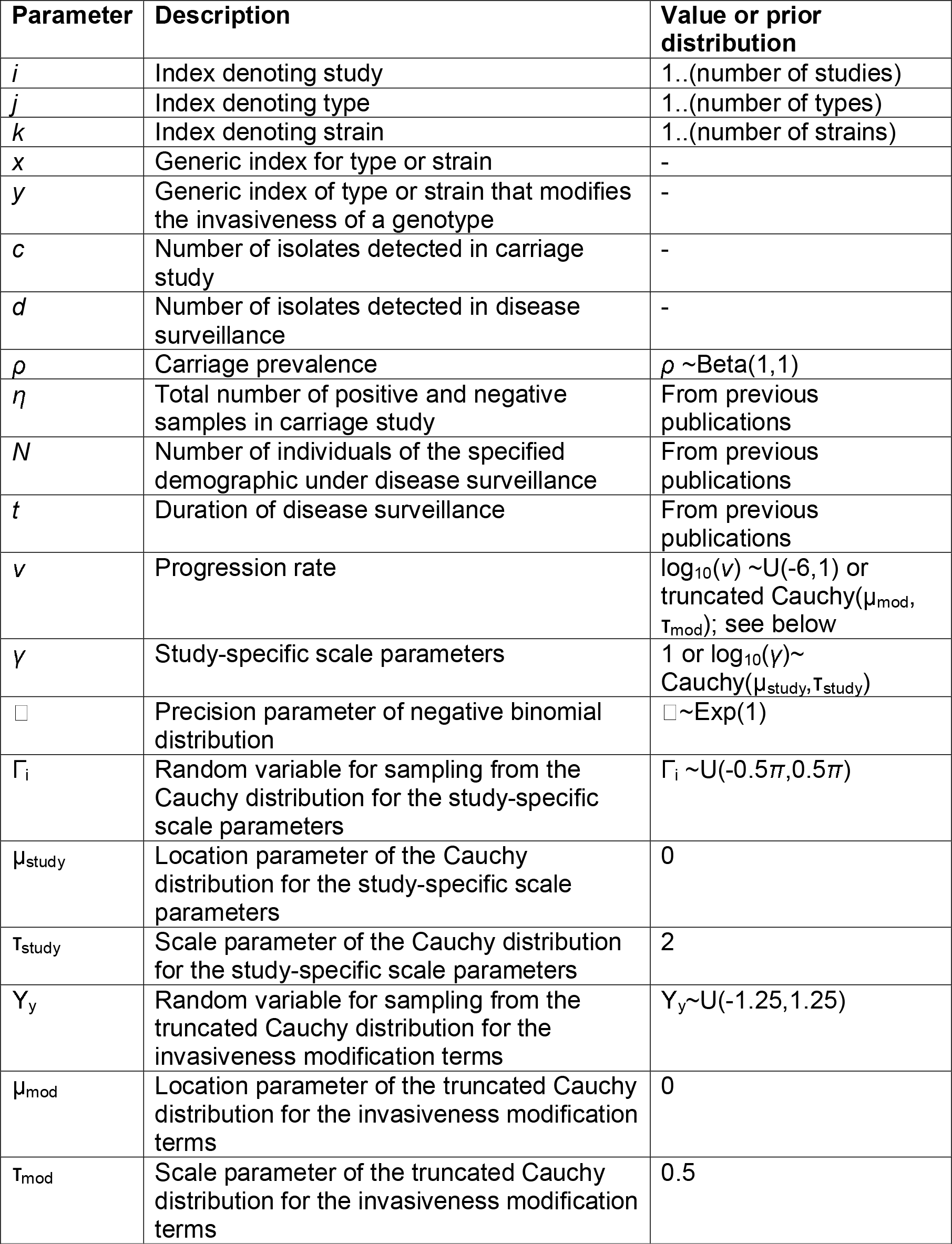
Summary of parameters used in the models.

As the scaling factors were relative measures of disease prevalence between populations, γ*_i_* for the population with the largest sample size was fixed at one. The scaling factor for each other population *i* was allowed to vary higher or lower. The Cauchy distribution was assumed as the prior for the logarithm of γ*_i_*, as this describes the ratio of two normally-distributed random variables. However, the Hamiltonian Monte Carlo algorithm used for model fitting does not efficiently sample heavy-tailed distributions, and therefore the prior was reparameterised to use a random variable Γ_i_ with a unit-sized uniform prior [58]:

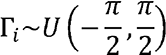

The Γ_i_ values were then transformed to generate γ_i_ values with a Cauchy distribution using the location ( = 0) and scale (: = 2) parameters:

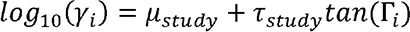

The : value allowed for variation in observed incidence over multiple orders of magnitude between datasets, ensuring the model makes allowance for substantial heterogeneity in observed disease incidence [59].

A similar reparameterization of the Cauchy distribution was used for the models in which type and strain background independently contributed to a genotype’s progression rate (the strain-modified type-determined, and type-modified strain-determined, progression rate models). The categorisation modelled as determining the progression rate (*v_x_*) had the same prior as when a single factor determined the progression rate. This was multiplied by a second value, *v_y_*, determined by the classification modifying the progression rate. The prior distribution for the logarithm of this value was a truncated Cauchy, symmetrical around zero. This represented the expectation that the modification of the progression rate would be small (approximately one) in most cases, but may be substantial in rare instances. The truncation was found to be necessary for efficient sampling with the Hamiltonian Monte Carlo algorithm. This was parameterised with a random variable Υ_y_, sampled from a uniform distribution with narrower boundaries than those for the analogous Γ_i_ variable:

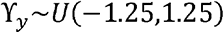

The Υ_y_ values were then transformed to generate *v_y_* values using the location ( = 0) and scale (: = 0.5) parameters:

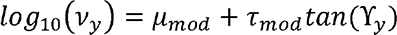

This meant *v_y_* values could vary over three orders of magnitude (between 0.0313 and 32.0), while enabling models to converge with reasonable MCMC lengths.

The precision parameter of the negative binomial distribution, 0 an exponential distribution. Values of 0 that are large relative to IE, suggest there is little evidence of overdispersion, and therefore these parameter values should be associated with small prior probabilities. As IE, must always be close to, or above, one in case-carrier studies, an exponential rate parameter of one was used to reduce the likelihood of the negative binomial model when 0 was above this value. Yet this broad, always positive, distribution enabled estimation of smaller, albeit non-zero, 0 values for representing highly overdispersed data:

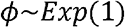

Model parameters are summarised in Table 1. All models were implemented using R [60] and Rstan [61]. Each model was fitted using two Hamiltonian Markov chain Monte Carlo (MCMC) evaluations for 25,000 iterations. Convergence was assessed through analysing the MCMC traces, using the criteria of requiring all is values to be below 1.05, and no reports of divergent transitions in the sampling [58]. This was achieved using default MCMC sampling parameters for the serotype analyses, but the strain analyses required a smaller step size (0.01), higher acceptance probability (0.99) and larger tree depth (20). The data and code used for the analyses are available as an R package from https://github.com/nickjcroucher/progressionEstimation/.

### Model evaluation and comparison

The ability of models to recover known parameter values from synthetic datasets was tested by simulation, as described in Text S1. Models were compared using likelihoods, independent of priors, through approximate leave-one-out cross-validation with the loo package [62]. The marginal likelihoods of different models, given the data, were compared with Bayes factors calculated using the bridgesampling package [63]. The same number of iterations were used to calculate the logarithmic marginal likelihoods as were used to fit the models. Evaluation of our results against odds ratio calculations used the metafor package [64], as described previously [29]. Model outputs were analysed and plotted with the tidyverse and ggpubr packages [65, 66].

#### S. pneumoniae serotype data

The matched carriage and invasive pneumococcal disease serotype datasets were combined from two meta-analyses [29, 36]. Twelve of the reported studies were omitted due to lack of publicly-available data [31,67–72]; difficultly in defining independent cross-sectional carriage samples [73, 74]; small sample sizes once stratified by age [75], or study design biased towards particular serotypes [76]. If a PCV were introduced during a study, then samples were divided into pre-PCV and post-PCV datasets. This resulted in 21 systematically-sampled and comprehensively serotyped paired asymptomatic carriage and disease samples (Table S1). Of these studies, 20 included data on child carriage and child IPD, and five included data on child carriage and adult IPD.

Data that were only resolved to serogroup level, based on the currently-known serotypes, were omitted from the analysis, although the overall number of samples taken (η*_i_*) was not adjusted. However, data on serotypes in historical studies that are now known to correspond to multiple serotypes (e.g. serotype 6A now being resolved into 6A and 6C [77]) were included unaltered. Additionally, isolates of serotypes 15B and 15C were combined into the single 15B/C category.

#### S. pneumoniae strain data

Analysis of variation of invasiveness between *S. pneumoniae* strains in children used population genomic data from South Africa [37], a mixture of population genomic and genotyped data from the USA [37], and genotyped data from Finland [74], Oxford [31] and Stockholm [78]. The equivalent analysis of strain invasiveness using isolates mainly from adult disease used data from Portugal [54]. Some of the genotyped datasets were not as thoroughly documented as the serotyped datasets, and therefore it was not appropriate to fit models that lacked a study-specific scale factor that reduced their reliance upon accurate carriage sample information (Text S2).

## Results

### Bayesian meta-analysis of *S. pneumoniae* serotype invasiveness

Twenty matched IPD case and nasopharyngeal carriage datasets were used to quantify *S. pneumoniae* serotype invasiveness in children (Table S1). From these, 7,340 carriage isolates and 2,851 disease isolates were extracted, across 72 serotypes (Fig. S1). Multiple models of invasiveness were proposed to analyse these data (see Methods). The null model assumed there was no systematic difference between serotypes’ invasiveness across datasets. The type-specific model assumed each serotype had a characteristic invasiveness that was consistent across locations. The study-adjusted null model assumed invasiveness did not vary across serotypes, but could differ between studies. Finally, the study-adjusted type-specific model allowed invasiveness to vary between both types and studies. All four approaches to estimating the expected numbers of disease cases and carriage isolates were fitted assuming the number of disease cases would follow a Poisson distribution, and allowing for overdispersion by fitting a negative binomial distribution to these values, resulting in eight total models. These models were able to accurately recover progression rates (Fig. S2) and infer overdispersion (Fig. S3) from simulated data (Text S1), with the true values used to generate the synthetic datasets lying within the 95% credibility intervals of the estimates from the fit of the corresponding model.

There was limited power for distinguishing the different model structures when the counts of both carriage and disease isolates were low for a given type in a study. Therefore paired observations from an individual study were only used for the model comparison if either the number of carried, or disease, isolates was at least five. This reduced the dataset to 7,048 carriage isolates and 2,617 disease isolates across 45 serotypes (Fig. S4). The eight models were each fitted to the data using two Hamiltonian Markov chain Monte Carlo samplings, both run for 25,000 iterations. All chains appeared to have converged, based on the traces of the posterior probabilities (Fig. S5) and all is values being below 1.001 (Fig. S6). Comparisons of the observed and predicted isolate counts showed all models could accurately reproduce the observations from carriage studies, with the exception of the null and study-adjusted Poisson models (Fig. 1). However, only the most complex models, adjusting for both *S. pneumoniae* serotype and study, were able to accurately reproduce the disease case counts.

**Figure 1.**
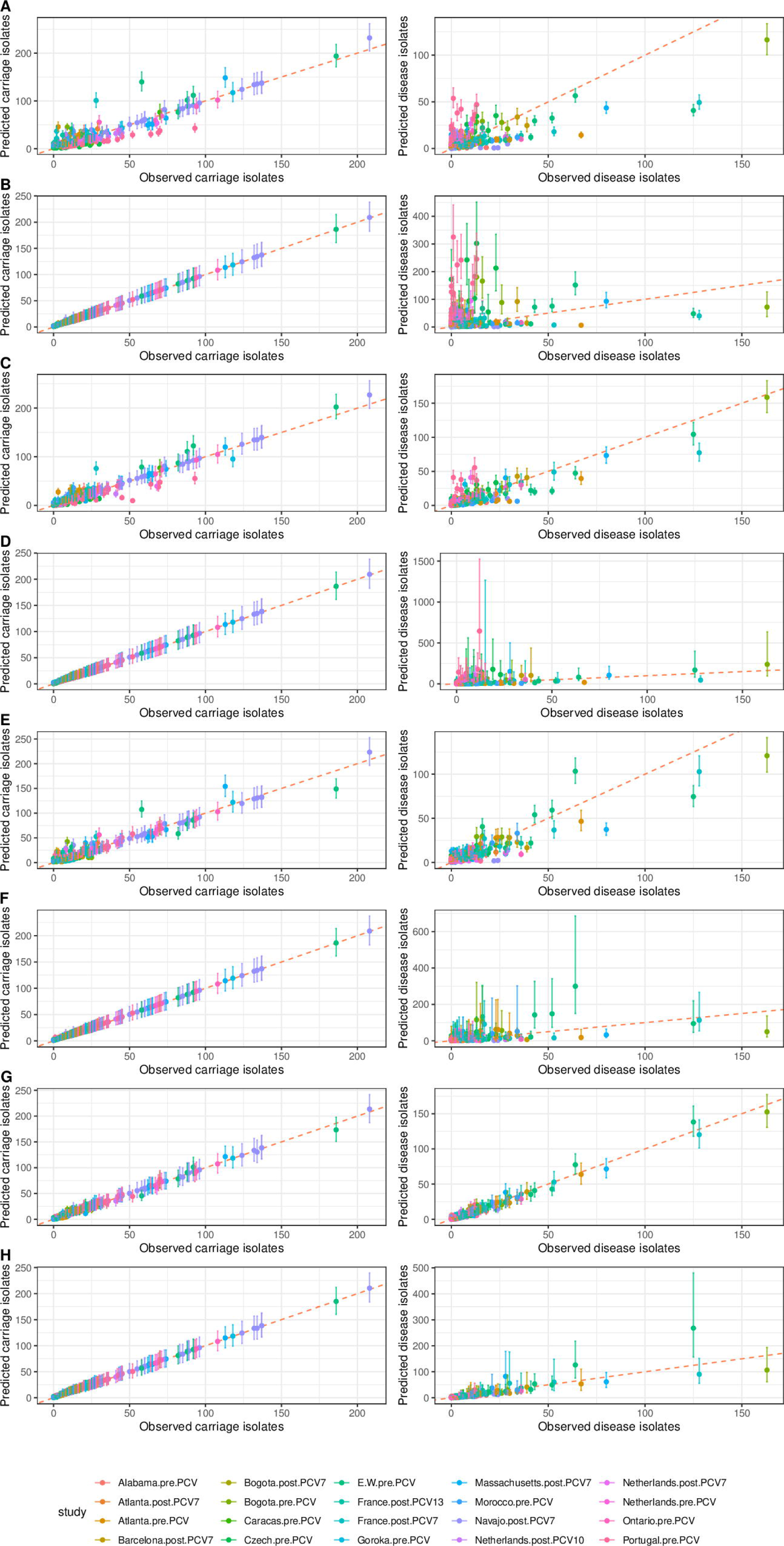
Comparison of observed and predicted counts of each serotype within each study of *S. pneumoniae* invasiveness in children. The points are coloured by the study to which they correspond, and show the observed value on the horizontal axis, and the median predicted value on the vertical axis. The error bars show the 95% credibility intervals. The red dashed line shows the line of identity, corresponding to a perfect match between prediction and observation. The left column shows the correspondence for carriage data (values of *c_i.j_*), and the right column shows the correspondence for disease isolates (values of *d_i.j_*). Each row corresponds to a different model: (A) null Poisson model; (B) null negative binomial model; (C) type-specific Poisson model; (D) type-specific negative binomial model; (E) study-adjusted Poisson model; (F) study-adjusted negative binomial model; (G) study-adjusted type-specific Poisson model; (H) study-adjusted type-specific negative binomial model.

These deviations can be quantified as root mean squared error (RMSE) values (Fig. S7). These show the negative binomial models replicate the carriage values closely, relative to the equivalent Poisson models, with correspondingly greater deviation from the disease counts enabled by the overdispersion permitted by the negative binomial distribution’s precision parameter, 0 That the 0 values were below one for the three simplest negative binomial models suggested there was substantial unmodelled variation (Fig. S8). However, for the study-adjusted type-specific model, the 0 value rose above one, indicating less overdispersion. Correspondingly, the similarly low RMSE values for the Poisson and negative binomial versions of this model suggested adjusting for study and serotype enabled progression rates to be estimated robustly (Fig. S7).

The success of the study-adjusted type-specific models in reproducing the observations may represent overfitting by the most complex models. Therefore formal model comparisons were undertaken using leave-one-out cross-validation (LOO-CV) [62]. Although this suggested the study-adjusted type-specific models were the most appropriate, the large number of model parameters meant the Pareto k statistic diagnostic values were too high for these comparisons to be reliable (Table S2) [79]. This was still true if the models were compared using only the likelihoods calculated from the disease counts, which were more constrained than those calculated from the carriage data (Table S3). Hence models were instead compared using Bayes factors calculated from bridge sampling [63]. This approach was able to accurately assign simulated data to the model under which it was generated (Table S4). The Bayes factors identified the null Poisson model as the most poorly fitting, whereas the study-adjusted type-specific negative binomial model was the most strongly favoured by the data (Table S5).

### Identification of highly invasive non-vaccine serotypes

This study-adjusted type-specific negative binomial model was therefore applied to the full dataset. Both the MCMC traces and is statistics indicated the model fit converged after 25,000 iterations (Fig. S9). This enabled the estimation of invasiveness for 72 serotypes (Fig. 2; Dataset S1), including all 24 included in formulations that are currently licensed or under development [20–22]. The analysis found the serotypes associated with high invasiveness and narrow credibility intervals were the three serotypes added to PCV7 to generate PCV10 (1, 5 and 7F), which are also present in higher-valency PCVs, and serotype 12F, included in PCV20. Other serotypes not included in current vaccine designs, but likely to be highly invasive, include 9L, 19C, 24B, 24F, 25A, 27, 28A and 46. However, the credibility intervals of some of these estimates were substantial, resulting from small overall sample sizes.

**Figure 2.**
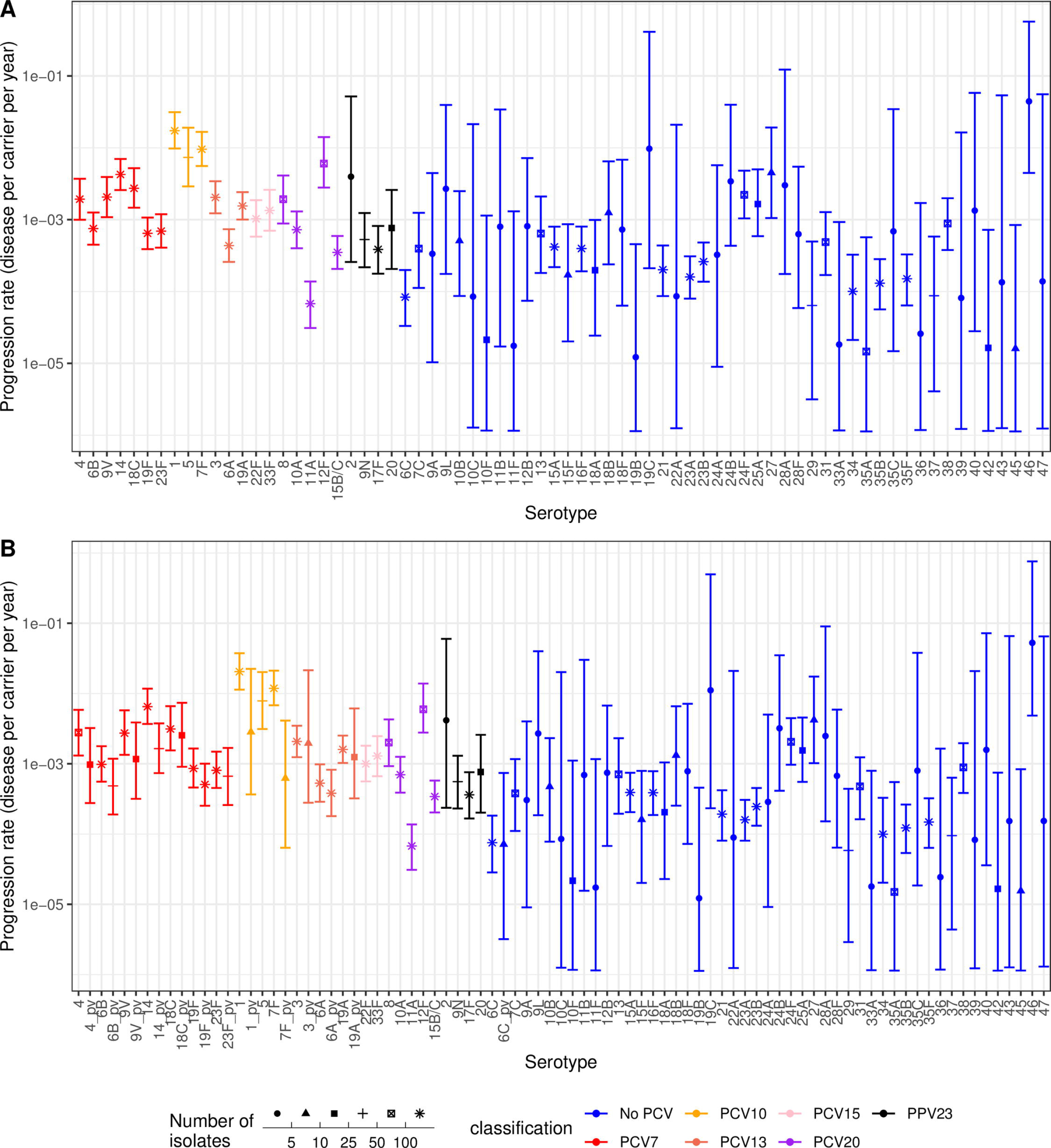
Invasiveness estimates for *S. pneumoniae* serotypes from the study-adjusted type-specific negative binomial model applied to the full dataset of serotypes from child carriage and disease. Points represent the median estimate, and are coloured according to the vaccine formulation in which they are found; all higher valency formulations encompass the serotypes found in lower valency formulations, with the exception of 6A not being present in PPV23. The point shape represents the sample size, summed across disease and carriage isolates from all studies, on which the estimates are based. The error bars show the 95% credibility intervals. (A) Estimates for all 72 serotypes in the full dataset. (B) Estimates when distinguishing between vaccine-type serotypes in unvaccinated and vaccinated populations. Vaccine serotypes in vaccinated populations are denoted with the suffix “_pv”.

The serotypes associated with low invasiveness and narrow credible intervals were 11A (included in PCV20), 6C, 21, 23A, 23B, 34, 35B and 35F. Many other serotypes (e.g. 10F, 11F, 19B, 33A, 35A, 36, 39, 42 and 45) are likely to have low invasiveness, but these estimates have wide credibility intervals. The higher uncertainty generally corresponded with lower sample sizes, although some of these serotypes had a total sample size above 10 (e.g. 10F, 35A and 42), but were nevertheless rare in IPD. The reference population for the model fit was the post-PCV7 sample from the Navajo nation [39], as this study contributed the largest number of isolates (Fig. S1). The model fits found the maximal progression rates were above 10^-2^ IPD cases per carrier per year, with the minimum point estimates below 10^-4^ IPD cases per carrier per year. However, the incidence of IPD in the Navajo population is high, with excellent disease surveillance [80], and therefore other studies were expected to report lower disease cases per carriage episode. Correspondingly, the study-adjustment scale factors were below one for most other datasets (Fig. S10), suggesting many locations would expect to detect around 30-50% of the number of IPD cases per carriage episode for the same carriage serotype composition.

The fitted study-adjustment scale factors were all associated with similarly narrow credibility intervals, demonstrating there were enough shared types between samples to robustly infer these parameters. For four locations (Atlanta, Bogota, the Netherlands and France), both pre-PCV and post-PCV samples were included in the meta-analysis, which might be expected to have similar scale factors, if the host population and disease surveillance were consistent across these eras. However, this was not the case for studies from Bogota. The model’s assumption that serotype invasiveness is constant across studies may be violated by a reduction in vaccine-type invasiveness following PCV introduction, if vaccine-induced immunity inhibited progression from carriage to IPD. Therefore the study-adjusted type-specific negative binomial model was refitted, treating vaccine-type serotypes as different types pre-and post-PCV (including 6A as a PCV7 type, and 6C as a PCV13 type [81]; Fig. S11). The point estimates of invasiveness for most vaccine serotypes dropped following PCV introduction, with the evidence strongest for serotypes 14 and 7F falling after the introduction of PCV7 and PCV10, respectively (Fig. 2). This fit further improved the consistency of scale factors across the Netherlands and Atlanta, but did not resolve the discrepancy between samples from Bogota (Fig. S12). These data suggest PCVs can reduce the invasiveness of vaccine types before they are eliminated by herd immunity, and therefore distinguishing between pre-and post-PCV invasiveness may help standardisation across meta-analyses.

### Comparison of Bayesian modelling with odds ratios

The outputs of the adjusted type-specific negative binomial model were compared to those from estimating pneumococcal invasiveness from the same dataset using odds ratios (Fig. 3). Odds ratios were combined across studies using either random (Fig. 3A) or fixed effects (Fig. 3B) models, and the results disaggregated based on the total number of isolates, summed across carriage and disease, for each serotype. Across all sample sizes, there was a significant positive correlation between the two measures of invasiveness. The range of invasiveness values extended over two orders of magnitude for both methods. Hence the study-adjusted type-specific negative binomial model produced similar relative estimates of invasiveness to odds ratio analyses, while having the advantage of providing both overall, and location-specific, absolute estimates of these progression rates that can be used in quantitative analyses.

**Figure 3.**
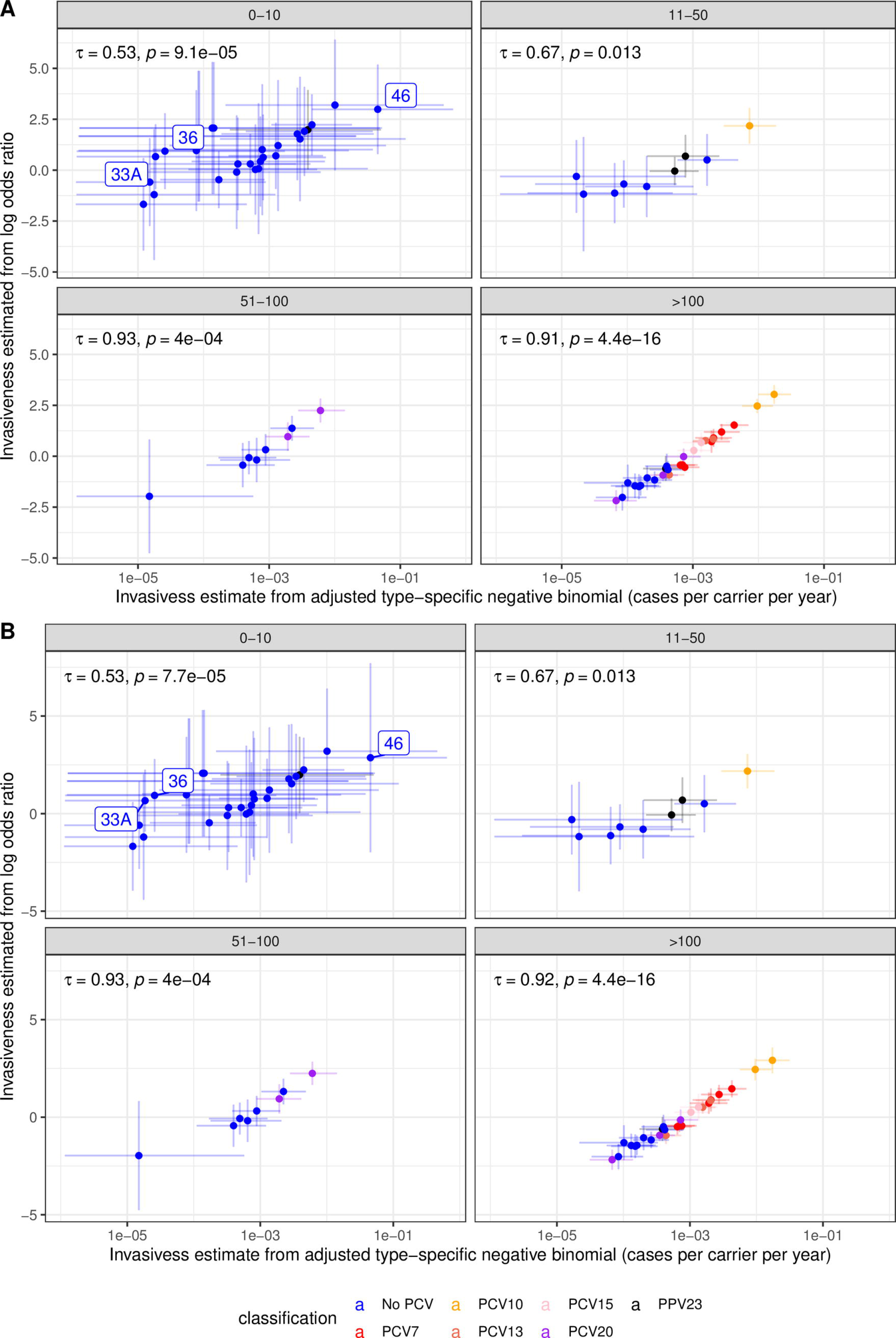
Comparison of the serotype invasiveness estimates from the study-adjusted type-specific negative binomial model with those from meta-analyses of the same dataset calculated as logarithmic odds ratios combined using (A) fixed effects models and (B) random effects models. Points are coloured based on the vaccine formulations in which serotypes are found. The horizontal error bars are the 95% credibility intervals from the Bayesian model, and the vertical error bars show the 95% confidence intervals from the logarithmic odds ratio analyses. Each plot is divided based on the sample size associated with each serotype, calculated as the sum of isolates from carriage and disease across all studies. The correlation between the two estimates is summarised as Kendall’s τ on each panel, with an associated *p* value.

Fixed effects analyses assume there is a single invasiveness value across all locations, whereas random effects analyses allow for variation in the underlying invasiveness estimate between studies. The identification of between-location heterogeneity typically suggests random effects models are more appropriate for analyses of pneumococcal invasiveness [29,36,37]. This variation may either reflect genuine differences in serotype behaviour, or arise from standardisation relative to a variable mix of serotypes between locations. By accounting for this variation, random effects models often calculate wider confidence intervals than fixed effect models (Fig. 3). Additionally, random effects models are typically not recommended where there are fewer than five studies in which a serotype features [82], and therefore meta-analyses of odds ratios may be a mixture of fixed-and random-effects models [37]. By contrast, this single, consistent Bayesian model can be applied across serotypes, regardless of their distribution across studies. Furthermore, this framework explicitly accounts for the uncertainty associated with overdispersed data through the negative binomial precision parameter, 0 (Fig. 1). This variation is quantified separately from the uncertainty in the progression rate estimate. These advantages are in addition to avoiding standardisation using differing mixtures of serotypes, as for odds ratios.

Deviations between the methods in the rank order of invasiveness point estimates were most evident at small sample sizes. This was most notable for serotypes 33A and 36, both of which were associated with relatively high logarithmic odds ratios (above zero) and relatively low Bayesian progression rate estimates (below 10^-4^ cases per carrier per year). Both were observed only in carriage: five isolates across four studies for serotype 33A, and one isolate in each of three studies for serotype 36. Hence the Bayesian model estimates provide a more informative representation of the limited available data.

At the upper end of the scale, the Bayesian model found the most highly invasive serotype to be 46, with credibility intervals indicating this serotype is unlikely to be of intermediate invasiveness [30]. By contrast, the confidence intervals for the odds ratios calculated for serotype 46 were larger relative to the variation between serotypes, with those calculated using random effects spanning the full range of invasiveness point estimates across the species. This serotype was observed in two studies, but never isolated from carriage, and there is further circumstantial evidence that serotype 46 is likely to be highly invasive (see Discussion). Hence this Bayesian framework can use case-carrier studies to provide informative estimates of invasiveness even from small sample sizes.

Odds ratios were previously found to correlate with another measure of invasiveness, the “attack rate”, which in turn negatively correlated with carriage duration [18]. This could result from IPD being most common shortly after the acquisition of a novel serotype in the nasopharynx, which occurs more frequently for serotypes carried for only a short period in each host. The invasiveness values from child IPD data were compared with recent estimates of carriage duration from multi-state modelling of longitudinally-sampled carriage studies in Maela, Thailand [57, 83].

Although the highest invasiveness values were associated with short carriage durations, likely as longer carriage duration is expected to result in higher cross-sectional carriage prevalences, there was no strong overall relationship (Fig. S13).

### Differences in invasiveness between host age demographics

Although *S. pneumoniae* primarily circulate between children, they frequently cause disease in adults. Five datasets were identified in which adult disease cases could be matched with child nasopharyngeal carriage datasets, to estimate serotype invasiveness in adults. Overall, 3,756 carriage isolates and 3,041 disease isolates were extracted across 53 serotypes (Fig. S14). As with the child IPD samples, this dataset was filtered for observations where either the number of carriage or disease isolates was at least five for the model comparison, to improve the power for identifying the best fitting model. This left 3,704 carriage isolates and 2,969 disease isolates across 40 serotypes for this model selection analysis (Fig. S15). As with the child dataset, two 25,000 iteration Hamiltonian MCMCs were used to fit each of the eight models, which converged based on the traces of the posterior probability (Fig. S16) and is values being below 1.05 (Fig. S17). Similar to the analysis of invasiveness in children, comparisons of the observed and predicted isolate counts again showed that only the models adjusting for both *S. pneumoniae* serotype and study were able to accurately reproduce the adult disease case counts (Fig. S18). Bayes factors concurred that the study-adjusted type-specific negative binomial model was the most likely, given the data (Table S6).

Thus, this model was applied to the complete adult disease dataset. MCMC traces and is statistics again indicated the model fit converged after 25,000 iterations (Fig. S19). Like the child analysis, the reference population for the model fit was the post-PCV7 sample from the Navajo nation [39]. The high incidence of IPD in this population is again evident in the adult population, and therefore the other studies had lower study-adjustment scale factors (Fig. S20). Most locations expected to detect 30-50% of the number of IPD cases per carriage episode for the same carriage serotype composition.

This analysis therefore enabled the estimation of invasiveness of 53 serotypes in adults (Fig. 4A; Dataset S2), including the 24 included in PCV formulations currently licensed or under development. The analysis shows the serotypes associated with high invasiveness and narrow credibility intervals not in currently-used PCVs were serotypes 8 and 12F, included in the PCV20 formulation; serotype 20, included in PPV23, and the non-vaccine type serotype 13. Other serotypes not in current vaccine designs, but likely to be highly invasive, included serotypes 9L and 12B. However, the credibility intervals of these estimates were substantial, resulting from small overall sample sizes. The non-PCV serotypes associated with low invasiveness (upper 95% credibility interval estimates <10^-3^ disease cases per carrier per year) were serotypes 6C, 18B, 21, 23A, 23B, 35B and 35F.

**Figure 4.**
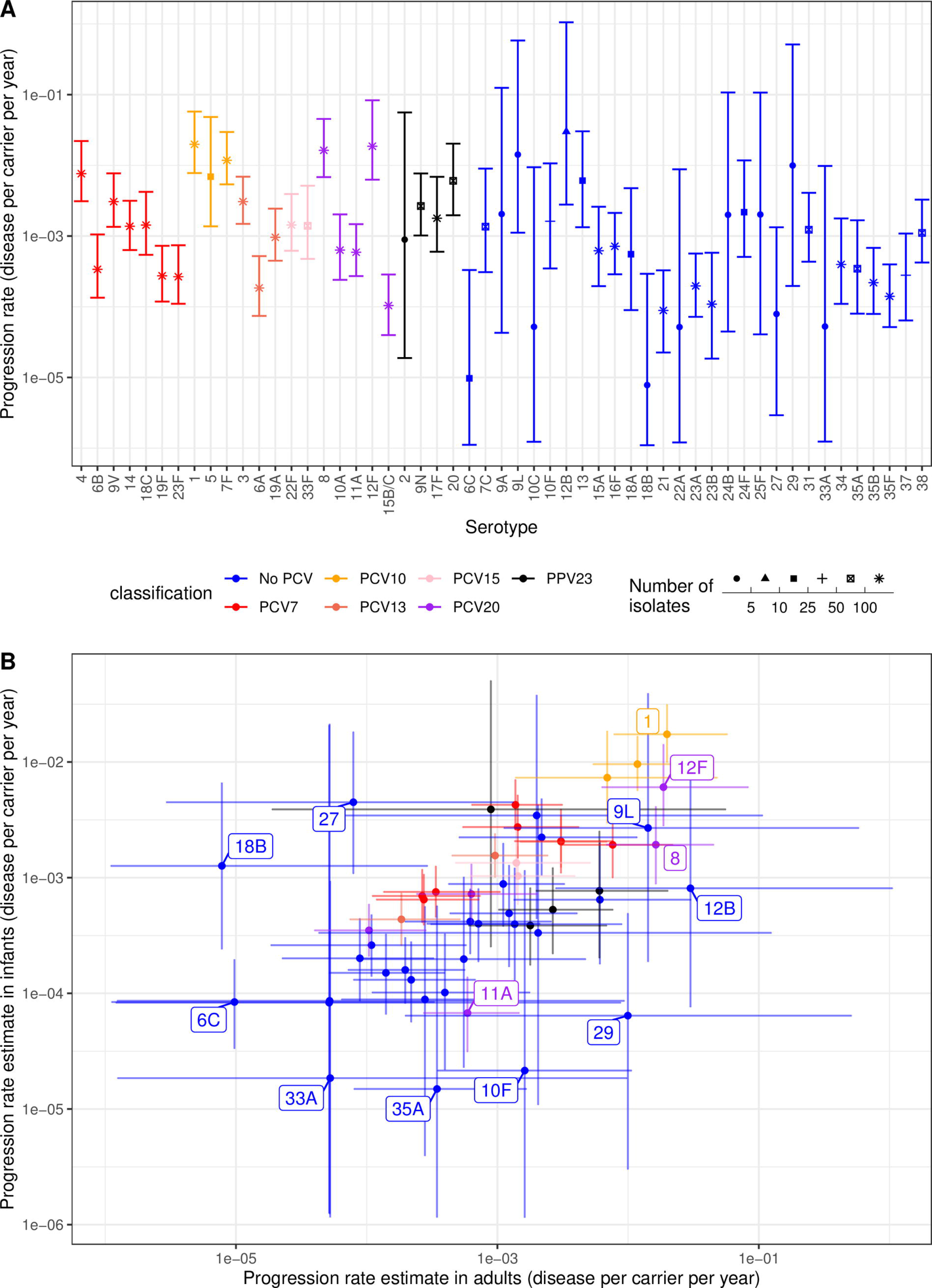
Invasiveness estimates for *S. pneumoniae* serotypes from the study-adjusted type-specific negative binomial model applied to the full dataset of serotypes from child carriage and adult disease. Points represent the median estimates, and are coloured according to the vaccine formulation in which they are found; all higher valency formulations encompass the serotypes found in lower valency formulations, with the exception of 6A not being present in PPV23. The error bars show the 95% credibility intervals. (A) Estimates for all 53 serotypes in the full dataset. The point shape represents the sample size, summed across disease and carriage isolates from all studies, on which the estimates are based. (B) Scatterplot comparing the invasiveness estimates for adults and children.

Comparisons with the corresponding child invasiveness estimates (Fig. 4B) showed many PCV serotypes had similar invasiveness estimates in both children and adults [39], with PCV7 and PCV10 types having mid-range and high invasiveness estimates for both children and adults, respectively. The invasiveness of serotype 6C, which is affected by PCV13-induced immunity against the 6A component, was low in both children and adults. The non-PCV serotype 9L appears to be highly invasive in both populations, although this value was associated with considerable uncertainty. However, other non-PCV types showed a discrepancy between the two age groups. For example, serotypes 12B and 29 were estimated to be highly invasive in adults but not children, though the credible interval ranges were large for both.

Serotypes included in the PCV15 and PCV20 formulations appeared to have mid-range invasiveness in both age groups, except for the rarely invasive serotype 11A, and the consistently highly invasive serotypes 8 and 12F. This suggests the removal of these latter two serotypes through herd immunity from higher-valency PCV infant immunisation programmes would likely be beneficial to the adult population [84, 85]. However, this would concomitantly reduce the effectiveness of the current PPV23 adult vaccine, due to its overlap in serotype coverage with PCVs meaning it would protect against a lower proportion of the post-vaccine *S. pneumoniae* population [29, 86]. PPV23’s residual effect would be determined by the prevalence and adult invasiveness of the non-PCV PPV23 serotypes. Both serotypes 2 and 20 were estimated to be highly invasive for both age groups, albeit with broad credible intervals for adults for the former, due to it only being included in one adult dataset. The remaining non-PCV PPV23 serotypes (9N and 17F) exhibited elevated invasiveness compared to many non-vaccine types, suggesting that PPV23 may offer protection against a limited number of higher-risk serotypes following the implementation of PCV15 or PCV20 infant vaccination programmes.

### Invasiveness varies between pneumococcal strains and serotypes

*S. pneumoniae* populations are genetically diverse, and can be divided into discrete strains [52, 87]. Isolates of a particular serotype are often found across multiple strain backgrounds, and a single strain may be associated with multiple serotypes through switching [53, 88]. To test how each of these characteristics affected pneumococcal progression rates, five different models of isolates’ invasiveness were fitted using the study-adjusted type-specific model framework (see Methods). These corresponded to the hypotheses that an isolate’s invasiveness was determined by its serotype; by its strain background; primarily by its serotype, but modified by strain background; primarily by its strain background, but modified by its serotype; and determined by the combination of its serotype and strain background. All five models were fitted using Poisson and negative binomial distributions.

These ten models were used to conduct a meta-analysis of six studies in which both serotype and strain background could be determined, for at least some isolates (Fig. S21). Three of the studies (post-PCV7 and post-PCV13 South Africa; post-PCV7 USA) were modified versions of a comparison primarily conducted using genomic data [37]. These were combined with pre-PCV genotyped studies from Oxford, UK [31]; Stockholm, Sweden [78], and Finland [74] (see Text S2 and Table S7). For each dataset *i*, isolates were grouped by both their serotype *j* and strain *k*. As with the child serotype-only analysis, models were fitted to a subset of these samples in which either *c_i,j,k_* or *d_i,j,k_* was at least five (Fig. S22). To focus on testing for within-category variation, only strains associated with five different serotypes, and serotypes associated with five different strains, were included in the analysis. The final dataset comprised 11 serotypes, 35 strains, and 46 serotype-strain combinations. All models converged on stable set of parameter estimates, as inferred from the traces of logarithmic posterior probability values (Fig. S23) and distribution of is values (Fig. S24). Very few *c_i,j,k_* or *d_i,j,k_* observations were inconsistent with the 95% credible intervals calculated from any of the fitted models (Fig. S25). There was little evidence of negative binomial distributions improving the fit of these models, and the precision parameters were correspondingly high, albeit with evidence of greater overdispersion when serotype information was omitted from the model fit (Fig. S26). The lowest RMSEs were associated with the models that accounted for both strain background and serotype, either through modification, or when considered in combination (Fig. S27).

Model comparison with LOO-CV found the best-performing models were those that combined information on isolates’ serotype and strain background (Tables S8 and S9), although the Pareto k statistic diagnostic values were again too high for this comparison to be reliable. Comparisons using Bayes factors agreed, identifying the most likely model as that in which invasiveness was primarily determined by serotype, but modified by strain, with *d_i,j,k_* following a Poisson distribution (Table S10). Some carriage sample sizes had to be approximated in this analysis (for the Oxford and Finland studies in particular; see Text S2), but the study adjustment factors were robustly estimated and significantly differed across studies, suggesting the model was able to compensate for this aspect of the data (Fig. S28). To test whether these problems could have affected relative model likelihoods, the model comparison was repeated with a 100-fold change in the carriage sample size values for those studies in which the precise number was not reported. The study-adjustment factor meant the comparisons were relatively insensitive to these changes, and the same model was identified as the most likely, given the data (Table S11). Hence model comparisons demonstrated that neither serotype nor strain background alone determines a genotype’s invasiveness.

### Identification of invasive pneumococcal genotypes

The best-fitting type-determined strain-modified Poisson model converged on a stable solution when applied to the full dataset (Fig. S29). The invasiveness estimates can be plotted to analyse variation by serotype within strains (Fig. 5; Dataset S3). Some strains expressed serotypes of uniformly low invasiveness (serotypes 11A, 15A and 20A within GPSC22), whereas others had consistently higher invasiveness estimates (serotypes 4 and 19A within GPSC27). By contrast, 19A had an elevated invasiveness relative to 19F within GPSC1, and relative to serotype 15B/C in GPSC4. Similarly, serotypes 8 and 33F were substantially more invasive than 11A within GPSC3. Hence there was considerable variation in estimated invasiveness both between, and within, strains.

**Figure 5.**
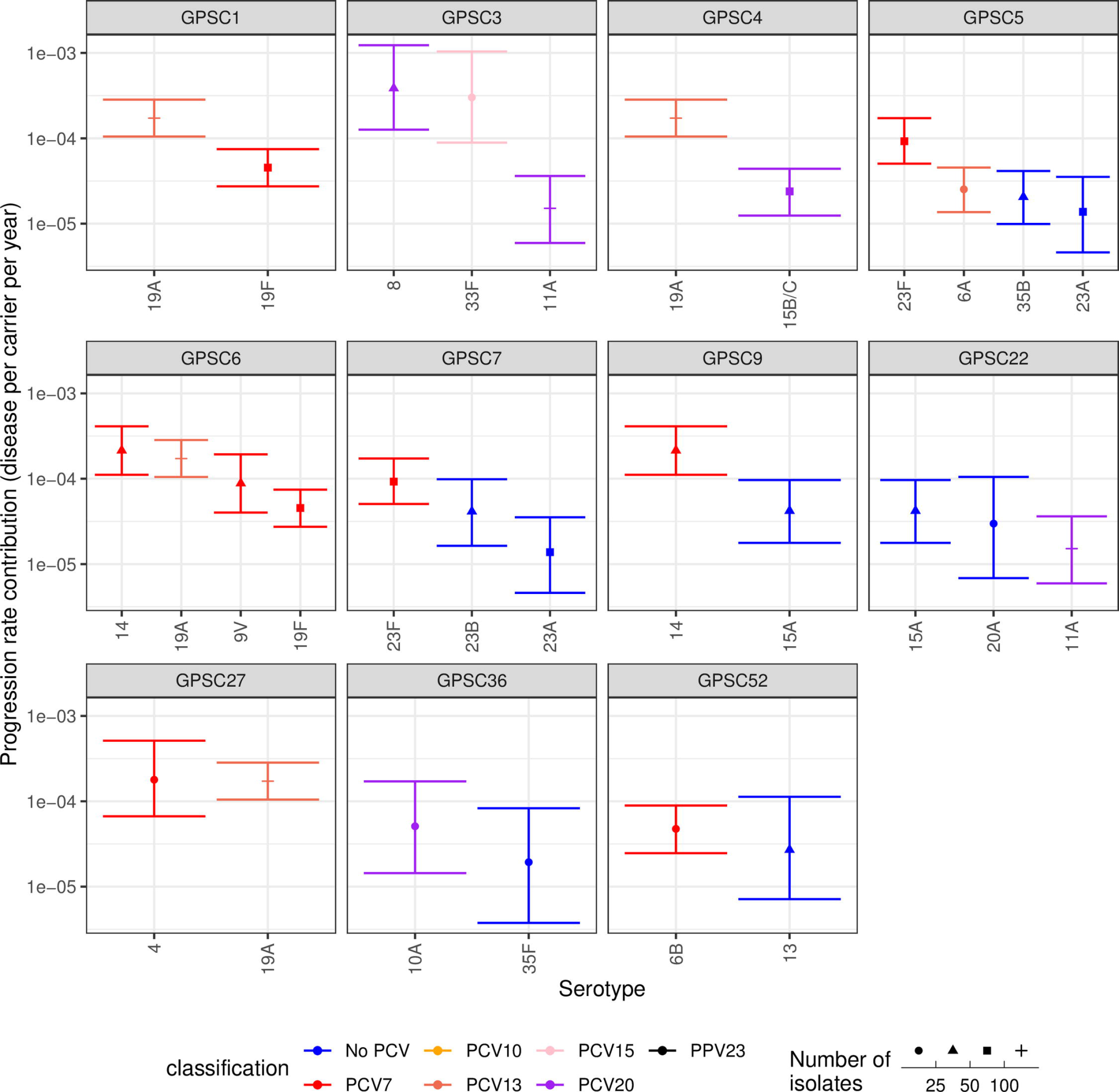
Estimates of the invasiveness associated with serotypes, arranged by the strains in which they are found. Data are only shown for strains found in multiple serotype-strain combinations represented by at least ten isolates across the studies with genotype information. Points represent the median estimate, and are coloured by the vaccine formulations in which the corresponding serotype is present. The error bars represent the 95% credible intervals. The shape of the point represents the sample size on which the estimate is based.

Similarly, the invasiveness contributions of strains were plotted grouped by serotype (Fig. 6). The meta-analysis identified significant evidence to support many of the observations of within-serotype invasiveness variation noted in the individual analyses (Table S12). Heterogeneity was most strongly evident within some of the paediatric types [19], including 6A, 6B, 14, 19F, 19A, 23F and 23A. This may represent these common serotypes being distributed across a wider diversity of strain backgrounds. However, the estimates for other low invasiveness serotypes disseminated across multiple backgrounds, such as 6C, 15B/C and 35B, were more consistent. Furthermore, some highly-invasive serotypes (such as 1 and 12F) were similarly invasive in multiple strains. This implies some serotypes may have a strong effect on an isolate’s invasiveness.

**Figure 6.**
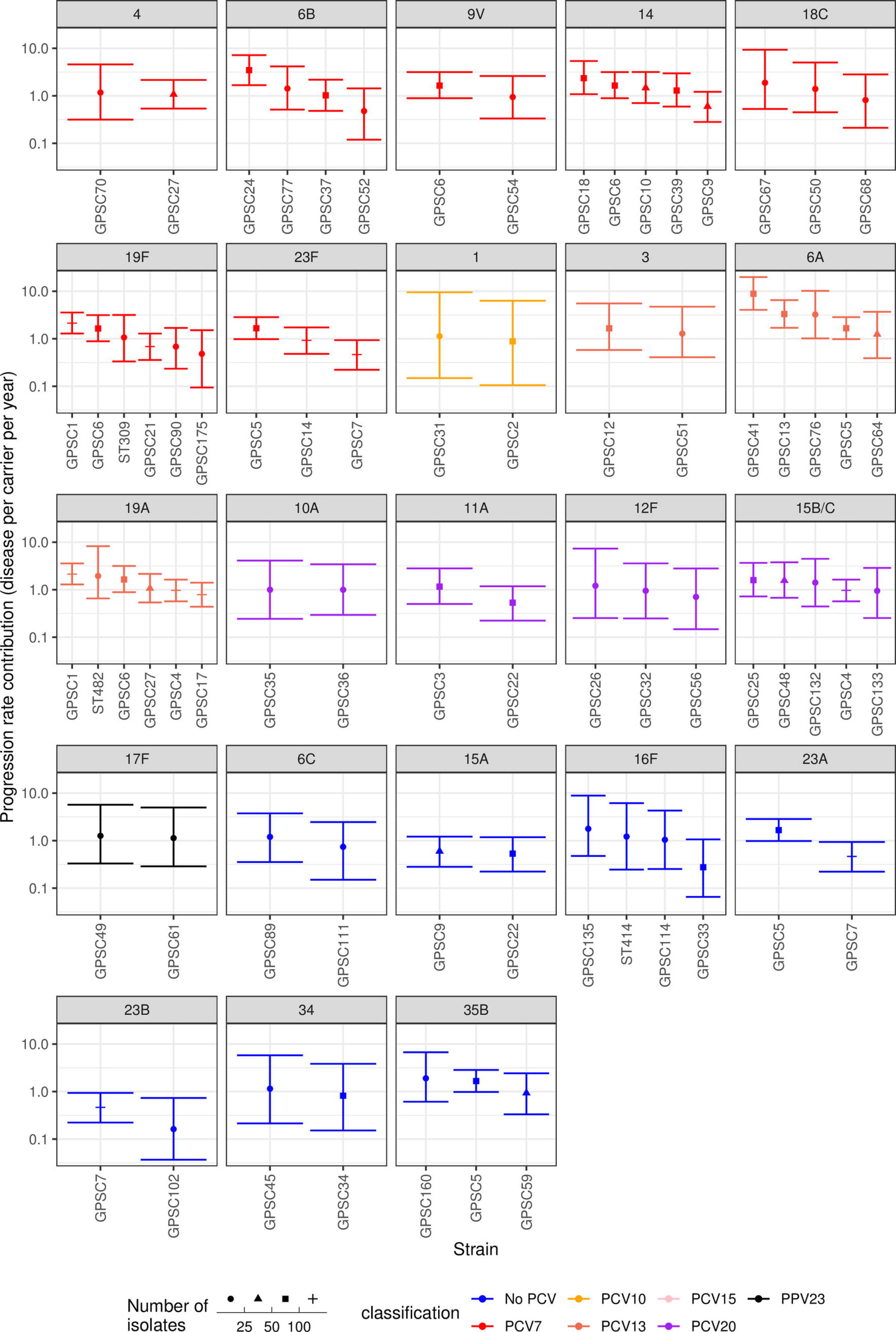
Estimates of the coefficient by which strains modify the invasiveness of their expressed serotype, arranged by the serotypes with which they are associated. Data are only shown for serotypes found in multiple serotype-strain combinations represented by at least ten isolates across the included studies. Points represent the median estimate, and are coloured by the vaccine formulations in which the corresponding serotype is present. The error bars represent the 95% credibililty intervals. The shape of the point represents the sample size on which the estimate is based.

The overall invasiveness estimate was plotted for each serotype-strain combination in the full dataset, including low frequency types, to identify high-risk genotypes that might emerge as causes of IPD post-PCV (Fig. S30). The genotypes identified as most concerning in the near-term were typically those expressing serotypes not targeted by current PCVs and regarded as highly invasive, such as 8 and 12F. Yet there were also examples of genotypes with high invasiveness point estimates expressing serotypes not included in any planned PCVs (e.g. 15A, 18B, 25F, 28A, 33A and 33D), albeit these values were inferred from small sample sizes. Most of these are likely to remain rare, or regress to the population-wide mean as more data emerge, but ongoing surveillance to rapidly identify whether any represent a potentially problematic type post-PCV will be valuable.

An additional genotyped dataset from Portugal (Fig. S31), in which the majority of disease cases were isolated from adults [54], was analysed with the type-determined strain-modified Poisson model (Fig. S32). There were few instances of multiple serotypes being isolated from the same strain, but several serotypes were distributed across multiple genetic backgrounds (Fig. S33). This found evidence of heterogeneity in invasiveness within serotypes 3, 6A and 23F. The elevated invasiveness of serotype 3 was associated with the GPSC6 strain, found to have high invasiveness when expressing multiple serotypes in children (Fig. 6). For serotype 6A, this heterogeneity involved strains that did not appear in the child dataset (Fig. 6), suggesting further within-serotype variation in invasiveness may emerge as studies include a greater diversity of host age groups.

## Discussion

This study describes novel models for using data from cases and carriers to estimate the hazard of an asymptomatically carried pathogen causing a defined disease over a time period, previously defined as a progression rate [2]. This quantity is crucial for determining the efficacy of partial coverage vaccines, such as the PCVs [28, 29], and the flexible framework developed here should be applicable to any diverse microbial population. These models calculate progression rates as absolute quantities, rather than dimensionless ratios, as with previous meta-analyses [29,36,38]. These estimates can enable the translation of alterations in microbial population structures to changes in disease incidence, which is crucial for understanding the consequences of strain- or type-specific interventions [29]. Furthermore, given suitable datasets for multiple pathogens, these models could also be used to estimate changes in disease burdens for alterations across microbiota.

When applied to *S. pneumoniae* serotype data, the best-performing model structures were those in which type-specific progression rates were adjusted by dataset-specific scale factors. In principle, these scale factors should correlate with the burden of disease in a given host population, representing a combination of socio-economic factors and co-morbidities that affect IPD incidence [89, 90]. Yet in practice, they will also represent the probability of an isolate causing IPD being included in a study dataset. This is affected by multiple factors, including the probability of attending a hospital participating in the study, the efficiency of retrieving bacteria from blood cultures, and the consistency with which cultured isolates are collected by research centres [89].

These model structures facilitate comparisons across populations by standardising progression rate estimates between datasets using all common serotypes or strains shared between them. Existing methods for meta-analysis typically standardise data relative to a single type, required to be present in all datasets [36, 38], or combine estimates derived from ratios calculated relative to variable mixes of types in different locations. The dataset-adjusted type-specific progression rate models are better suited to the extensive international variation in *S. pneumoniae* population structures [37, 40], particularly given their ongoing diversification following vaccine introduction [34]. Comparison with odds ratios showed that estimates were correlated at large sample sizes, demonstrating both methods converged on similar conclusions when data were available on many isolates. However, this comes with the caveat that 16 of the 21 studies in the meta-analysis originated from North America or Europe, with most collected prior to PCV introduction. Hence there is a pressing need for paired case and carrier studies from post-PCV settings, particularly from Asia and Africa. As more diverse populations are combined, the improved standardisation across studies from these Bayesian analyses will become more important.

At smaller sample sizes, the Bayesian models were able to produce more informative estimates of invasiveness than odds ratios. Information on types that are rare in individual datasets is particularly important for an opportunistic pathogen as diverse as *S. pneumoniae*, in which serotypes can emerge and expand rapidly post-PCV [25,34,84]. A recent meta-analysis using odds ratios highlighted the conclusion that 12F was the only non-PCV13 serotype to have an invasiveness higher than 19A [36], the serotype responsible for much post-PCV7 IPD [34, 91]. However, this previous study limited its results to 25 serotypes, omitting less common examples that could rise in frequency when serotype replacement is driven by higher-valency vaccines [34, 91]. The Bayesian analysis described in this work identified rare serotypes that may be highly invasive in adults (e.g. 9L, 12B and 29) and children (e.g. 19C, 28A and 46), based on small sample sizes. Serotype 46 has already been observed causing child IPD post-PCV13 within the strain GPSC26, which is more commonly associated with the highly-invasive serotype 12F [33]. These two capsule types are structurally similar [92], which is common for serotypes expressed by isolates of the same strain [53]. Future post-PCV13 surveillance will likely show whether 12F and 46 are truly similarly invasive. As higher-valency PCVs become available, with the potential to remove commonly-carried serotypes such as 11A and 15B/C [20, 93], reducing the uncertainty in invasiveness estimates for remaining non-vaccine types will be of great importance in understanding the vaccine-associated changes in the incidence of IPD.

The need for meta-analysis of multiple datasets is exacerbated by the model comparison in this study that concluded that invasiveness is affected by both strain background and serotype. This conclusion is consistent with the results of some individual studies [37,54,55], although not all [31]. Such a result raises the spectre of separately estimating invasiveness for the hundreds of known *S. pneumoniae* strains, many of which are rare in individual populations [37, 87], in addition to the over 100 known serotypes. Yet the selected model structure suggests serotype is the primary determinant of invasiveness. Consistent with this conclusons, some serotypes (e.g. 1, 12F) exhibited high invasiveness across multiple strain backgrounds, whereas others were consistently low (e.g. 15B/C, 35F).

By contrast, within some paediatric serotypes (6A, 6B, 23F and 23A), there was strong evidence of strains with distinct invasiveness estimates (defined as estimates with non-overlapping 95% credibility intervals). Weaker evidence for within-serotype variation in progression rates (identified through median point estimates for one strain being outside the 95% credibility intervals of another) was observed in four further paediatric serotypes: 14, 19F, 19A and 23B. This may reflect paediatric serotypes having little effect on invasiveness, or their greater prevalence in the pre-PCV pneumococcal population providing greater power to detect variation between strains [32]. Heterogeneity has previously been noted in serotype 14, with both GPSC6 (corresponding to PMEN3, or Spain^9V^-3 [88]) [54, 74] and GPSC18 [31, 37] identified as highly invasive clones (Table S12). In this analysis, both were estimated to be more invasive than GPSC9 within the same serotype. This could result in heterogeneous vaccine impacts between regions, as elimination of serotype 14 by PCVs will likely have a greater benefit across Europe and South America, where it is often associated with the more invasive GPSC6 (https://microreact.org/project/gpsGPSC6), rather than Africa and India, where it is often associated with the less invasive GPSC9 (https://microreact.org/project/gpsGPSC9).

Whether these models are useful long-term, or will ultimately be superseded by alternatives based on genome-wide association studies, will depend on the extent to which progression rates are affected by interactions between genetic loci [94]. It may prove possible to explain a high proportion of the variation in invasiveness between microbes through a tractable number of polymorphic loci that each independently contribute to a microbe’s invasiveness using genome-wide association studies [47–50]. If so, then the individual estimates for strains can be replaced with a simpler genome-based approach, which would be capable of predicting the invasiveness of previously unseen genotypes. However, if invasiveness depends on combinations of loci [94], it may only be possible to estimate progression rates for extant common types from epidemiological studies. Other alternative model structures include random effects models and multivariate regression analyses, which can incorporate information on the host population characteristics [2,39,95]. However, these models typically require comprehensive surveillance of disease and detailed information on host populations, which limits the range of studies that can be combined in meta-analyses of diverse bacterial populations.

Even using more conventional approaches to estimating invasiveness, whole genome sequencing clearly represents an important tool in future case-carrier studies of opportunistic pathogen invasiveness. In addition to assigning isolates to strains, genomic data is a reliable and cost-effective means of classifying microbes according to their capsular structures [96, 97]. Furthermore, it also enables the extraction of additional clinically-relevant information, such as antibiotic resistance loci [88, 98], that may be incorporated into future extensions of these models.

However, the use of genomic data exacerbates an underlying problem with these models, as high-quality genome assemblies are not necessarily generated from all isolates recovered from carriage. The model assumes there are no false negative swabs in carriage studies, meaning there is an expectation that all carriage samples will be present in the genomic data. Hence the omission of genomic data adds to a false negative rate that, even with serotype data, already reflects the imperfect detection of colonisation by nasopharyngeal swabbing. This can be the consequence of all resident bacteria being missed, or the difficulty of detecting lower prevalence types in instances of multiple carriage [99]. Unless there is variation in the ability to detect carriage of different types, the underestimation of carriage means all progression rates will be uniformly overestimated. However, if only a subset of datasets in a meta-analysis has a lower sensitivity for detecting carriage, the model will adjust for this by increasing the associated scale factors. Hence the model structure is sufficiently flexible to account for such differences. This is necessary, as even with standardised protocols for detecting colonisation, the population-wide estimates for carriage rates for a location will depend on the exact demographics sampled.

Some of these problems with detecting colonisation can be addressed using new techniques with improved sensitivity for detecting multiple serotypes in carriage [100]. However, to simultaneously obtain the genotyping information needed to precisely characterise an isolate’s invasiveness using such mixed samples requires deep genome sequencing, or similar high-sensitivity molecular genotyping techniques. Such data will not only improve the input to epidemiological models, such as those outlined here, but also potential future models based on combinations of genetic loci. Yet meta-analyses of case-carrier studies can also be improved by simple changes to reporting, without any alterations to current procedures. The data and models described in this work are made available for re-use, modification, and application to other multi-strain microbes (see Methods). The fitting of the model to new datasets will require studies to report their raw data in a standard format, and include the total number of swabs included in the carriage study, as well as an informed estimate of the size of the population under surveillance for IPD. To improve comparability between future studies, it would also be ideal for studies to stratify these data by age category, to enable specific subsets to be employed in different meta-analyses studying specific demographics. Therefore, as the repertoire of vaccines against diverse pathogens expands, we can hope to monitor and understand their impact on population-wide microbial invasiveness using improved models, advances in sequencing technology, and greater transparency of reporting.

## Acknowledgements

We thank Raquel Sá-Leão, Mário Ramirez, Bill Hanage and Birgitta Henriques-Normark for providing further information on case-carrier datasets, and Daniel Weinberger for helpful discussions on model structure and fitting. AL was funded by an investigator-initiated grant from GSK to NJC. NJC was supported by the UK Medical Research Council and Department for International Development (grants MR/R015600/1 and MR/T016434/1) and a Sir Henry Dale Fellowship, jointly funded by Wellcome and the Royal Society (grant no. 104169/Z/14/A). The funders had no role in study design, data collection and analysis, decision to publish, or preparation of the manuscript.

## Data Availability

All data and code used for the described analyses are available in a GitHub repository at https://github.com/nickjcroucher/progressionEstimation/. This repository has been assigned the DOI 10.5281/zenodo.5762037.

