## Supplementary text, figures and tables for "Analysing pneumococcal invasiveness using Bayesian models of pathogen progression rates"

**Text S1-2**

**Figures S1-S33**

**Tables S1-12**

**Datasets S1-S3** (legends only)

**Text S1: Validation of model fits using simulated data**

Simulated data were generated for 20 types of a microbe, each sampled across 20 studies. These were used to test the eight models that could be applied to typed microbial populations: the null Poisson model, the null negative binomial model, the type-specific Poisson model, the type-specific negative binomial model, the study-adjusted Poisson model, the study-adjusted negative binomial model, the study-adjusted type-specific Poisson model, and the study-adjusted type-specific negative model.

The carriage frequency of each type *j* in each study *i*, $\rho_{i,j}$, was randomly generated from the distribution:

$$\rho_{i,j} \sim U(0,0.05)$$

This ensured the sum of $\rho_{i,j}$ within a study remained below one. The progression rates were either set to a single fixed value of $v$ across all *j* (null and study-adjusted models), else the value of $v_j$ was fixed across studies as a random value generated from the distribution:

$$log_{10}(\nu_j) \sim U(-5,-1)$$

This was truncated relative to the full range of values allowed by the model fit, to ensure the estimation of 95% credibility intervals were unlikely to be affected by the prior distribution boundaries.

Across studies, the number of carriage samples, $\eta_i$, was drawn from a Normal distribution, rounded to the nearest integer:

$$\eta_i \sim [N(1000,250)]$$

The sizes of the populations under surveillance for disease $N_i$ was set to be more variable, as observed in the actual data (Table S1):

$$log_{10}(N_i) \sim U(4,6)$$

For the study-adjusted models, the study scaling parameter $\gamma_i$ was set to one for $i$=1. For the other studies, $\gamma_i$ was randomly drawn from the distribution:

$$log_{10}(\gamma_i) \sim N(0,2)$$

For all models using a negative binomial distribution, the precision parameter ϕ was set to 0.1.

For all eight tested models, counts of carriage isolates of each type in each study ($c_{i,j}$) were generated from the distribution:

$$c_{i,j} \sim Binom(\rho_{i,j}, \eta_i)$$

For each tested model, the count of disease isolates of each type in each study ($d_{i,j}$) were generated from the Poisson or negative binomial distribution, using the appropriate calculation for $\mathbb{E}[d_{i,j}]$.

All eight models were fitted to data simulated from each model, using two MCMCs of $10^4$ iterations. Bayes factors were used to compare the model fits to each dataset using bridge sampling run for $10^4$ iterations.

**Text S2: Sample descriptions for genomic and genotyped datasets**

The *S. pneumoniae* population was previously divided into strains by the Global Pneumococcal Sequencing project [33] using PopPUNK [5]. Previous estimation of the effect of strain background on invasiveness by Gladstone *et al* [37] used two study populations. The first combined two carriage studies from Agincourt and Soweto, in South Africa, with nationwide surveillance of IPD by the National Institute for Communicable Diseases. After excluding carriage and disease isolates from individuals reported as being HIV positive, genomes were available for 1,089 carriage isolates and 855 IPD isolates from children under seven years old. Model fitting required calculating the number of swabs taken from HIV negative children under seven in the carriage studies. In Soweto, the reported age stratification meant this number could only be calculated for children under five [101], and therefore four carriage isolates from children aged five or six were excluded from the study. In Agincourt, the reported age stratification meant 33 isolates from children aged six had to be excluded [102]. As the prevalence of HIV was low in infants in the Agincourt study [103], and the exact number of swabs from HIV positive children was not reported, no adjustment was made to the sampling based on HIV status. The population of children under surveillance for IPD was estimated from nationwide census data between 2009-2013 [104]. All samples were stratified into the post-PCV7 (2009-2011) and post-PCV13 (2012-2013) periods [33].

The second study population combined a carriage survey from Massachusetts, USA and isolates from the Active Bacterial Core Surveillance (ABCS) of IPD across several states. Isolates were collected over the sampling periods of 2000-2001, 2003-2004, 2006-2007 and 2008-2009. As only 51 IPD samples were available for the earliest two periods, and the population was changing rapidly over these years [93], only samples from the more stable post-PCV7 2006-2009 period were studied [105]. This included 405 IPD isolates, modelled as coming from a time interval of two years (as samples only came from 2007 and 2009) from a population size calculated from the ABCS population matrix [106]. Of the carriage isolates, 280 were assigned to strains from genomic data, while 291 were only associated with multi-locus sequence typing (MLST) genotypes [107]. Gladstone *et al* assigned these isolates to strains using the close relationship between MLST and PopPUNK population classifications [5,37].

As these studies were estimating odds ratios across samples stratified by year, Gladstone *et al* randomly subsampled isolates to ensure an even ratio of disease and carriage at each sampling timepoint [37]. This correction is unnecessary with the models described in this study, as the $\gamma_i$ scaling parameters can adjust for differing levels of surveillance between samples (incorporating variation in both the samples collected, and the subset sequenced). Furthermore, such subsampling would distort the inferred $\rho_{i,x}$.

MLST genotypes were also available for isolates from the studies of serotype invasiveness from Stockholm [78], Finland [74,108] and Oxford [31]. Where possible, these were assigned to GPS strains using the isolate's MLST data, or the clonal complex to which they belonged at the time of publication [4]. If an isolate could not be assigned to a GPS strain, then its MLST sequence type was used to designate its genotype instead. Isolates that could not be unequivocally assigned a strain, serotype and source (carriage or disease) were omitted.

For the Oxford study, these criteria meant 84 of the 150 disease isolates, and 113 of the 351 carriage isolates, could be analysed. The carriage isolates were assembled from three different surveys, only one of which could be linked to a more detailed published description of the total carriage sample size [109]. As the isolates in this study were recovered from 213 individuals, and the Oxford study attempted to minimise multiple sampling from any one host [31], the overall carriage sample size for the three studies was approximated as three times the number of individuals in this published study (639).

For the Finland study, the criteria described above meant 143 of the 224 disease isolates, and 102 of the 217 carriage isolates, could be analysed. The carriage isolates were recovered from ten longitudinal samples from each of 329 individuals [35]. As the Finland study sought to avoid multiple sampling of the same carriage episode [74], the host population size was used as the carriage sample size.

For the Stockholm study, the criteria described above meant 65 of the 165 disease isolates, and 178 of the 550 carriage isolates, could be analysed. The carriage isolates were recovered from studies conducted between 1997 and 2004. Two published descriptions of the original carriage studies reported the recovery of 506 isolates from 1,330 individuals [110,111]. As this accounted for over 90% of the nasopharyngeal isolates, this number was used as the carriage sample size.

An additional study from Portugal used a collection of disease isolates primarily recovered from adults [54]. Applying the same criteria as to the child disease studies allowed unambiguous assignation of genotype and serotype to 463 of the 769 carriage isolates, and 152 of the 475 of the disease isolates. The carriage isolates were a subset of a previous study [112], and originated from 1,170 nasopharyngeal swabs (R. Sá-Leão, personal communication).

### Supplementary Figures

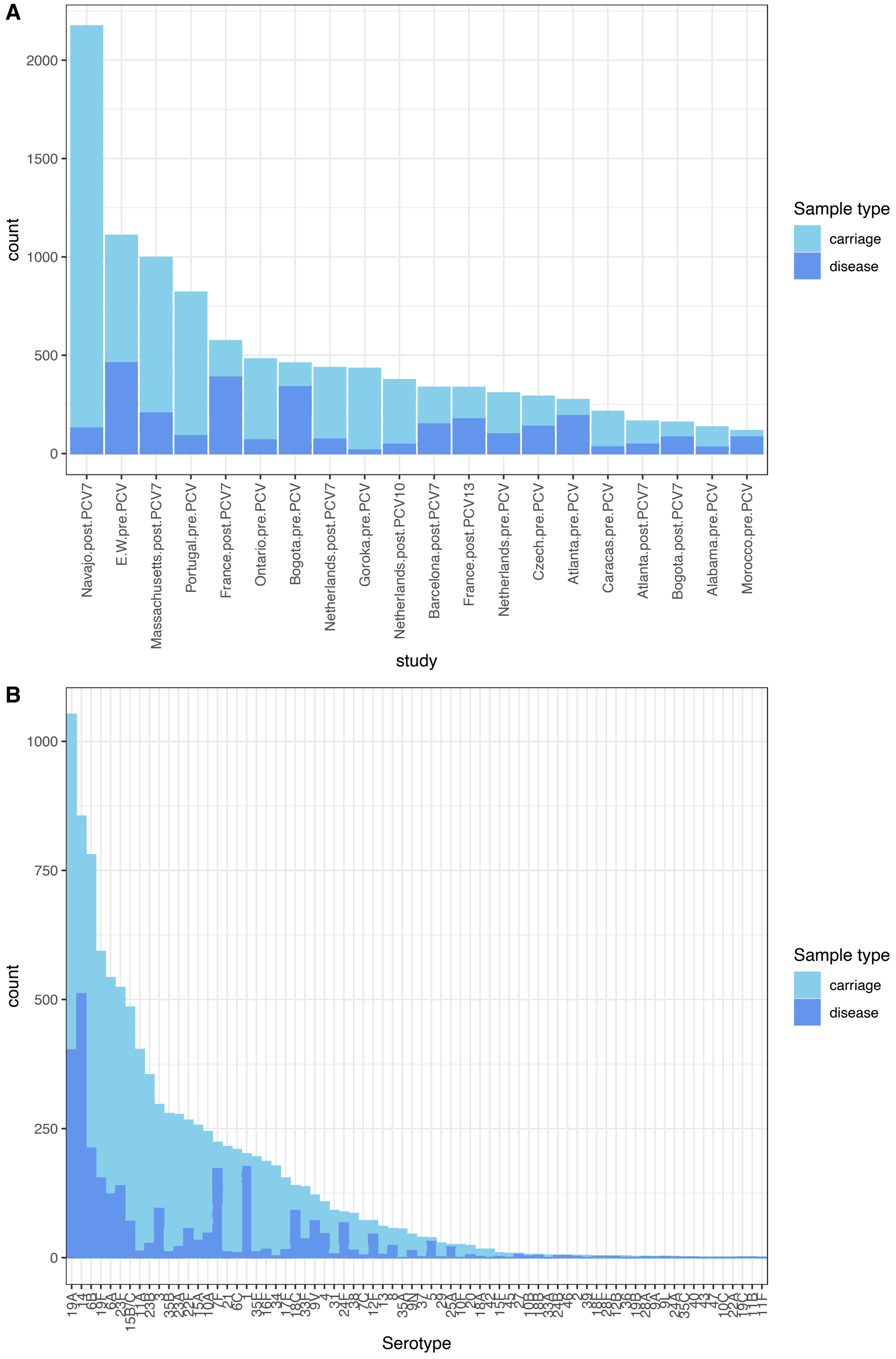


**Figure S1** Properties of the child serotype dataset. (A) Stacked bar plot showing the distribution of carriage and disease isolates between studies. (B) Stacked bar plot showing the distribution of carriage and disease isolates between serotypes.

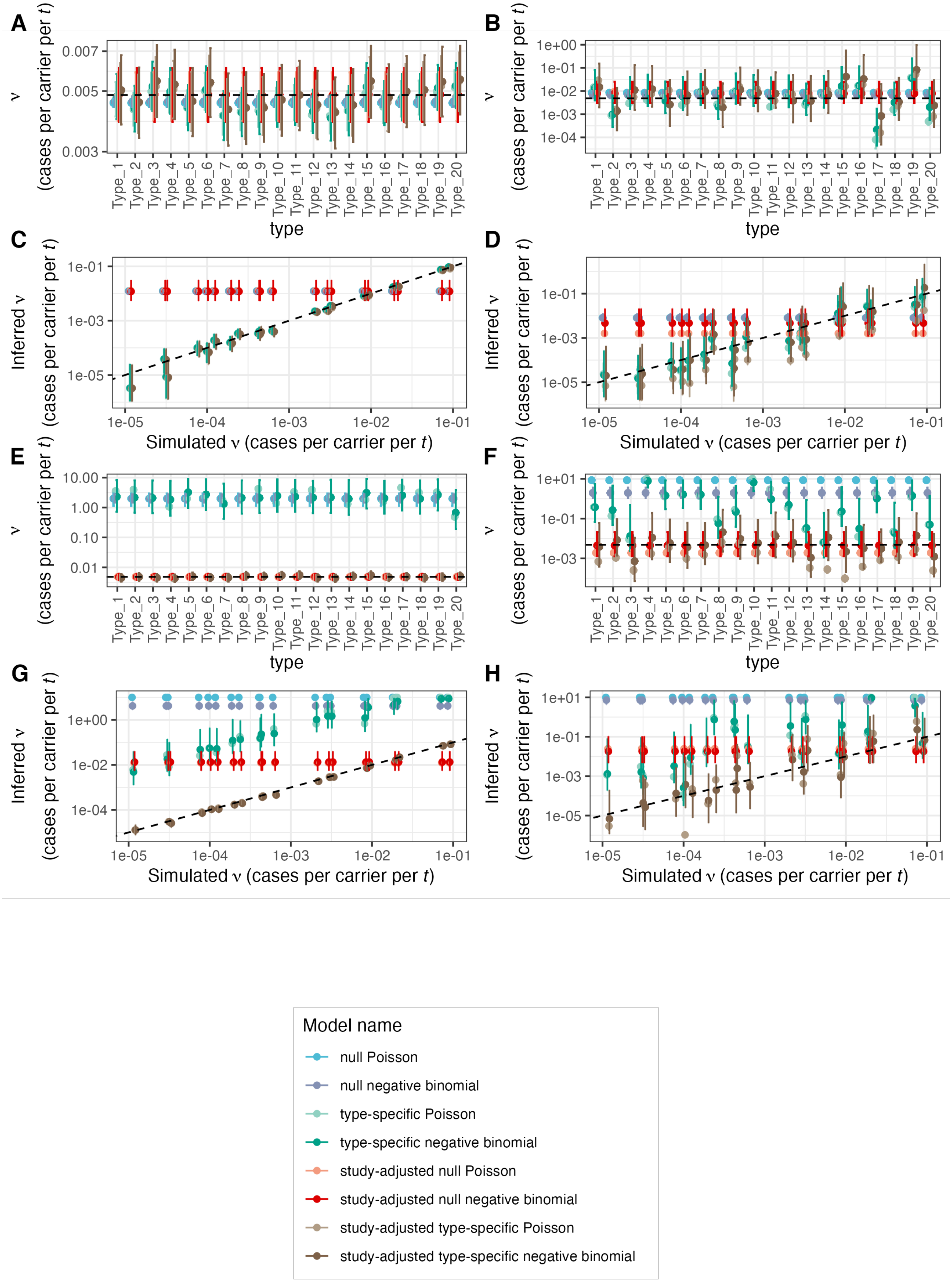

A
B
C
D
E
F
G
H
ν (cases per carrier per t)
Inferred ν (cases per carrier per t)
type
Simulated ν (cases per carrier per t)
Type_1
Type_2
Type_3
Type_4
Type_5
Type_6
Type_7
Type_8
Type_9
Type_10
Type_11
Type_12
Type_13
Type_14
Type_15
Type_16
Type_17
Type_18
Type_19
Type_20
0.003
0.005
0.007
1e+00
1e-01
1e-02
1e-03
1e-04
1e-05
10.00
1.00
0.10
0.01
1e+01
Model name
null Poisson
null negative binomial
type-specific Poisson
type-specific negative binomial
study-adjusted null Poisson
study-adjusted null negative binomial
study-adjusted type-specific Poisson
study-adjusted type-specific negative binomial

**Figure S2** Plots of estimated progression rate values from model fits to simulated data. Each panel shows the fit of eight different models, indicated by colour, to data simulated from a single model structure. Points represent the median estimate, and the error bars show the 95% credible intervals. The models from which data were simulated were (A) null Poisson, (B) null negative binomial, (C) type-specific Poisson, (D) type-specific negative binomial, (E) study-adjusted Poisson, (F) study-adjusted negative binomial, (G) study-adjusted type-specific Poisson, and (H) study-adjusted type-specific negative. For panels (A), (B), (E) and (F), all types had the same progression rate, and therefore the dashed horizontal line represents the single true value used in the simulations. For panels (B), (C), (G) and (H), each of the 20 types had a different progression rate, which determines the horizontal position of the point on the graph. In these four plots, the dashed line indicates the line of identity.

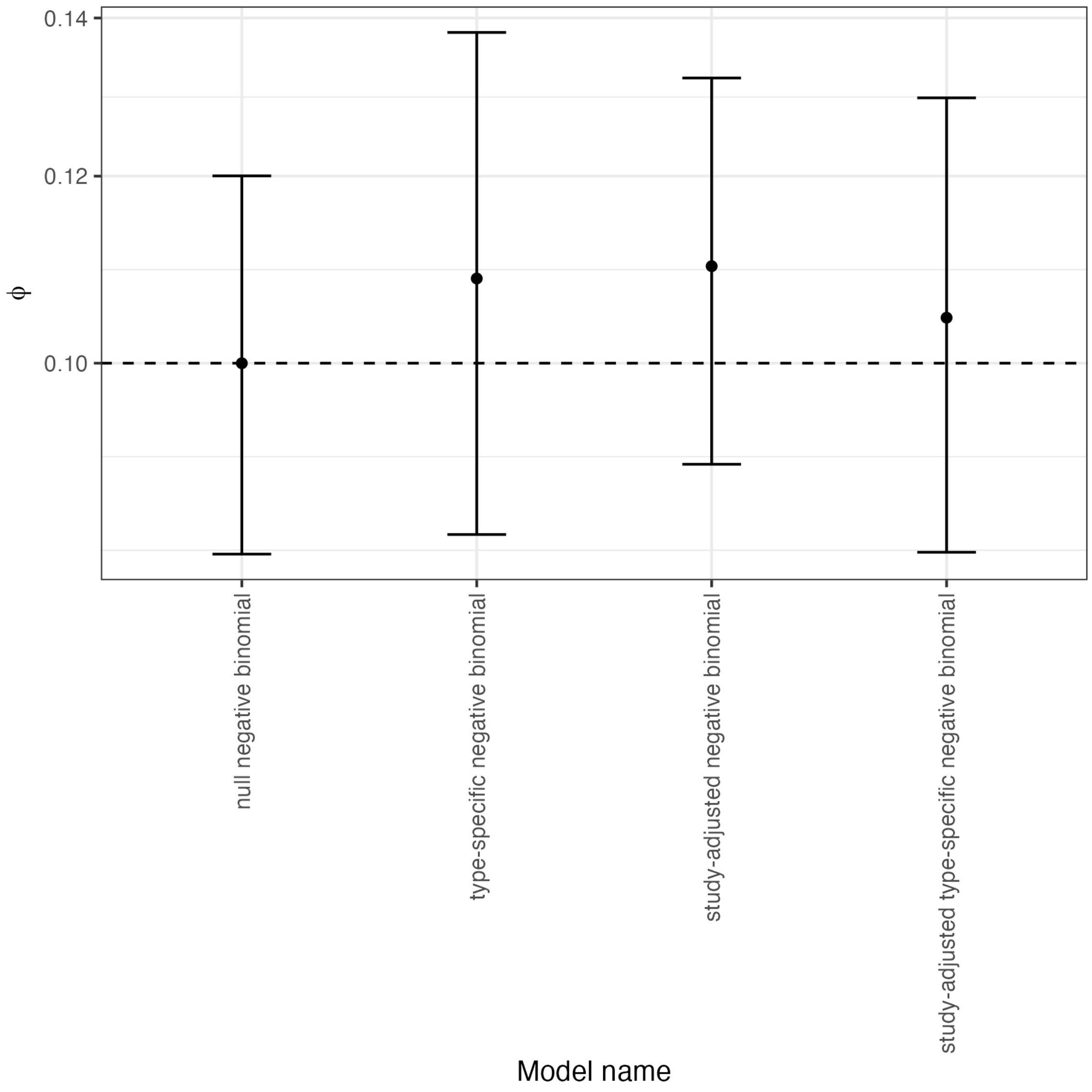


**Figure S3** Plot of the estimated values of the precision parameter ϕ from model fits to simulated data. The horizontal position of the point describes both the model from which data were generated, and the model that was fitted to the data. The vertical position of the point indicates the median estimate of the parameter, and the error bars represent the 95% credibility interval. The horizontal dashed line indicates the true value of the parameter used to simulate data, $\phi = 0.1$.

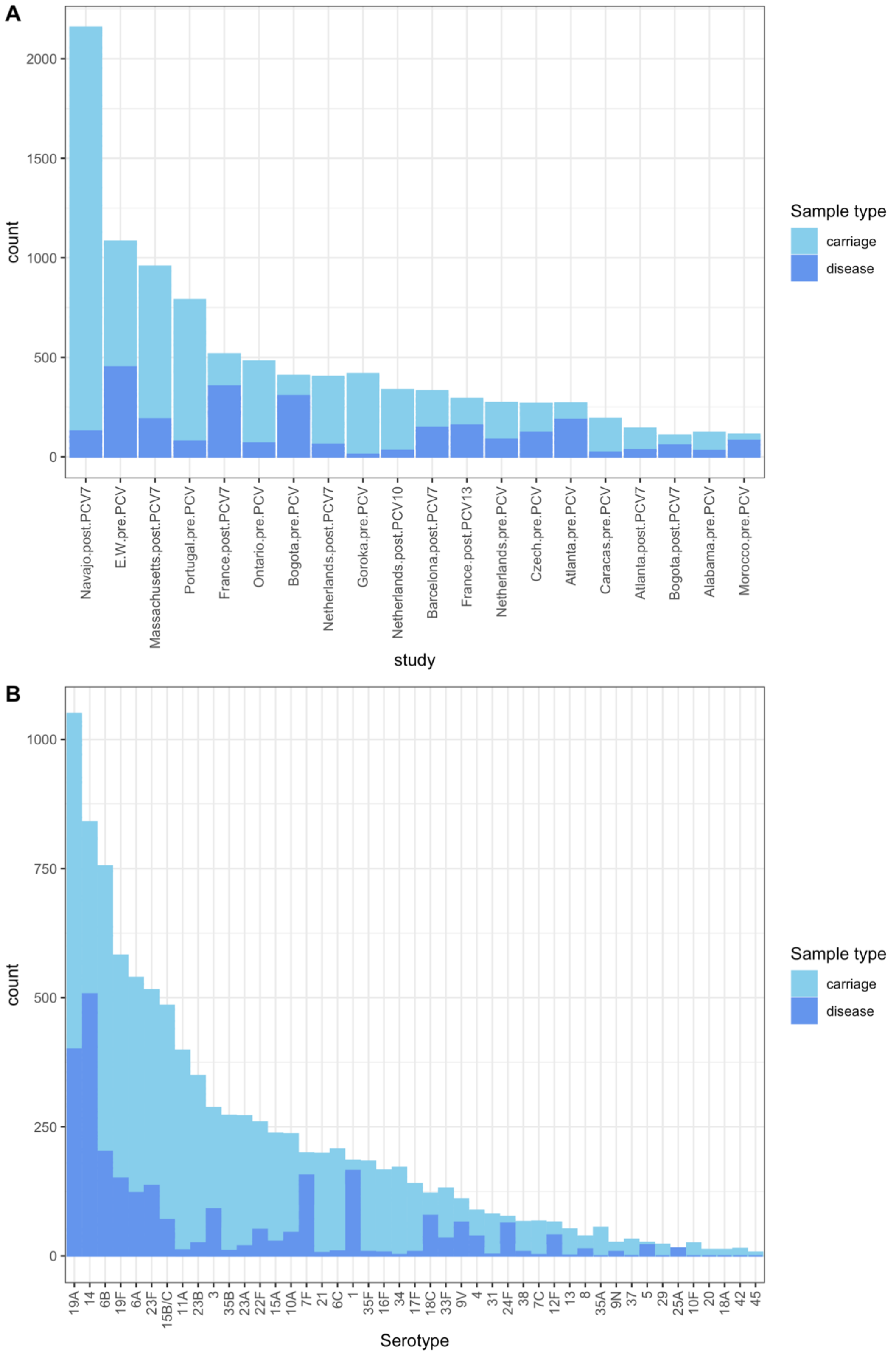


**Figure S4** Properties of the child serotype dataset after removing datapoints in which fewer than five isolates of a serotype were recorded from both carriage and disease in a given study. (A) Stacked bar plot showing the distribution of carriage and disease isolates between studies. (B) Stacked bar plot showing the distribution of carriage and disease isolates between serotypes.

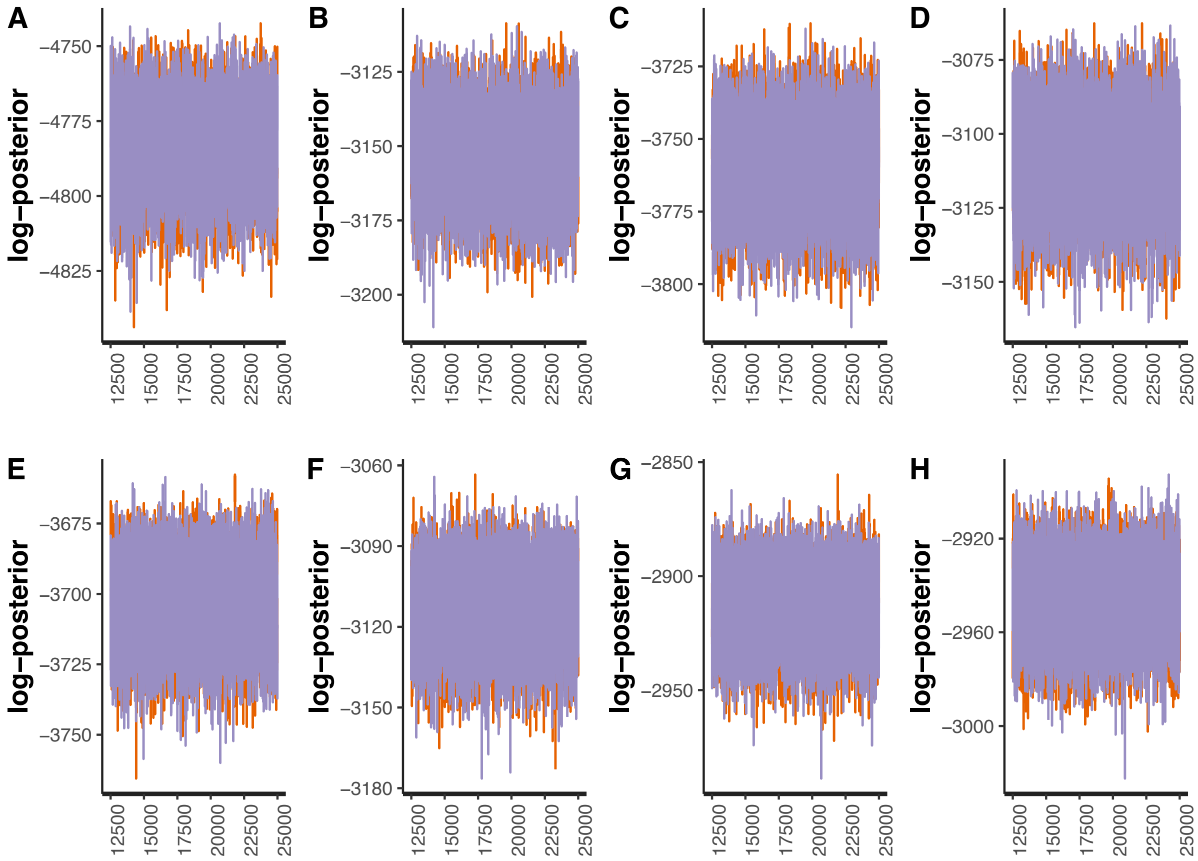


**Figure S5** Line plots showing the post-warmup MCMC traces for the logarithmic posterior probabilities across two independent chains for models fitted to the filtered serotype data from child carriage and disease. The horizontal axis shows the generation of the MCMC, with values for the two chains shown by orange and purple lines. Each panel corresponds to a different model: (A) null Poisson model; (B) null negative binomial model; (C) type-specific Poisson model; (D) type-specific negative binomial model; (E) study-adjusted Poisson model; (F) study-adjusted negative binomial model; (G) study-adjusted type-specific Poisson model; (H) study-adjusted type-specific negative binomial model.

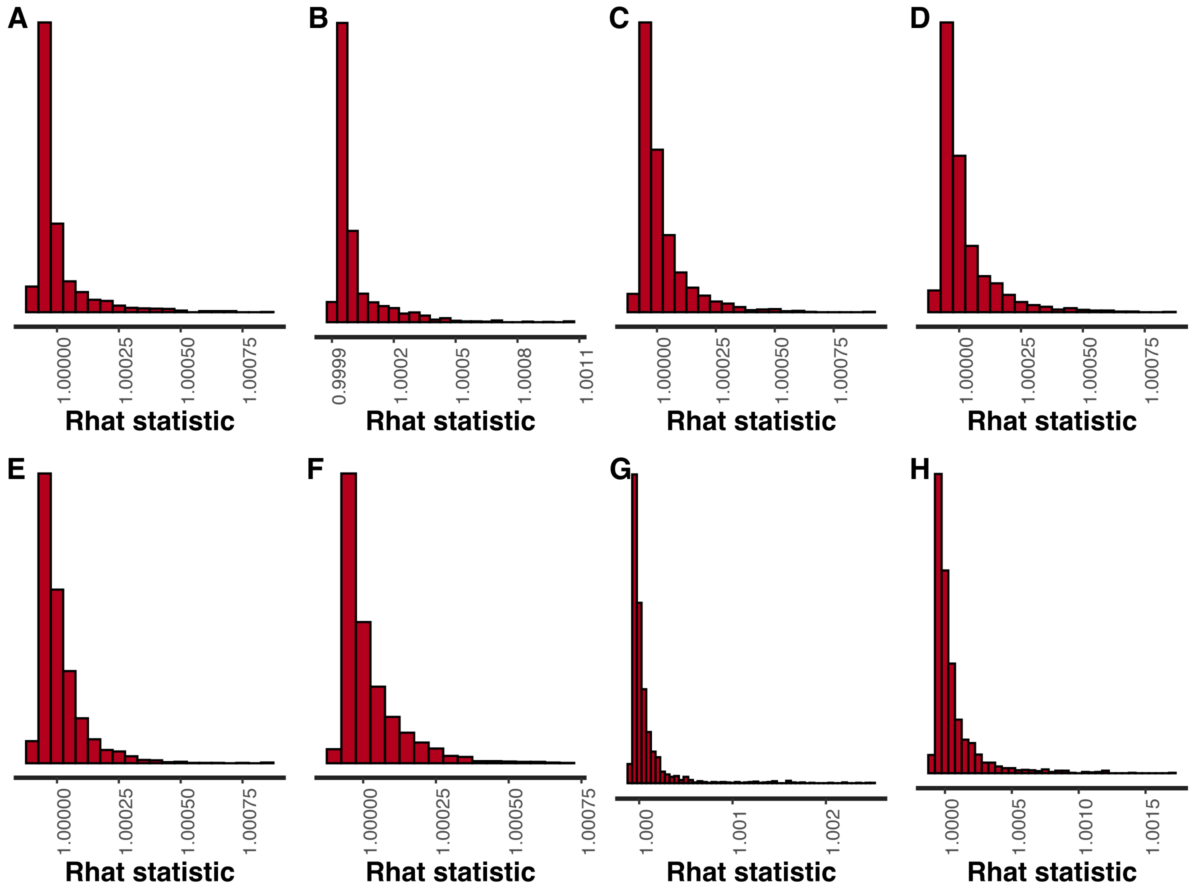


**Figure S6** Histograms of Rhat values calculated from the paired MCMCs for models fitted to the filtered serotype data from child carriage and disease. Each panel corresponds to a different model: (A) null Poisson model; (B) null negative binomial model; (C) type-specific Poisson model; (D) type-specific negative binomial model; (E) study-adjusted Poisson model; (F) study-adjusted negative binomial model; (G) study-adjusted type-specific Poisson model; (H) study-adjusted type-specific negative binomial model.

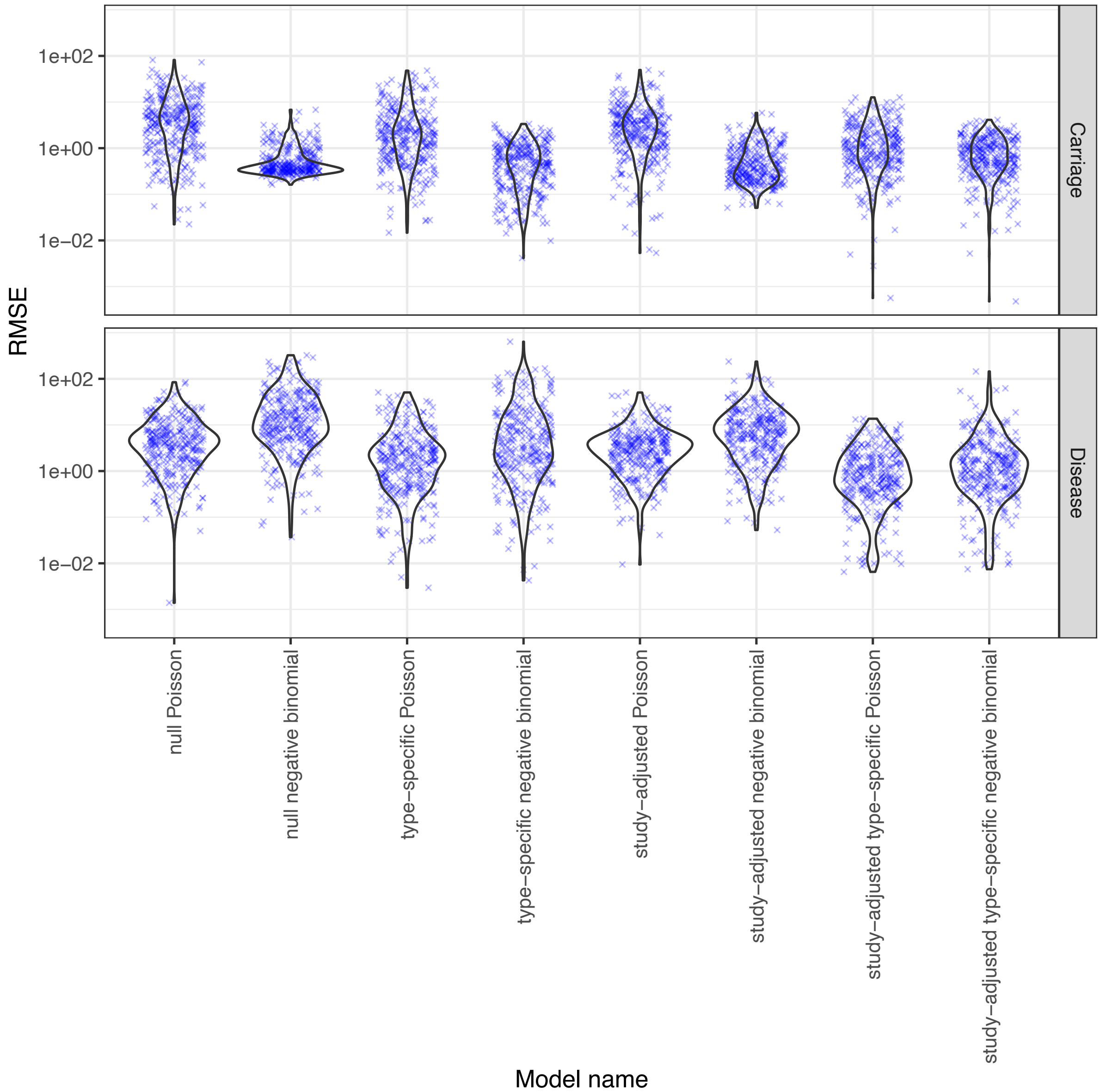


**Figure S7** Violin plots showing the root mean square error between observed and predicted values across model fits to the filtered serotype data from child carriage and disease. Blue crosses represent the individual observations.

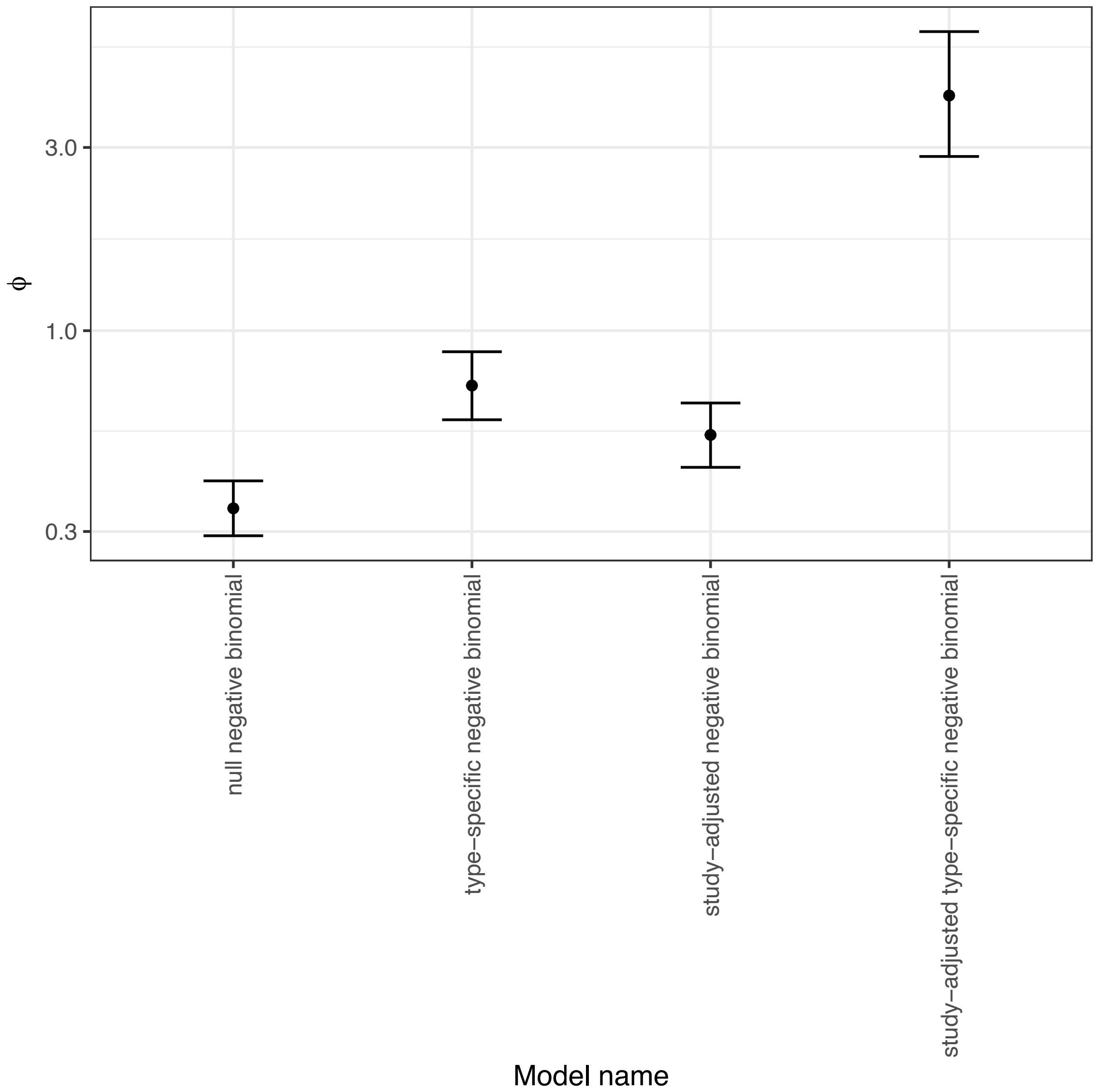


**Figure S8** Graph showing the values of the negative binomial distribution's precision parameter, ϕ, from model fits to the filtered serotype data from child carriage and disease. The points represent the median estimates from the MCMCs, and the error bars show the 95% credibility intervals.

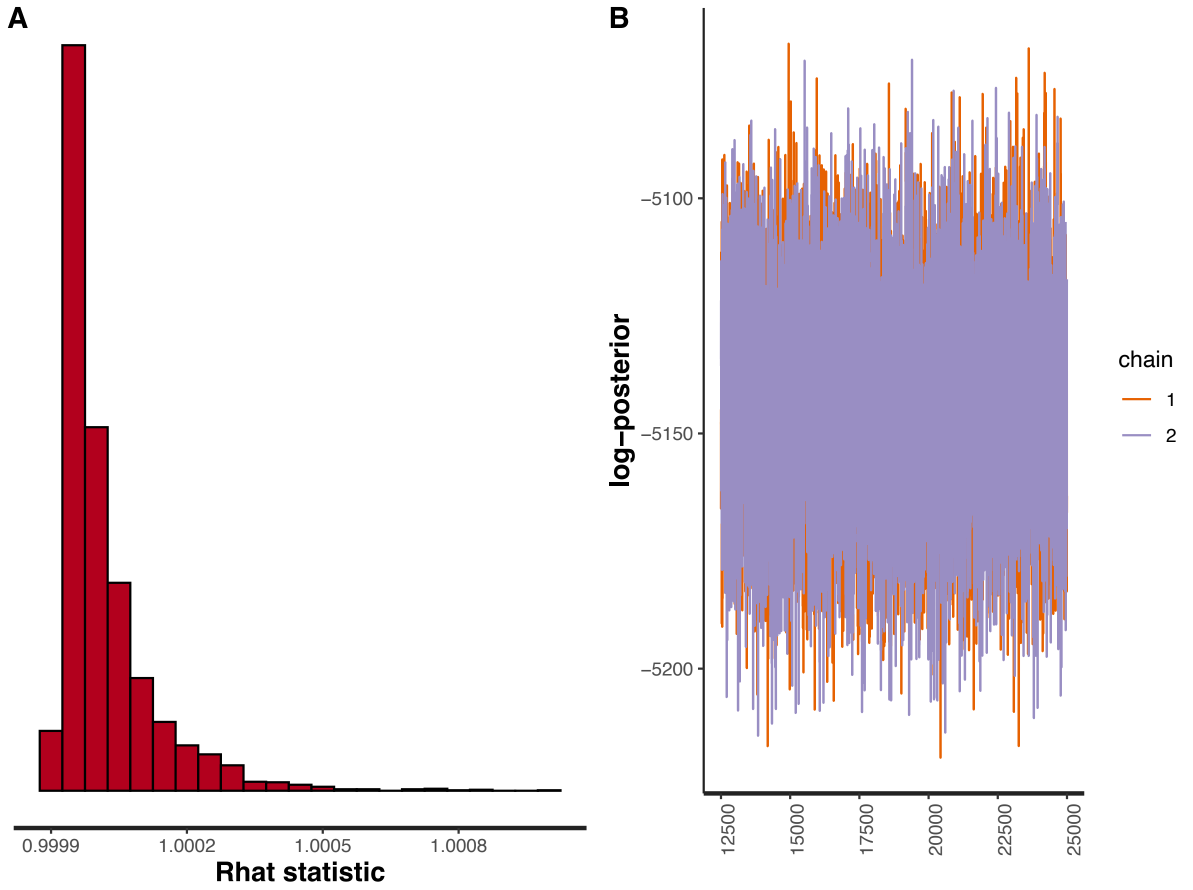


**Figure S9** Plots validating the fit of the study-adjusted type-specific negative binomial model to the full serotype data from child carriage and disease. (A) Histogram showing the distribution of Rhat values. (B) Post-warmup MCMC traces of the log posterior probability.

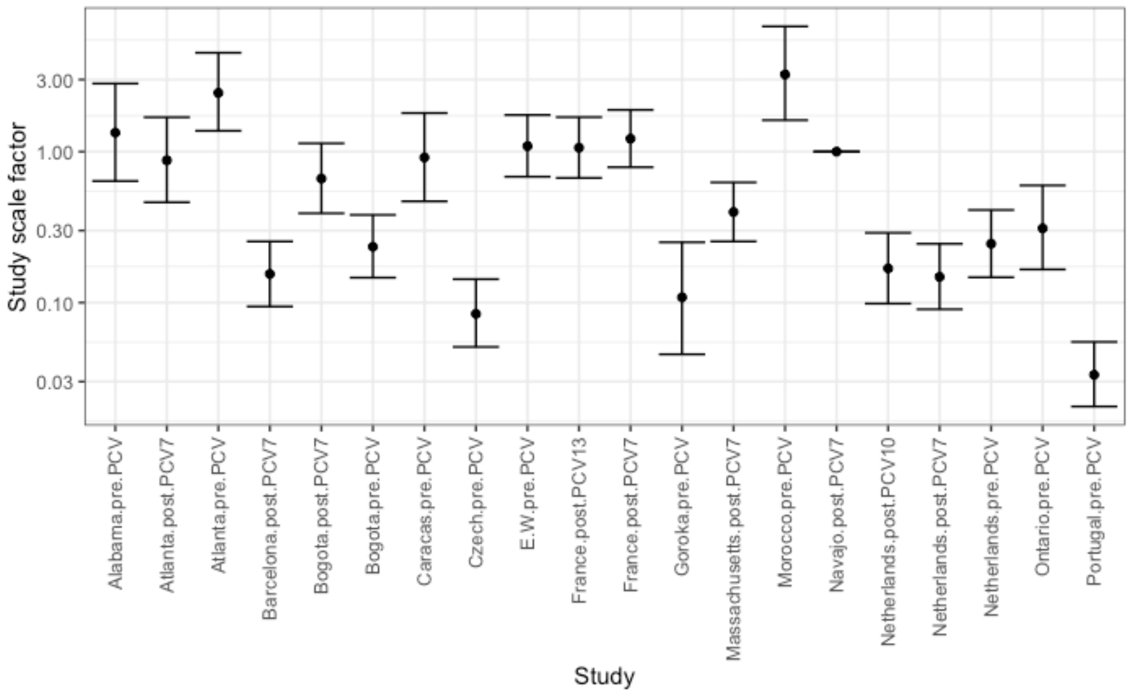


**Figure S10** Study-adjustment scale factors from the study-adjusted type-specific negative binomial model fitted to the full serotype data from child carriage and disease. The reference study, for which the value was fixed at one, was the Navajo post-PCV7 dataset, which had the greatest sample size in this meta-analysis (Fig. S1).

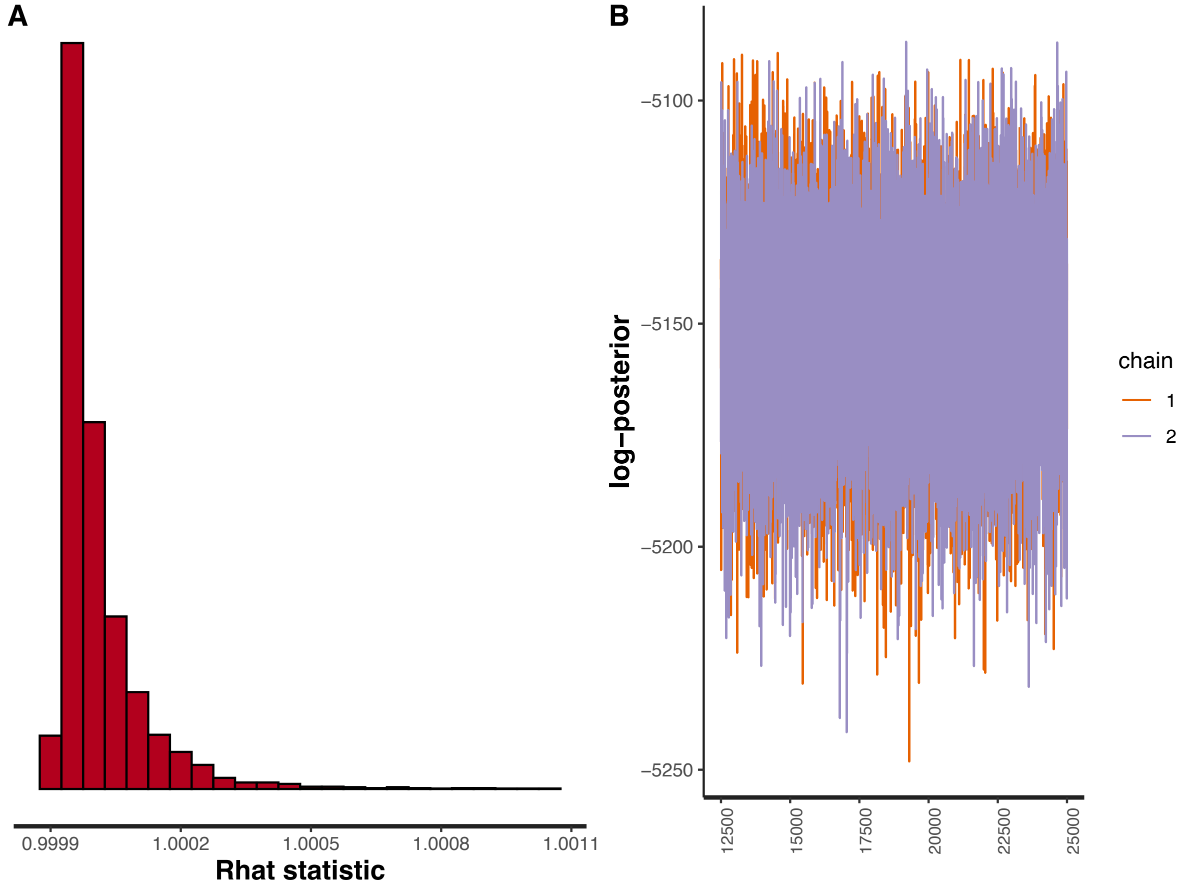


**Figure S11** Plots validating the fit of the study-adjusted type-specific negative binomial model to the full serotype data from child carriage and disease, when distinguishing between vaccine-type serotypes pre- and post-PCV. (A) Histogram showing the distribution of Rhat values. (B) Post-warmup MCMC traces of the log posterior probability.

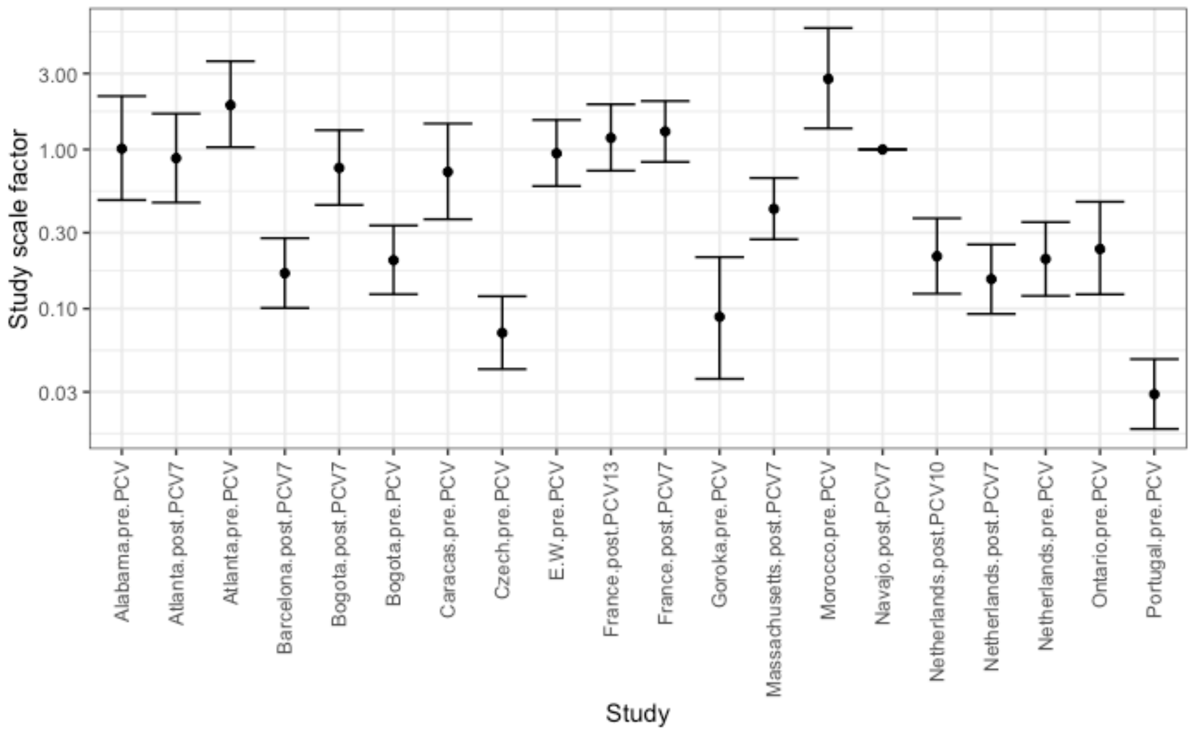


**Figure S12** Study-adjustment scale factors from the study-adjusted type-specific negative binomial model fitted to the full serotype data from child carriage and disease, when distinguishing between vaccine-type serotypes pre- and post-PCV. The reference study, for which the value was fixed at one, was the Navajo post-PCV7 dataset, which had the greatest sample size in this meta-analysis (Fig. S1).

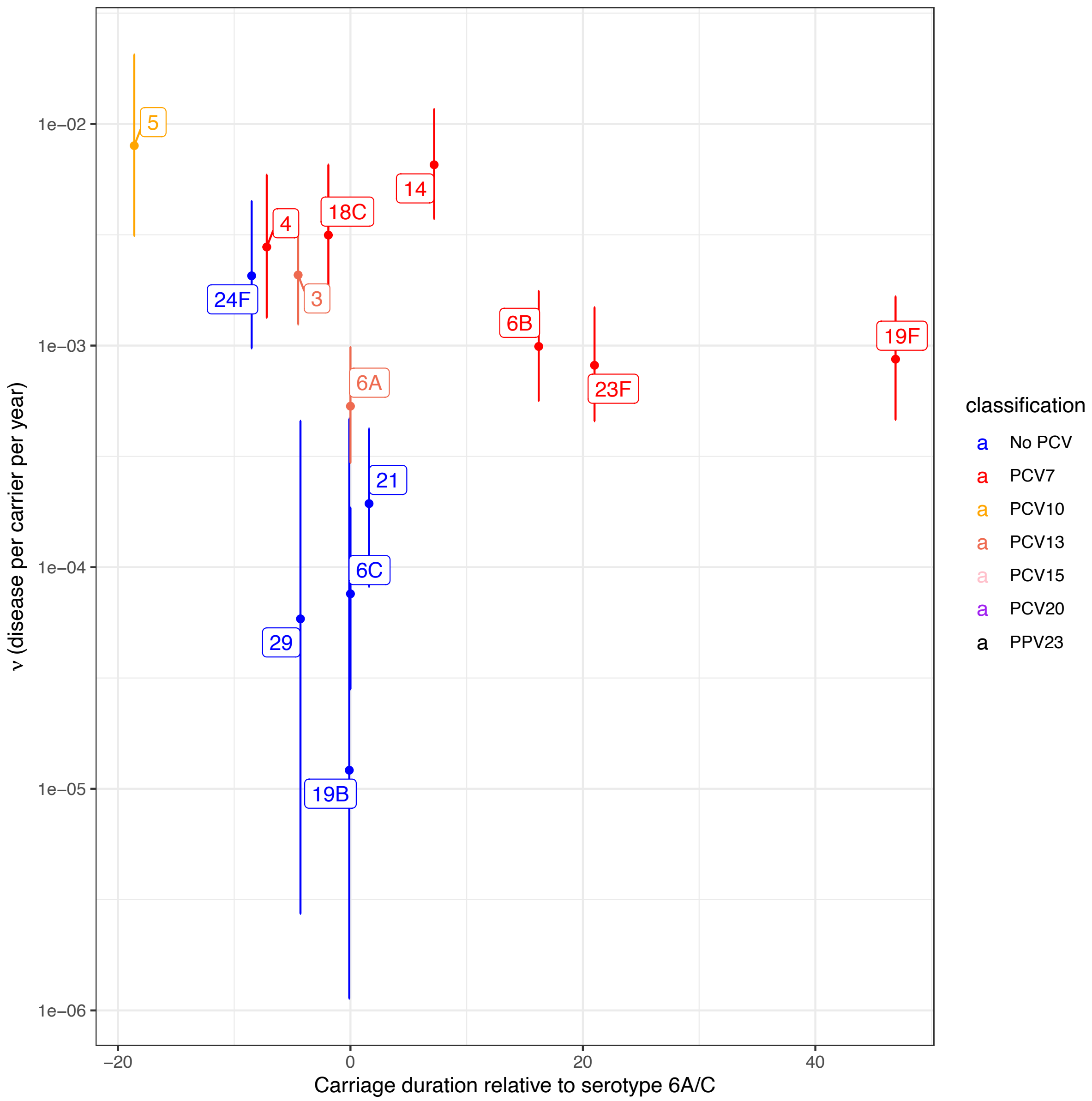


**Figure S13** Relationship between serotype invasiveness in children (as shown in Fig. 2) and carriage duration in a study of infant colonisation in Maela, Thailand. The carriage duration estimates were derived from a multi-variate lasso regression that included both serotype and antibiotic resistance phenotypes. Values were available for 14 serotypes, all relative to the carriage duration of serotype 6A/C, which was therefore assigned a value of zero days.

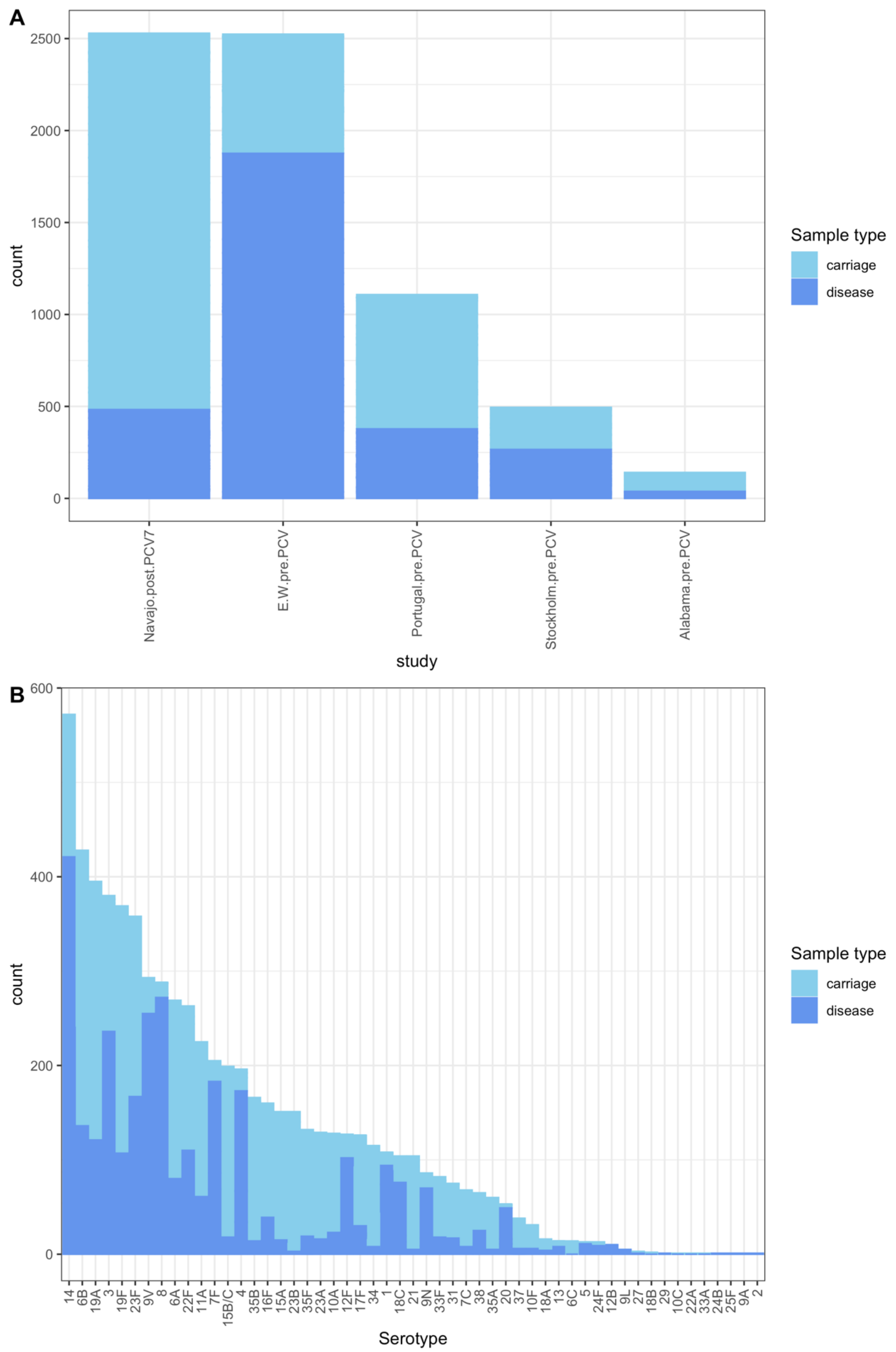


**Figure S14** Properties of the adult serotype dataset. (A) Stacked bar plot showing the distribution of carriage and disease isolates between studies. (B) Stacked bar plot showing the distribution of carriage and disease isolates between serotypes.

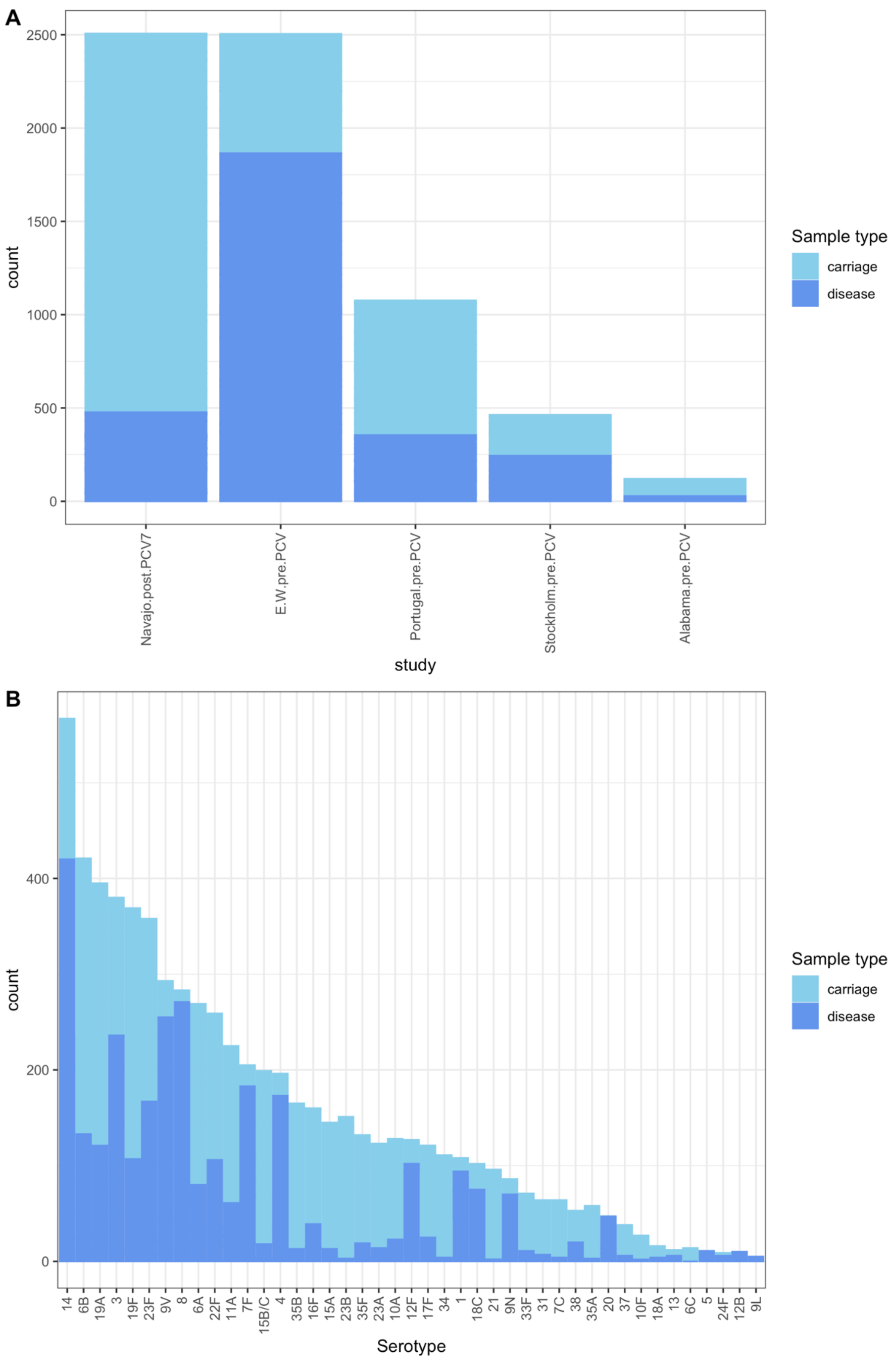


**Figure S15** Properties of the adult serotype dataset after removing datapoints in which fewer than five isolates of a serotype were recorded from both carriage and disease in a given study. (A) Stacked bar plot showing the distribution of carriage and disease isolates between studies. (B) Stacked bar plot showing the distribution of carriage and disease isolates between serotypes.

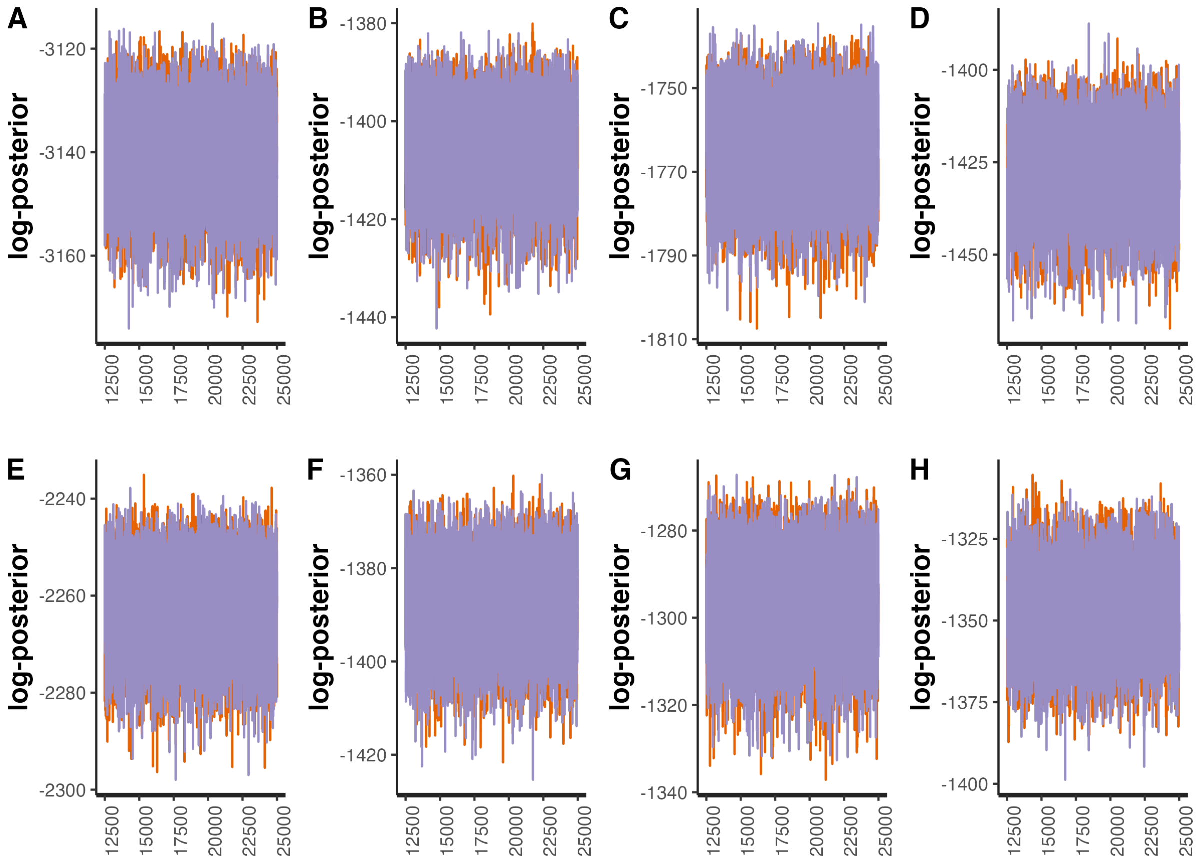


**Figure S16** Line plots showing the post-warmup MCMC traces for the logarithmic posterior probabilities across two independent chains for models fitted to the filtered serotype data from child carriage and adult disease. The horizontal axis shows the generation of the MCMC, with values for the two chains shown by orange and purple lines. Each panel corresponds to a different model: (A) null Poisson model; (B) null negative binomial model; (C) type-specific Poisson model; (D) type-specific negative binomial model; (E) study-adjusted Poisson model; (F) study-adjusted negative binomial model; (G) study-adjusted type-specific Poisson model; (H) study-adjusted type-specific negative binomial model.

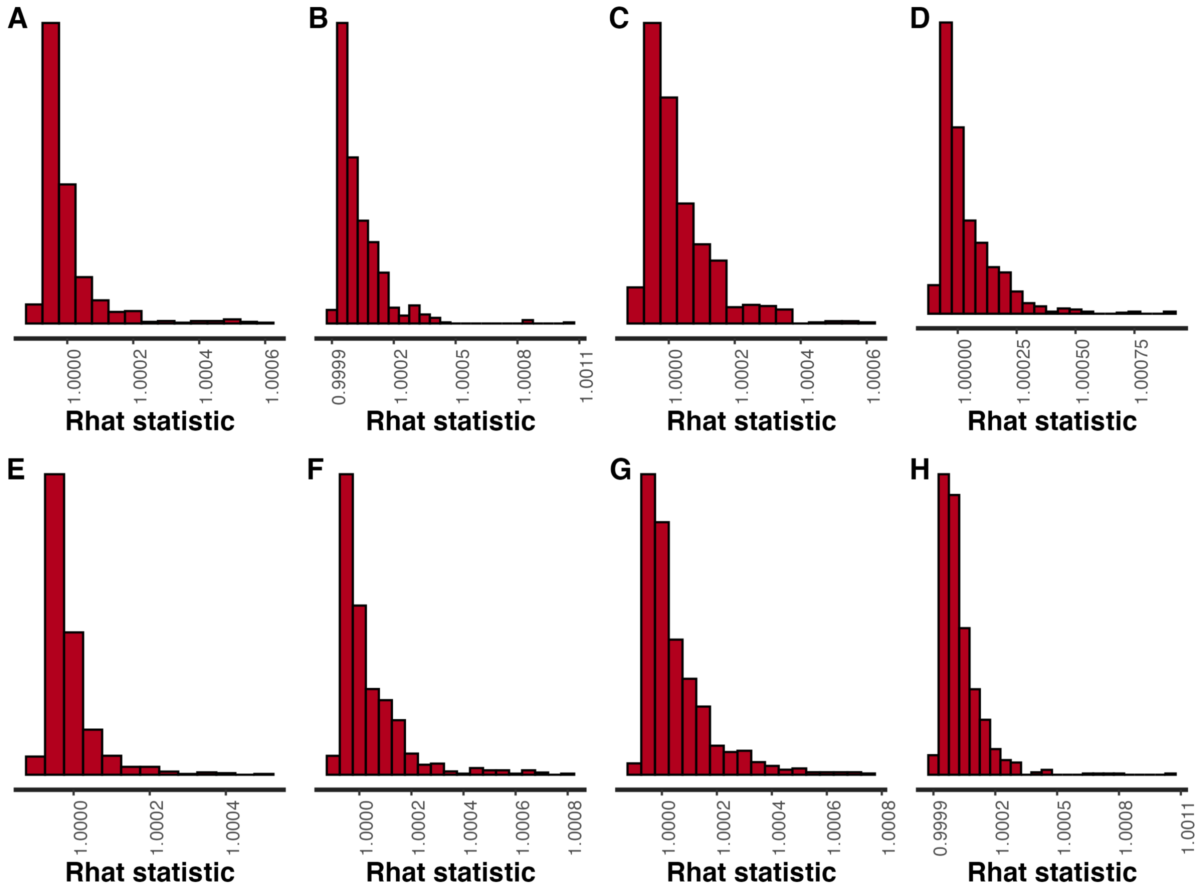


**Figure S17** Histograms of Rhat values calculated from paired MCMCs for models fitted to the filtered serotype data from child carriage and adult disease. Each panel corresponds to a different model: (A) null Poisson model; (B) null negative binomial model; (C) type-specific Poisson model; (D) type-specific negative binomial model; (E) study-adjusted Poisson model; (F) study-adjusted negative binomial model; (G) study-adjusted type-specific Poisson model; (H) study-adjusted type-specific negative binomial model.

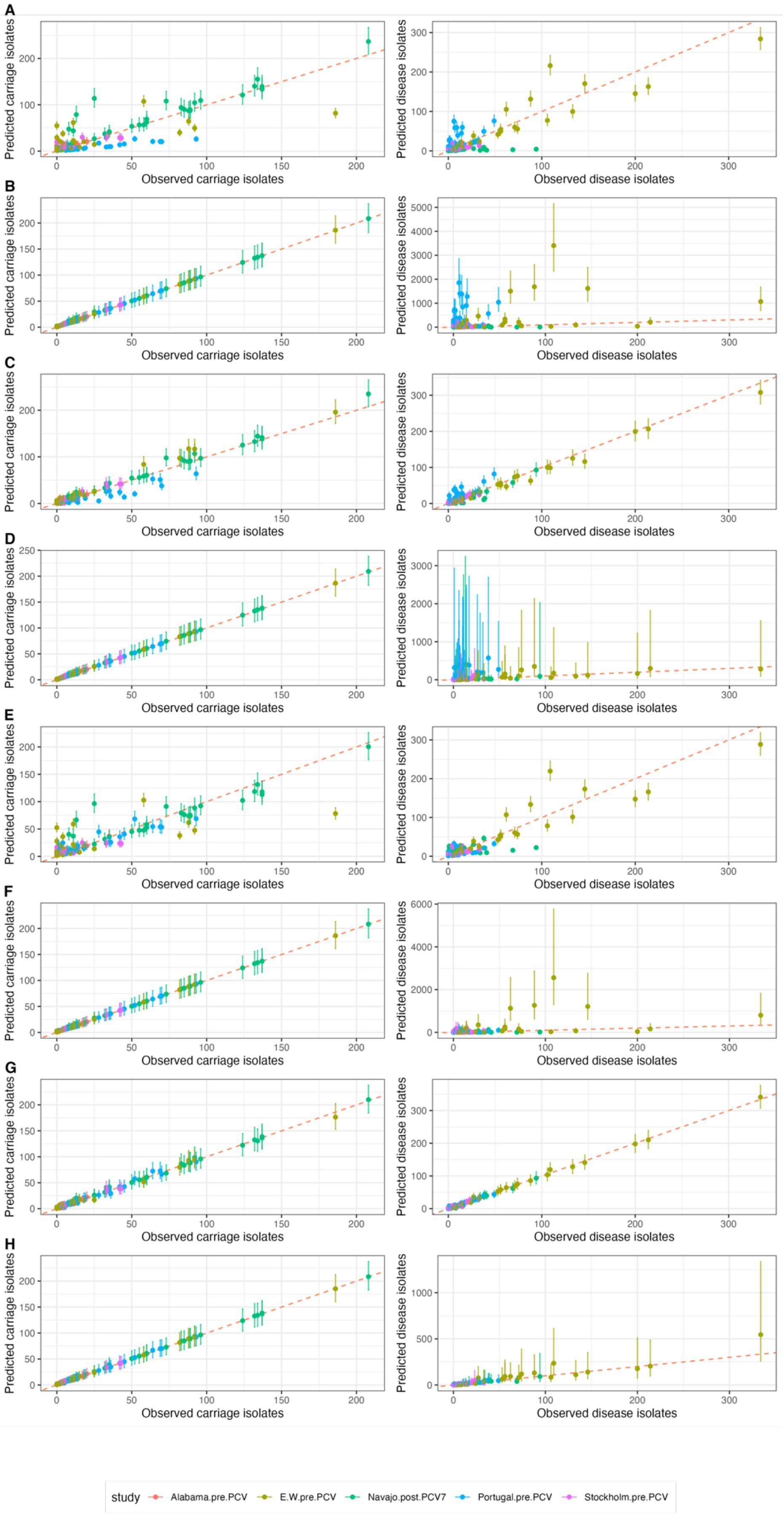

A
B
C
D
E
F
G
H
Predicted carriage isolates
Observed carriage isolates
Predicted disease isolates
Observed disease isolates
study
Alabama.pre.PCV
E.W.pre.PCV
Navajo.post.PCV7
Portugal.pre.PCV
Stockholm.pre.PCV

**Figure S18** Comparison of observed and predicted counts of each serotype within each study of adult disease relative to carriage in children. The plots in the left column display data on carriage in children. The plots in the right column display data on disease in adults. The points are coloured by the study to which they correspond, and represent the observed value on the horizontal axis, and the median predicted value on the vertical axis. The error bars show the 95% credibility intervals. The red dashed line shows the line of identity, corresponding to a perfect match between prediction and observation. The left column shows the correspondence for carriage data (values of $c_{i,j}$), and the right column shows the correspondence for disease isolates (values of $d_{i,j}$). Each row corresponds to a different model: (A) null Poisson model; (B) null negative binomial model; (C) type-specific Poisson model; (D) type-specific negative binomial model; (E) study-adjusted Poisson model; (F) study-adjusted negative binomial model; (G) study-adjusted type-specific Poisson model; (H) study-adjusted type-specific negative binomial model.

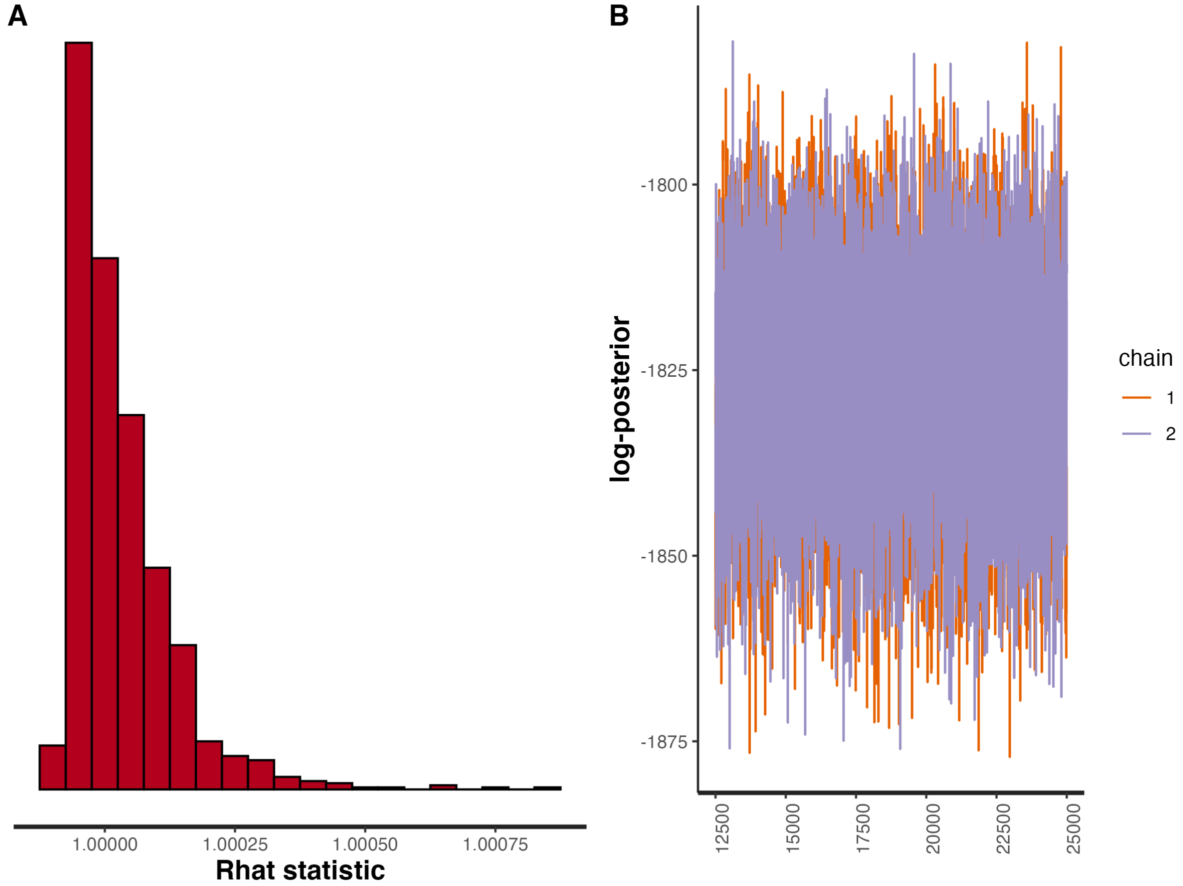


**Figure S19** Plots validating the fit of the study-adjusted type-specific negative binomial model to the full serotype data from child carriage and adult disease. (A) Histogram showing the distribution of Rhat values. (B) Post-warmup MCMC traces of the log posterior probability.

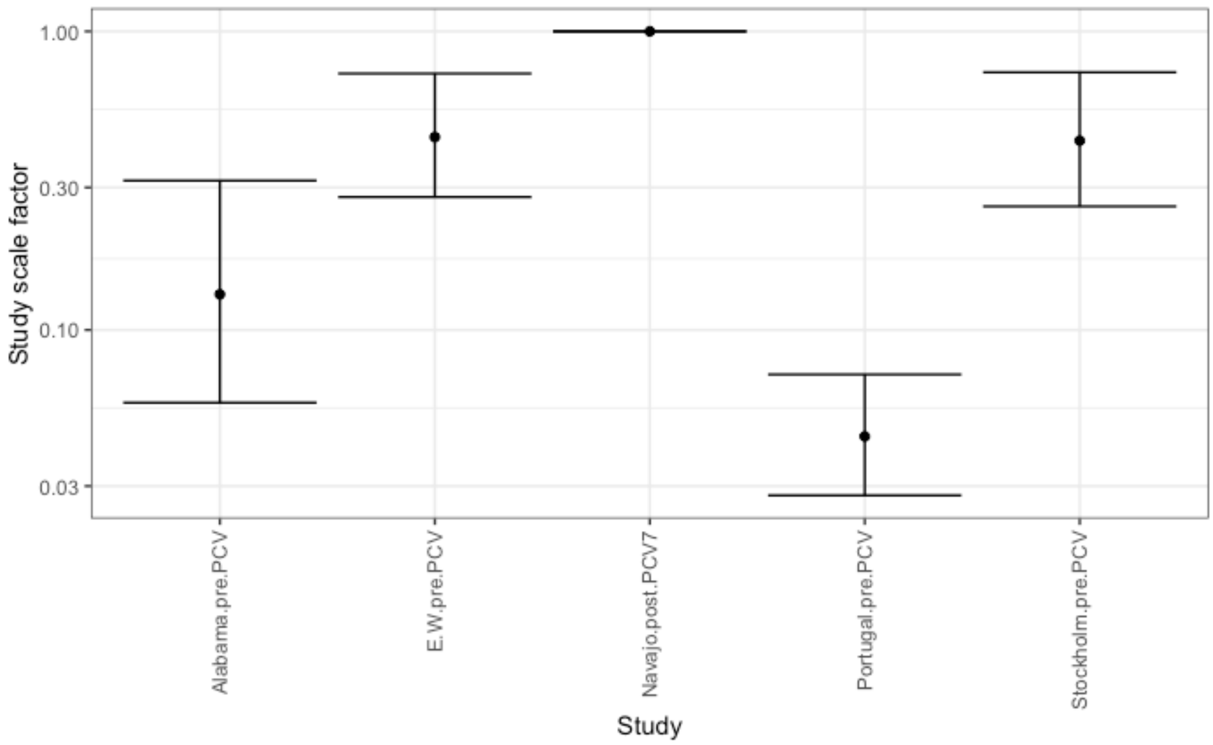


**Figure S20** Study-adjustment scale factors from the study-adjusted type-specific negative binomial model fitted to the full serotype data from child carriage and adult disease. The reference study, for which the value was fixed at one, was the Navajo post-PCV7 dataset, which had the greatest sample size in this meta-analysis (Fig. S15).

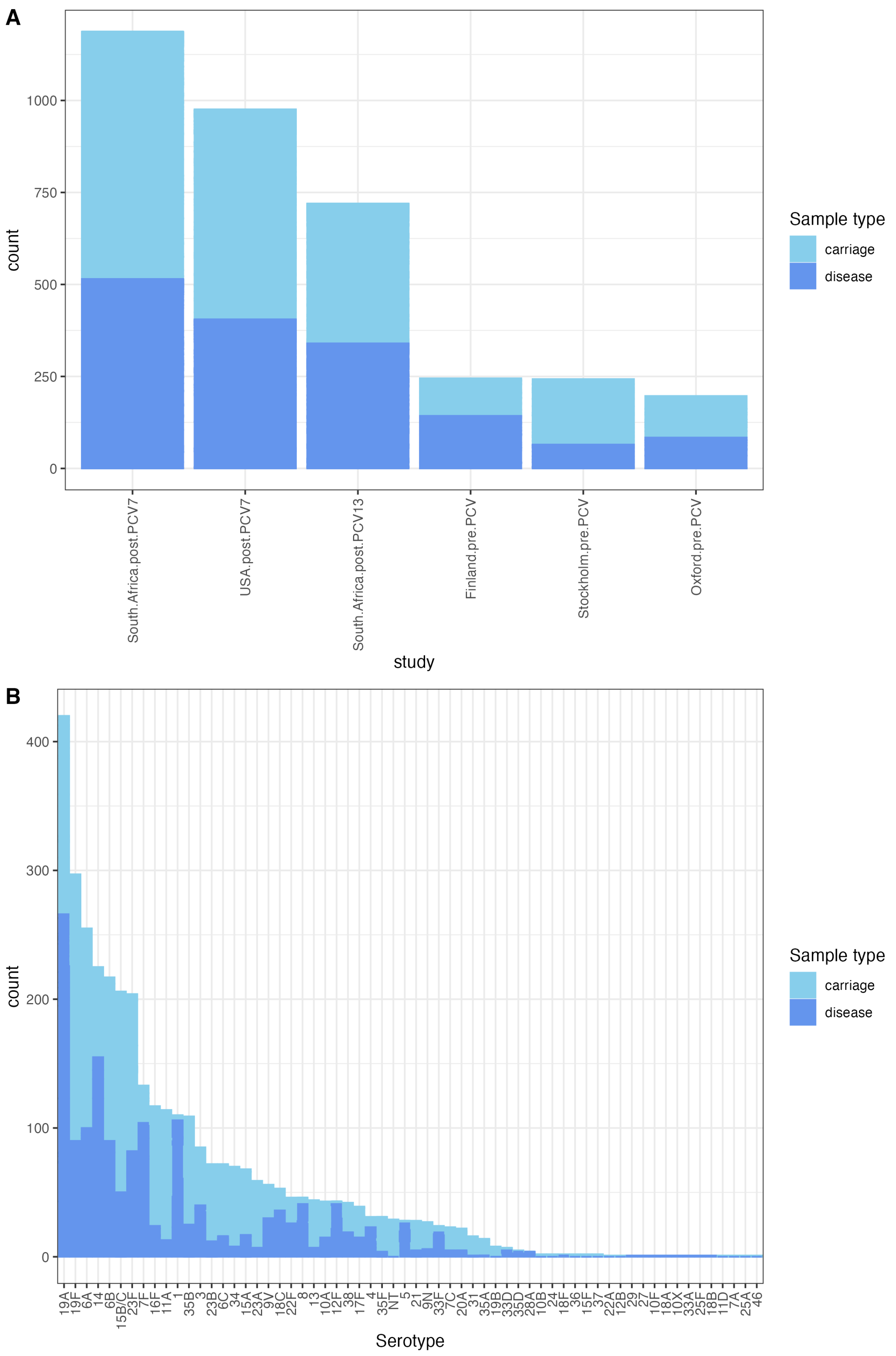


**Figure S21** Properties of the child serotype and strain dataset. (A) Stacked bar plot showing the distribution of carriage and disease isolates between studies. (B) Stacked bar plot showing the distribution of carriage and disease isolates between serotypes.

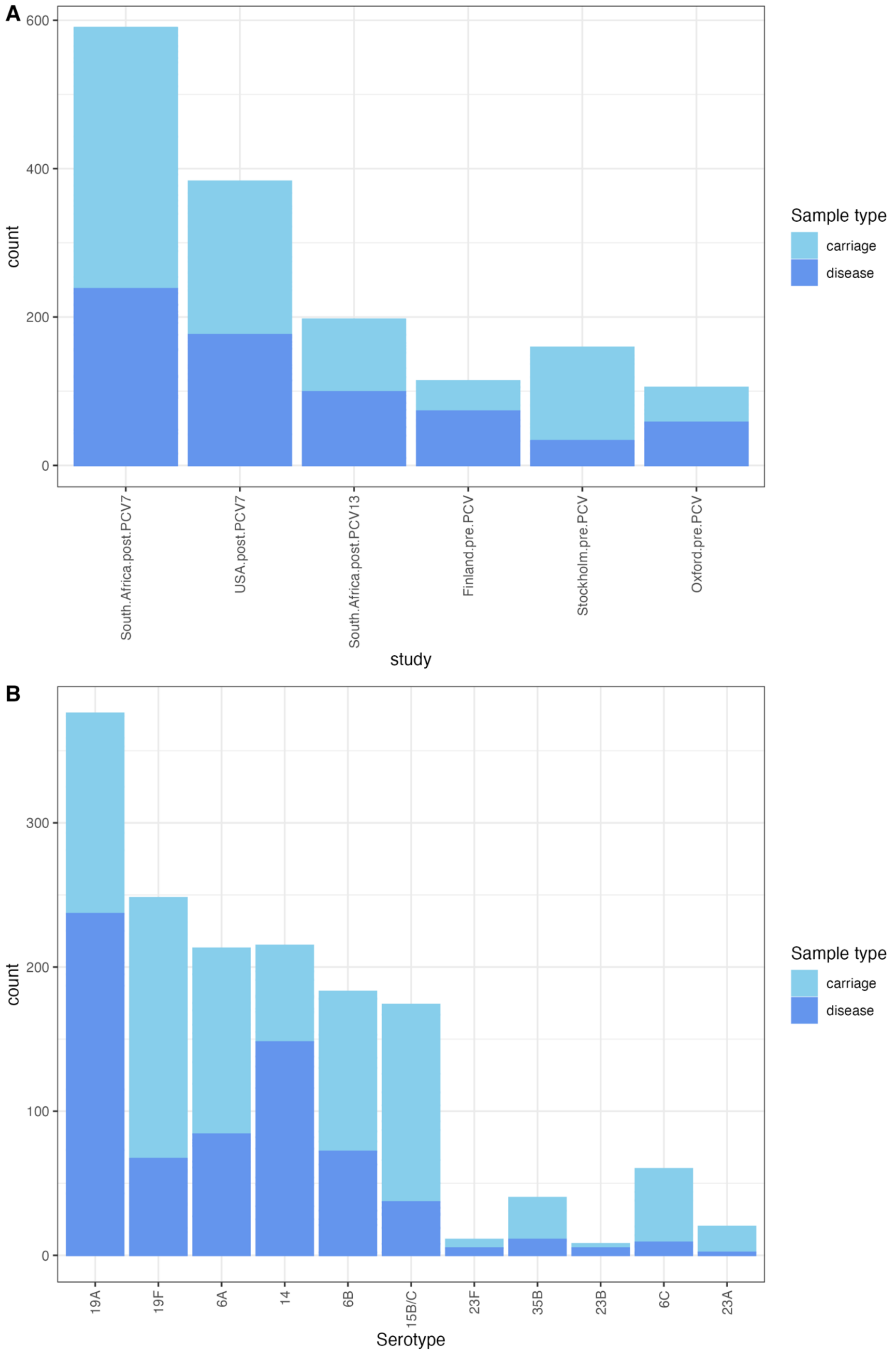


**Figure S22** Properties of the child serotype and strain dataset after removing datapoints in which fewer than five isolates of a serotype-strain combination were recorded from both carriage and disease in a given study, then retaining only serotypes associated with at least five different strains, and strains associated with at least five different serotypes. (A) Stacked bar plot showing the distribution of carriage and disease isolates between studies. (B) Stacked bar plot showing the distribution of carriage and disease isolates between serotypes.

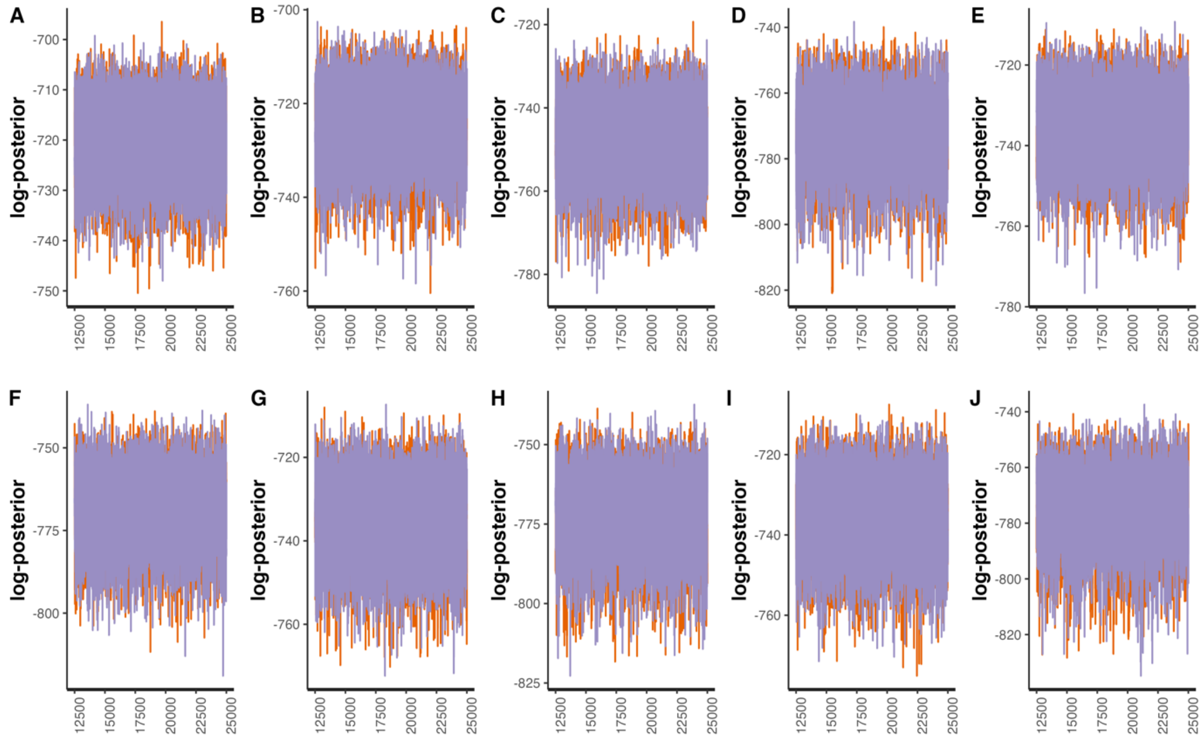


**Figure S23** Line plots showing the post-warmup MCMC traces for the logarithmic posterior probabilities across two independent chains for models fitted to the filtered strain and serotype data from child carriage and disease. Each panel corresponds to a model with a different method of associating isolates with an invasiveness estimate: (A) serotype-determined, Poisson-distributed invasiveness; (B) serotype-determined, negative binomially-distributed invasiveness; (C) strain-determined, Poisson-distributed invasiveness; (D) strain-determined, negative binomially-distributed invasiveness; (E) serotype-determined, strain-modified Poisson-distributed invasiveness; (F) serotype-determined, strain-modified negative binomially-distributed invasiveness; (G) strain-determined, serotype-modified Poisson-distributed invasiveness; (H) strain-determined, serotype-modified negative binomially-distributed invasiveness; (I) strain- and serotype-determined Poisson-distributed invasiveness; (J) strain- and serotype-determined negative binomially-distributed invasiveness.

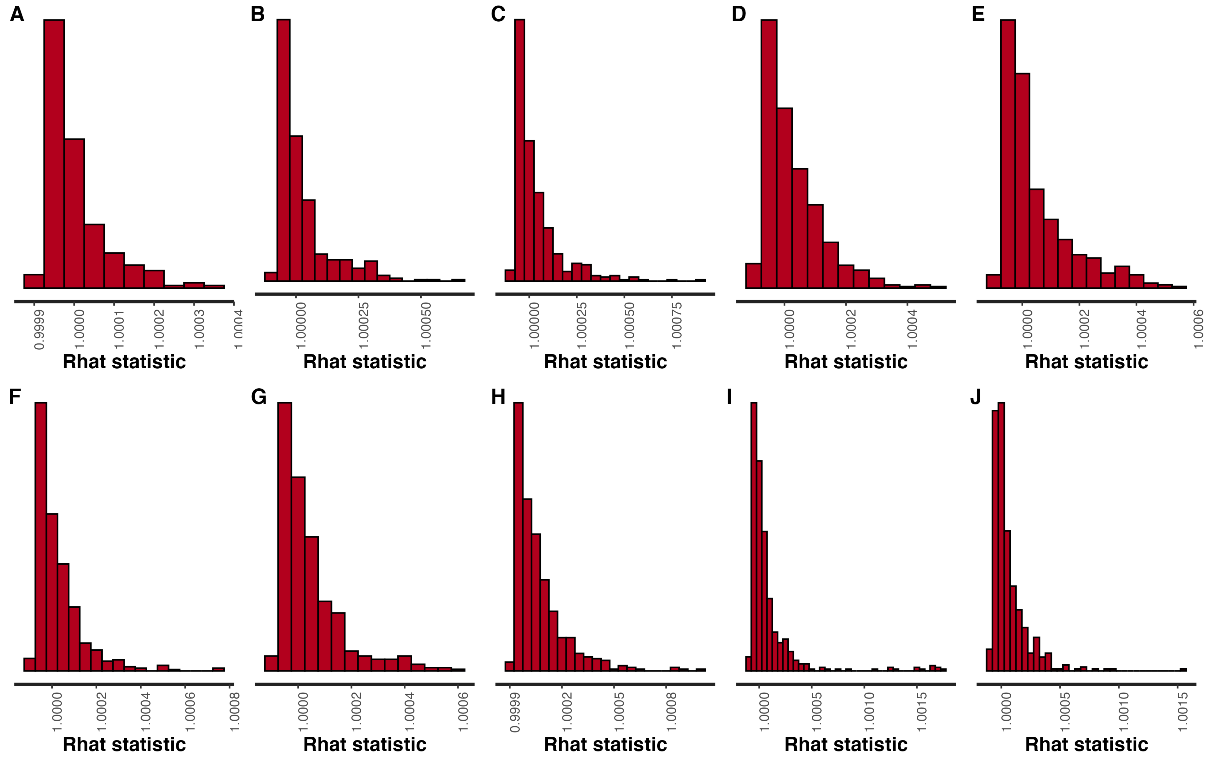


**Figure S24** Histograms of Rhat values calculated from paired MCMCs for models fitted to the filtered strain and serotype data from child carriage and disease. Each panel corresponds to a different model: (A) serotype-determined, Poisson-distributed invasiveness; (B) serotype-determined, negative binomially-distributed invasiveness; (C) strain-determined, Poisson-distributed invasiveness; (D) strain-determined, negative binomially-distributed invasiveness; (E) serotype-determined, strain-modified Poisson-distributed invasiveness; (F) serotype-determined, strain-modified negative binomially-distributed invasiveness; (G) strain-determined, serotype-modified Poisson-distributed invasiveness; (H) strain-determined, serotype-modified negative binomially-distributed invasiveness; (I) strain- and serotype-determined Poisson-distributed invasiveness; (J) strain- and serotype-determined negative binomially-distributed invasiveness.

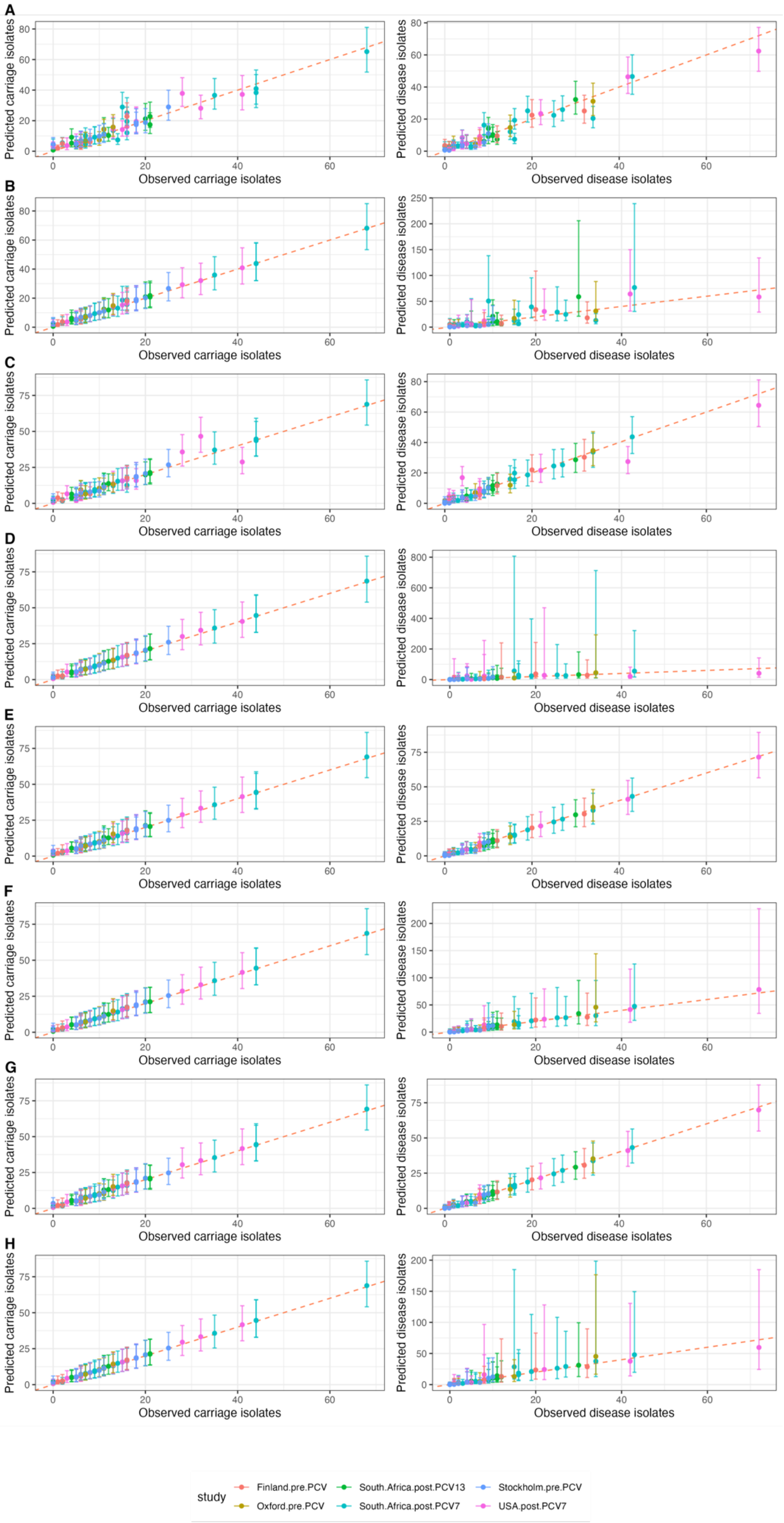
A
B
C
D
E
F
G
H
Predicted carriage isolates
Observed carriage isolates
Predicted disease isolates
Observed disease isolates
study
Finland.pre.PCV
Oxford.pre.PCV
South.Africa.post.PCV13
South.Africa.post.PCV7
Stockholm.pre.PCV
USA.post.PCV7

**Figure S25** Comparison of observed and predicted counts of isolates, categorised by both their serotype and strain background, within each study. The points are coloured by the study to which they correspond, and represent the observed value on the horizontal axis, and the median predicted values on the vertical axis. The error bars show the 95% credibility intervals. The red dashed line shows the line of identity, corresponding to a perfect match between prediction and observation. The left column shows the correspondence for carriage data (values of $c_{i,j,k}$), and the right column shows the correspondence for disease isolates (values of $d_{i,j,k}$). Each row corresponds to a different model: (A) serotype-determined, Poisson-distributed invasiveness; (B) serotype-determined, negative binomially-distributed invasiveness; (C) strain-determined, Poisson-distributed invasiveness; (D) strain-determined, negative binomially-distributed invasiveness; (E) serotype-determined, strain-modified Poisson-distributed invasiveness; (F) serotype-determined, strain-modified negative binomially-distributed invasiveness; (G) strain-determined, serotype-modified Poisson-distributed invasiveness; (H) strain-determined, serotype-modified negative binomially-distributed invasiveness; (I) strain- and serotype-determined Poisson-distributed invasiveness; (J) strain- and serotype-determined negative binomially-distributed invasiveness.

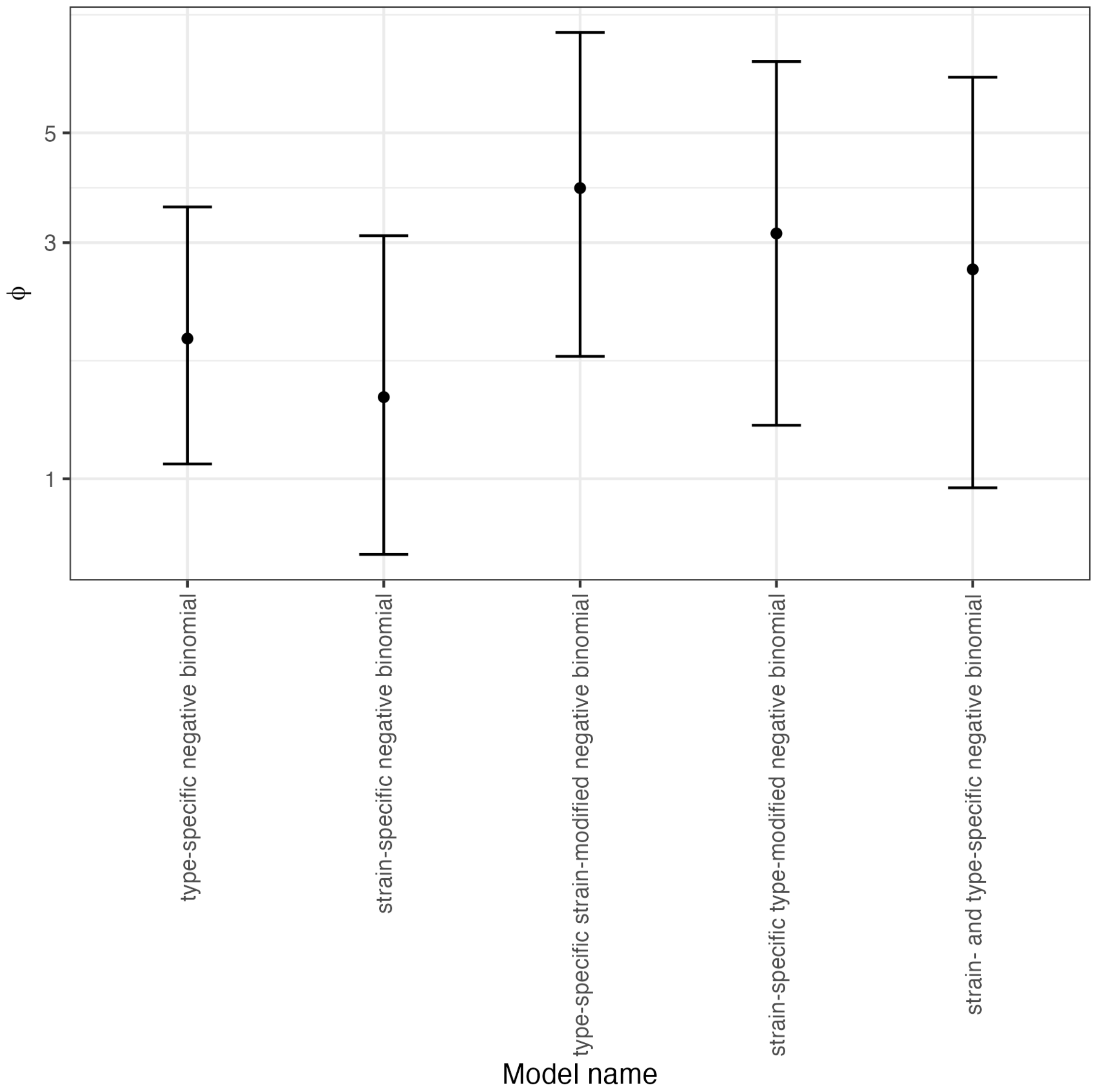


**Figure S26** Graph showing the values of the negative binomial distribution's precision parameter, ϕ, from model fits to the filtered strain and serotype data from child carriage and disease. The points represent the median estimates from the MCMCs, and the error bars show the 95% credibility interval.

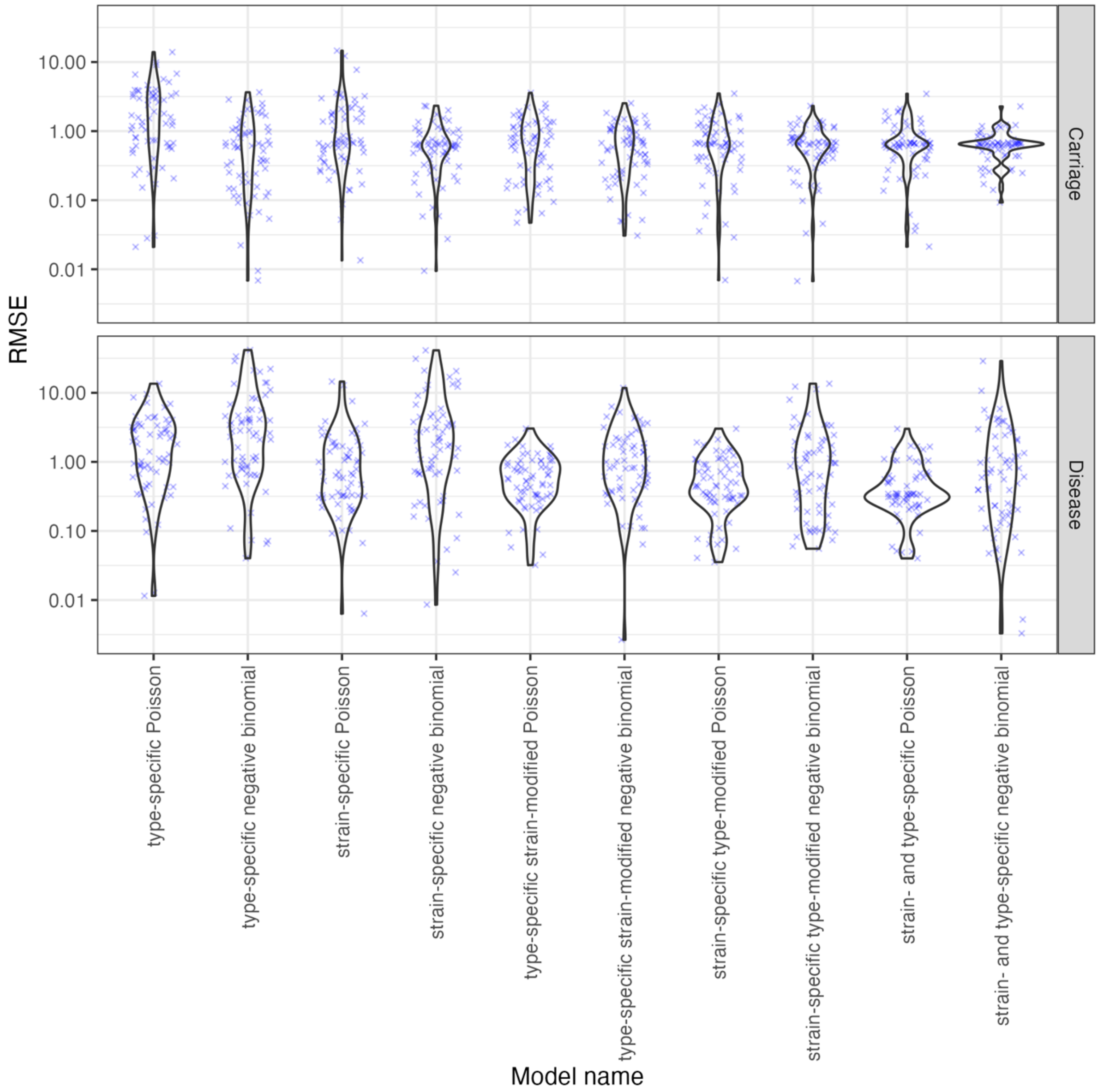


**Figure S27** Violin plots showing the root mean square error between observed and predicted values across model fits to the filtered strain and serotype data from child carriage and disease. Blue crosses represent the individual observations.

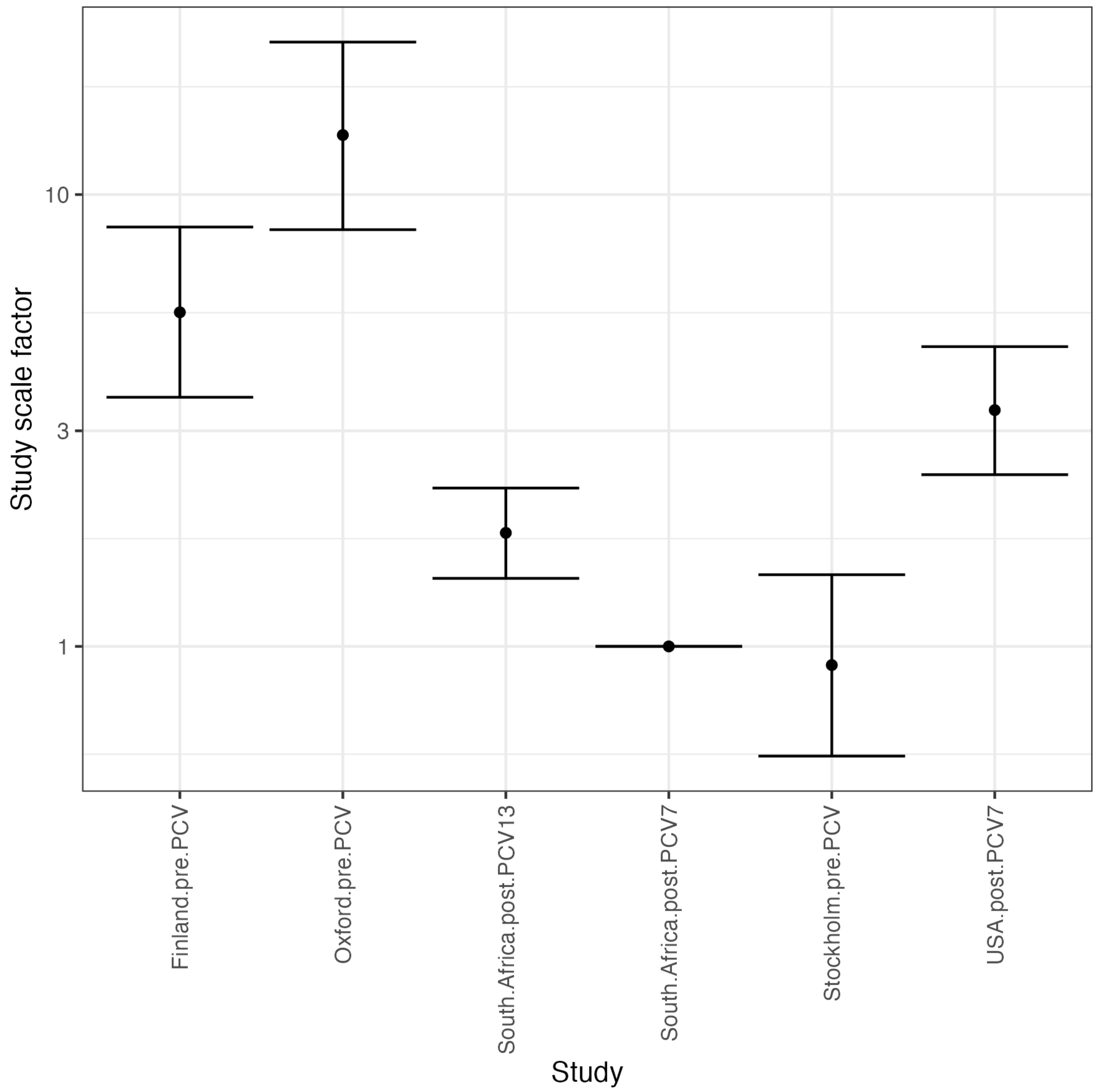


**Figure S28** Study-adjustment scale factors from the study-adjusted type-determined strain-modified Poisson model fitted to the full serotype and strain data from child carriage and disease. The reference study, for which the value was fixed at one, was the South Africa post-PCV7 dataset, which had the greatest sample size in this meta-analysis, and was associated with a known carriage sample size (Text S2).

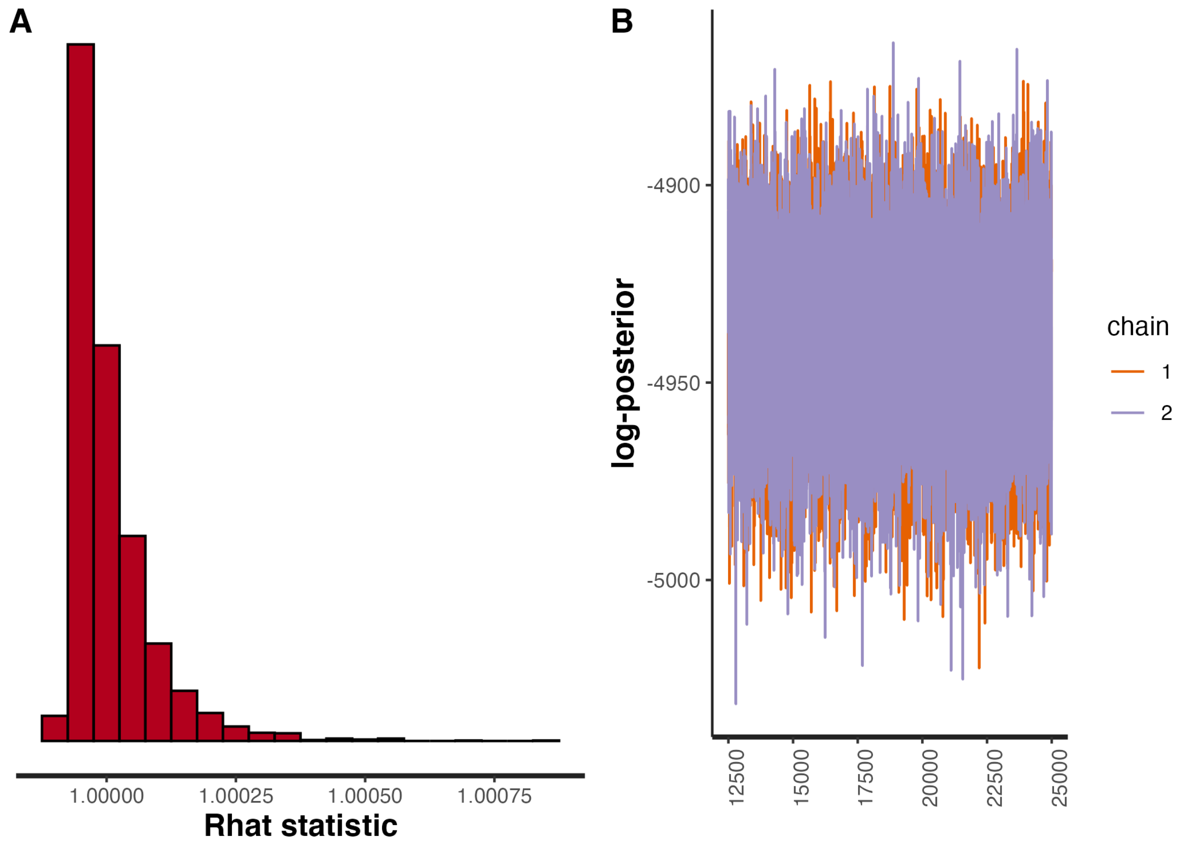


**Figure S29** Plots validating the fit of the study-adjusted type-determined strain-modified Poisson model to the full strain and serotype data from child carriage and disease. (A) Histogram showing the distribution of Rhat values. (B) Post-warmup MCMC traces of the log posterior probability.

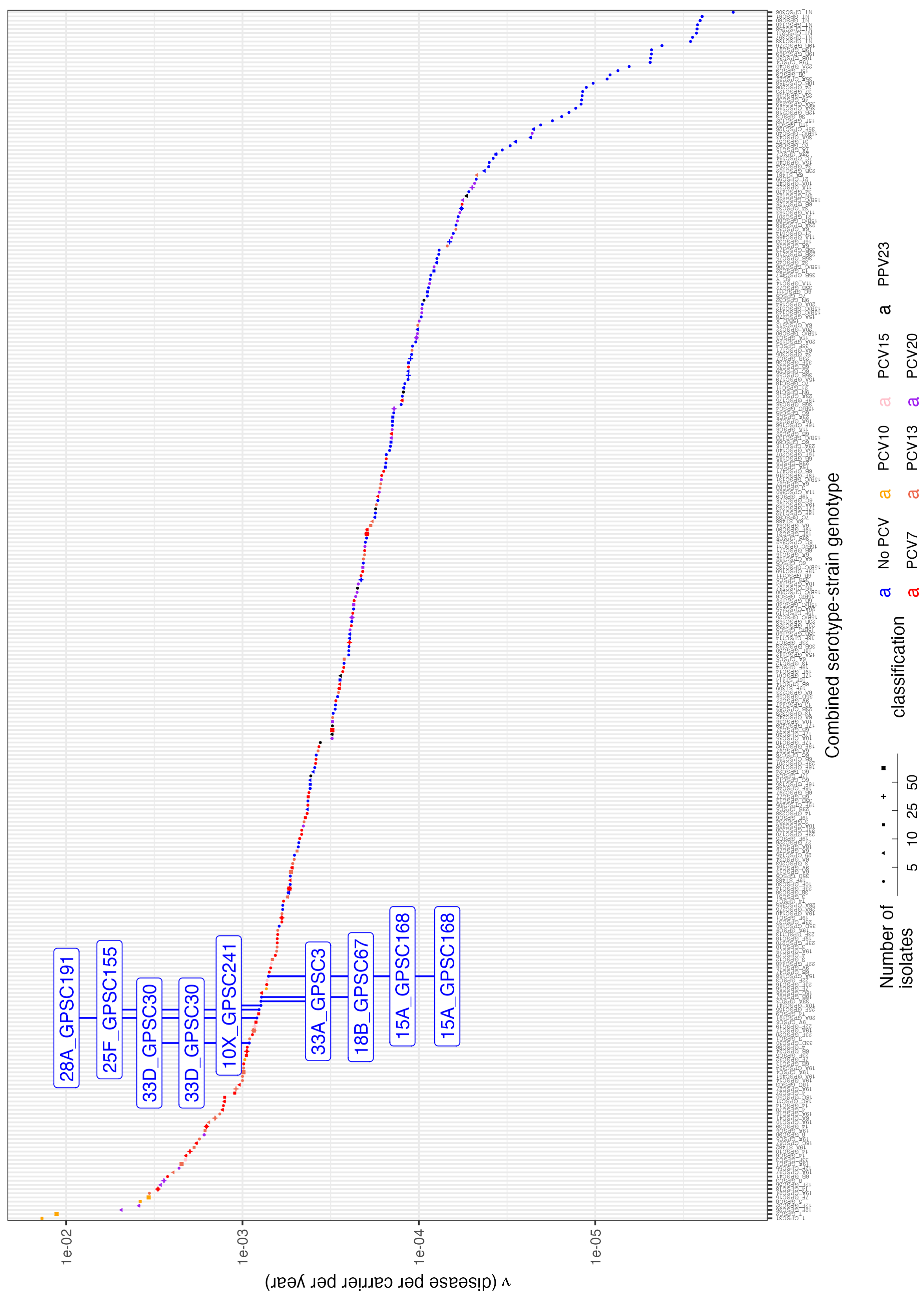


**Figure S30** Point estimates of invasiveness for all strain and serotype combinations in the full dataset from child carriage and disease. The shape of each point represents the number of isolates of each combination observed across carriage and disease samples. The serotype-strain combinations with the highest combined invasiveness that are not targeted by current PCV designs are labelled.

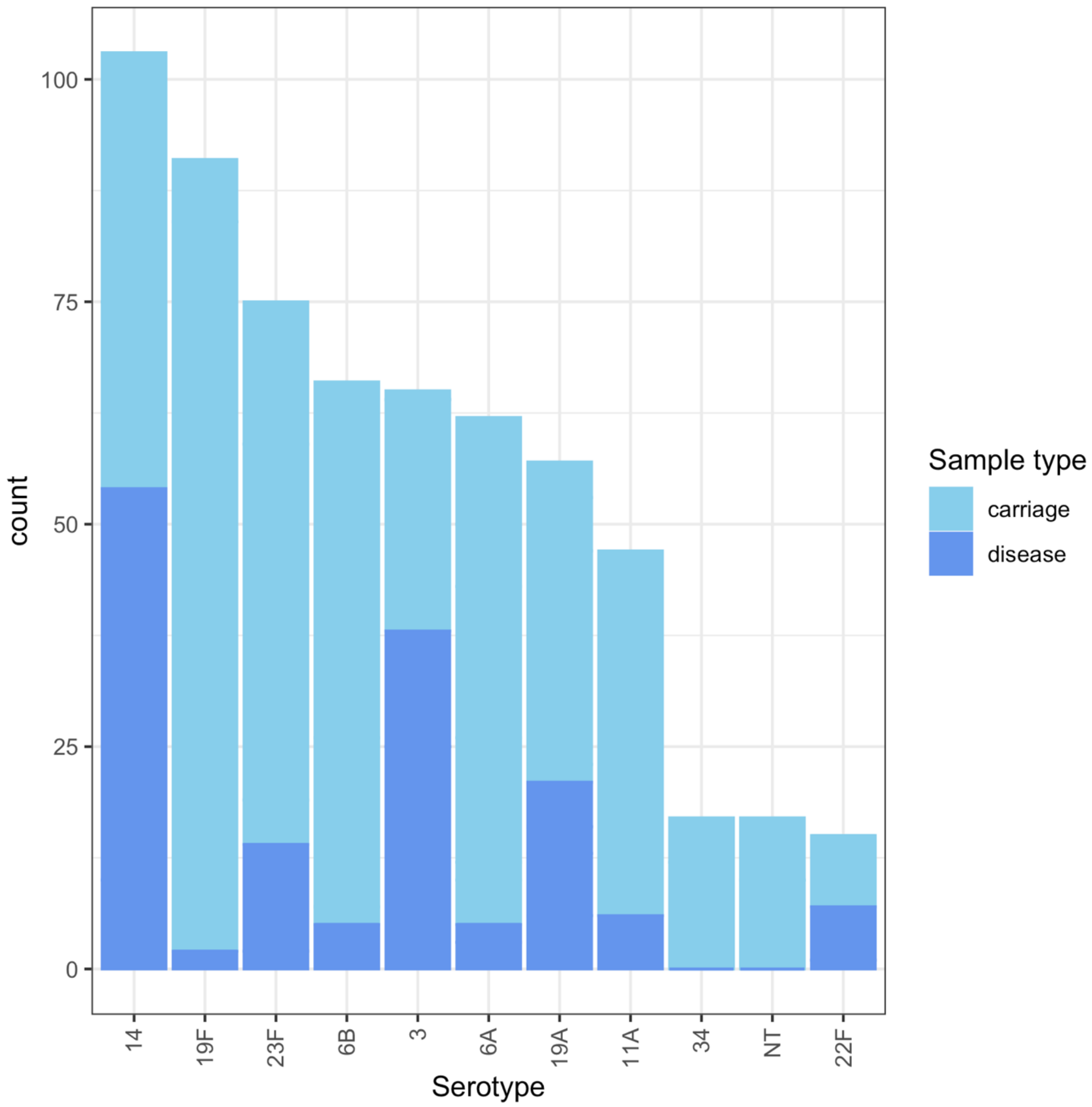


**Figure S31** Bar plot showing the distribution of carriage and disease isolates between serotypes for the dataset from Portugal comparing child carriage with primarily adult disease.

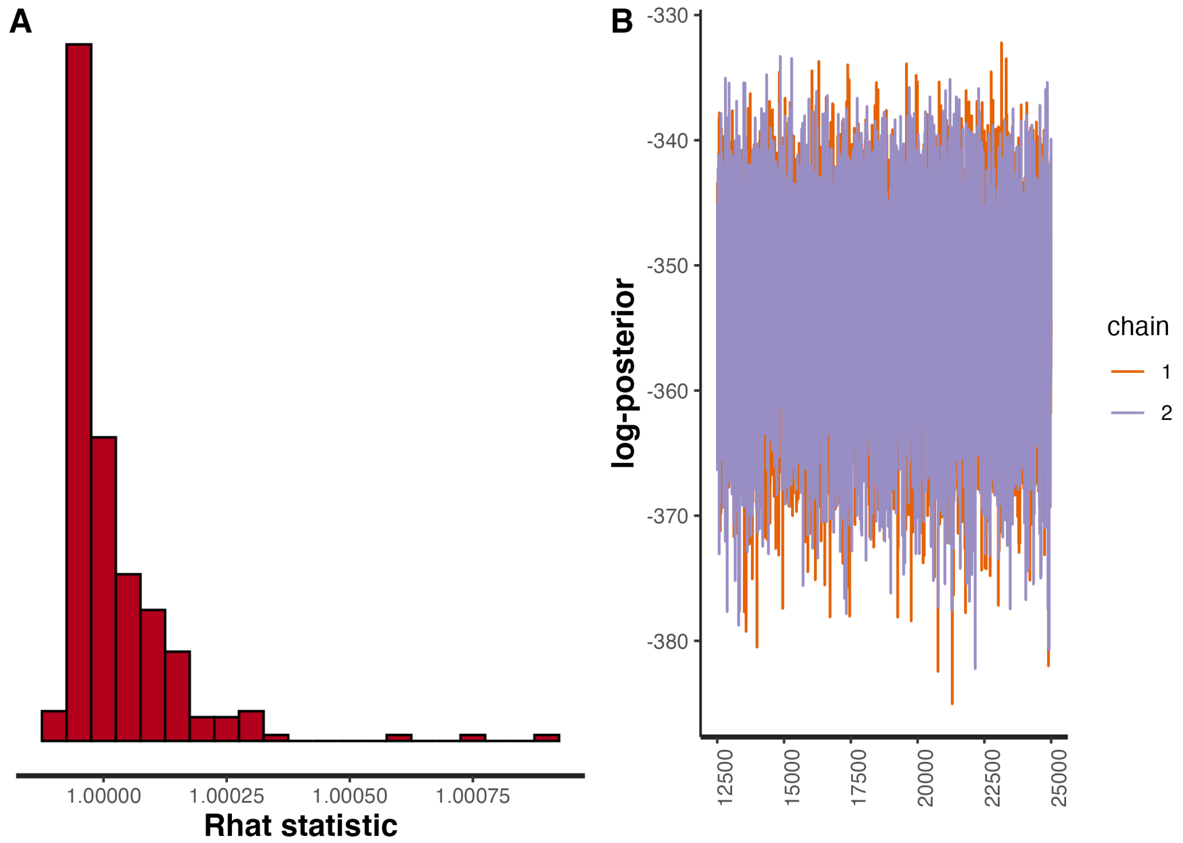


**Figure S32** Plots validating the fit of the type-determined strain-modified Poisson model to the strain and serotype data from child carriage and adult disease, with the combination of both serotype and strain determining an isolate's invasiveness. (A) Histogram showing the distribution of Rhat values. (B) Post-warmup MCMC traces of the log posterior probability.

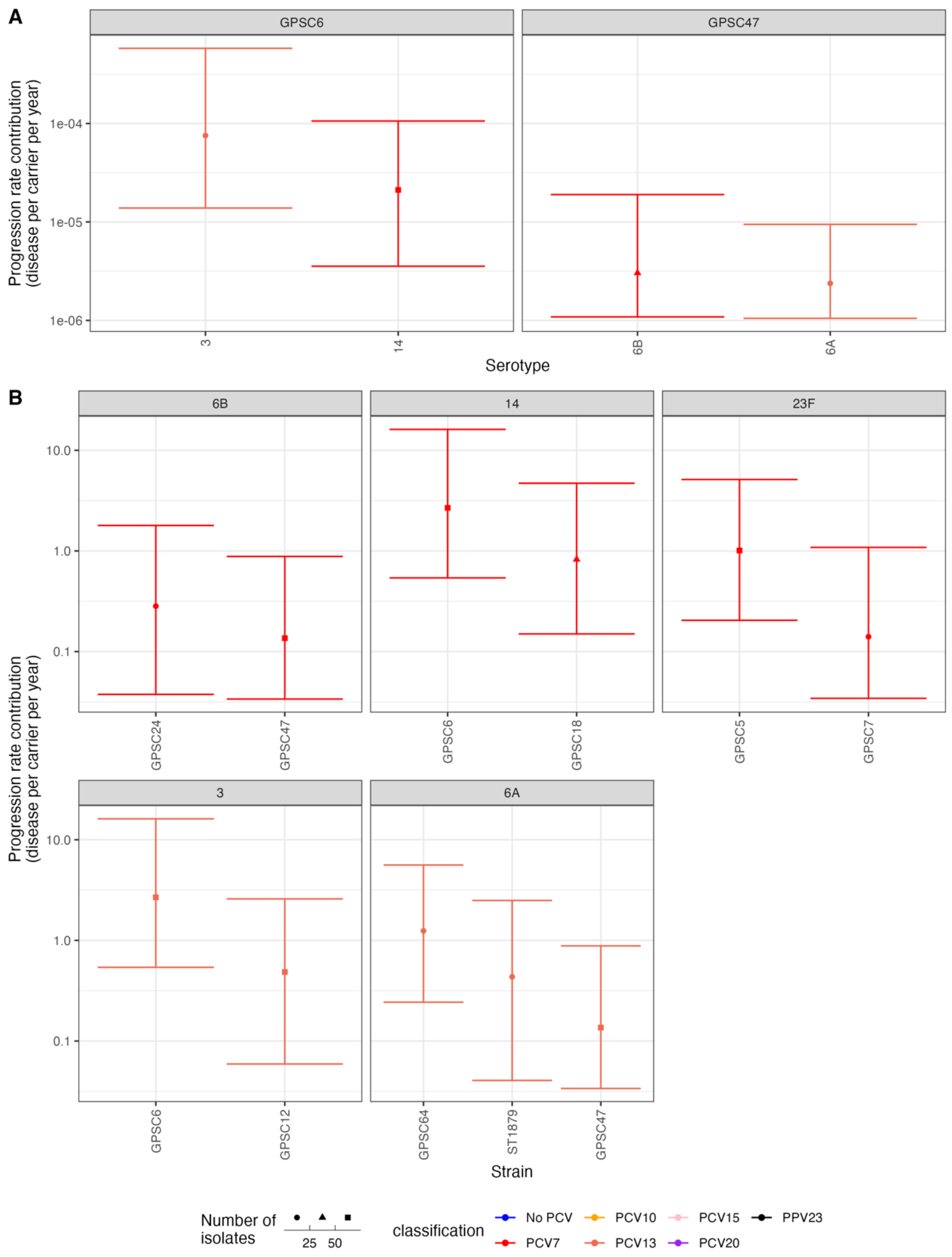


**Figure S33** Invasiveness estimates for strain and serotype combinations represented by at least ten isolates in the study of child carriage and adult disease. Points represent the median estimates, and are coloured by the vaccine formulations in which the corresponding serotype is present. The error bars represent 95% credibility intervals. The shape of the point represents the sample size on which the estimate is based. (A) Estimates of the invasiveness associated with serotypes, arranged by the strains in which they are found. Only strains expressing multiple serotypes are displayed. (B) Estimates of the coefficient by which strains modify the invasiveness of their expressed serotype, arranged by the serotypes with which they are associated. Only serotypes associated with multiple strains are displayed.

### Supplementary Tables

| Population | Vaccine period | Carriage study time interval | Carriage isolate source | Disease isolate source | No. of swabs | Population of children | No. disease isolates from children | Population of adults | No. disease isolates from adults | References |
|---|---|---|---|---|---|---|---|---|---|---|
| Alabama | Pre-PCV | July 1975 – December 1978 | Unvaccinated sick and healthy children <18 years old from Alabama | Unvaccinated children with IPD <18 years old from Alabama and unvaccinated adults hospitalised with pneumonia or IPD in Alabama | 827 | 19,316 | 114 | 232,373 | 86 | [113,114] |
| Atlanta | Pre-PCV | January – December 1995 | Unvaccinated children with recent URI in Atlanta <5 years old | Unvaccinated children <5 years old with IPD in Atlanta | 231 | 204,680 | 202 | - | - | [115] |
| Bogota | Pre-PCV | May 2005 – November 2006 | Healthy unvaccinated children <18 months old in Bogota | IPD in children <2 years old in Bogota | 197 | 357,200 | 353 | - | - | [116] |
| Caracas | Pre-PCV | December 2006 – January 2008 | Unvaccinated healthy children <6 years old in Caracas | Unvaccinated children with IPD <6 years old in Caracas | 1,004 | 146,125 | 36 | - | - | [117,118] |
| Czech | Pre-PCV | 1996 – 2005 | Unvaccinated healthy children 3 - 5 years old across the Czech Republic | Unvaccinated children <6 years old with IPD across the Czech Republic | 425 | 478,177 | 138 | - | - | [119,120] |
| England & Wales | Pre-PCV | July 1996 – June 2006 | Unvaccinated healthy children | Unvaccinated children <5 | 3,752 | 3,091,000 | 461 | 48,702,414 | 1,876 | [121] |

| | | | | | | | | | | |
|---|---|---|---|---|---|---|---|---|---|---|
| | | | <5 years old in Hertfordshire | years old across and predominantly unvaccinated individuals >4 years old across England & Wales | | | | | | |
| Goroka | Pre-PCV | 1981 – 1987 | Unvaccinated children attending clinics near Goroka Town | Unvaccinated children with IPD in Goroka hospital | 2,844 | 96,207 | 56 | - | - | [30,122] |
| Morocco | Pre-PCV | November 2010 – December 2011 | Unvaccinated healthy children in Rabat | Children hospitalised with severe pneumonia in Rabat <5 years old | 200 | 212,566 | 118 | - | - | [123,124] |
| Netherlands | Pre-PCV | June 2004 – May 2006 | Unvaccinated healthy children <2 years old in Noord-Holland, Zuid-Holland and Utrecht | Children <5 years with IPD across Netherlands | 321 | 250,924 | 100 | - | - | [125,126] |
| Ontario | Pre-PCV | 1995 | Unvaccinated healthy children primarily <4 years old in Toronto | Unvaccinated children <18 years old with IPD in Toronto and surrounding area | 1,139 | 580,507 | 89 | - | - | [127] |
| Portugal | Pre-PCV | January 2001 – December 2003 | Unvaccinated healthy children <7 years old in Lisbon and Oeiras | Unvaccinated children (<18 years) and adults (>18 years) across Portugal | 1,170 | 2,071,223 | 90 | 8,284,894 | 378 | [54,128] |

| | | | | | | | | | | |
|---|---|---|---|---|---|---|---|---|---|---|
| Stockholm | Pre-PCV | 1997 | Unvaccinated healthy children <7 years old in Stockholm | Unvaccinated adults (and a small number of children) hospitalised with IPD in Stockholm | 611 | - | - | 2,004,152 | 273 | [129,130] |
| Atlanta | Post-PCV7 | June 2008 – May 2009 | PCV7-vaccinated sick Georgia-resident children <5 years old | PCV7-vaccinated children <5 years old with IPD in Atlanta | 451 | 298,831 | 47 | - | - | [115] |
| Barcelona | Post-PCV7 | 2007 – 2011 | PCV7-vaccinated healthy children <7 years old in Barcelona | PCV7-vaccinated children <7 years old with IPD in Barcelona | 209 | 228,000 | 159 | - | - | [131] |
| Bogota | Post-PCV7 | June – November 2011 | PCV7-vaccinated healthy children <18 months old in Bogota | IPD in children <2 years old in Bogota | 246 | 357,200 | 91 | - | - | [116] |
| France | Post-PCV7 | January 2008 – December 2009 | PCV7-vaccinated healthy children <2 years old across France | PCV7-vaccinated children with IPD <2 years old across France | 1,212 | 838,866 | 388 | - | - | [132,133] |
| Massachusetts | Post-PCV7 | 2001 – April 2009 | PCV7-vaccinated children visiting physicians <7 years old across Massachusetts | PCV7-vaccinated children <7 years with IPD old across Massachusetts | 2,969 | 820,000 | 206 | - | | [134] |
| Navajo | Post-PCV7 | March 2006 - March 2008 | PCV7-vaccinated Navajo or White Mountain | Active IPD surveillance of PCV7-vaccinated | 6,541 | 65,048 | 132 | 201,553 | 514 | [39,41,135] |

| | | | | | | | | | | |
|---|---|---|---|---|---|---|---|---|---|---|
| | | | Apache native American children <9 years old | Navajo or White Mountain Apache native American children <7 years old and people at least 18 years old | | | | | | |
| Netherlands | Post-PCV7 | June 2008 – May 2012 | Vaccinated healthy children <2 years old in Noord-Holland, Zuid-Holland and Utrecht | IPD in children (<5 years) across Netherlands | 660 | 232,251 | 73 | - | - | [125,126] |
| France | Post-PCV13 | January 2012 – December 2013 | PCV13-vaccinated healthy children <2 years old across France | PCV13-vaccinated children with IPD <2 years old across France | 1,212 | 842,076 | 181 | - | - | [132,133] |
| Netherlands | Post-PCV10 | June 2012 – May 2016 | Vaccinated healthy children <2 years old in Noord-Holland, Zuid-Holland and Utrecht | IPD in children (<5 years) across Netherlands | 659 | 222,671 | 47 | - | - | [125,126] |

**Table S1**: Study populations used for the analysis of serotype invasiveness in children and adults.

| Model | ELPD difference | ELPD difference standard error |
|---|---|---|
| study-adjusted type-specific Poisson | 0.00 | 0.00 |
| study-adjusted type-specific negative binomial | -13.50 | 14.55 |
| type-specific negative binomial | -190.51 | 23.02 |
| study-adjusted null negative binomial | -239.41 | 25.27 |
| null negative binomial | -288.29 | 24.84 |
| study-adjusted null Poisson | -917.07 | 111.26 |
| type-specific Poisson | -950.42 | 144.80 |
| null Poisson | -1931.64 | 223.91 |

**Table S2** Model fit comparison with leave-one-out cross-validation (LOO-CV) using the child serotype data on carriage and disease. These values were generated using the logarithm of the likelihoods calculated within the models. Models are ranked by their expected log pointwise predictive density (ELPD), calculated from the individual pointwise log predictive densities across all observed data points. The ELPD difference column shows the difference between the ELPD of a model and that of the most likely model (this value is zero for the first row, corresponding to the most likely model given the data). The ELPD difference standard error is calculated from the distribution of individual pointwise log predictive densities from the same comparison.

| Model | ELPD difference | ELPD difference standard error |
|---|---|---|
| study-adjusted type-specific negative binomial | 0.00 | 0.00 |
| study-adjusted type-specific Poisson | -15.26 | 11.12 |
| type-specific negative binomial | -162.73 | 12.55 |
| study-adjusted null negative binomial | -201.65 | 15.53 |
| null negative binomial | -253.69 | 15.42 |
| type-specific Poisson | -815.05 | 96.09 |
| study-adjusted null Poisson | -825.44 | 116.27 |
| null Poisson | -1741.04 | 191.89 |

**Table S3** Model fit comparison with LOO-CV using the child serotype data on carriage and disease. These values were generated using the logarithm of the likelihoods calculated for the observations of isolates from disease only, as these were more constrained than the modelling of isolate counts from carriage. The table is displayed as described for Table S2.

| Simulated model | Best-fitting model | Second best-fitting model | Log(Bayes factor) relative to most likely model |
|---|---|---|---|
| null Poisson | null Poisson | null negative binomial | -26.32 |
| null negative binomial | null negative binomial | type-specific negative binomial | -21.24 |
| type-specific Poisson | type-specific Poisson | type-specific negative binomial | -25.15 |
| type-specific negative binomial | type-specific negative binomial | study-adjusted type-specific negative binomial | -25.37 |
| study-adjusted null Poisson | study-adjusted null Poisson | study-adjusted null negative binomial | -41.09 |
| study-adjusted null negative binomial | study-adjusted null negative binomial | study-adjusted type-specific negative binomial | -26.80 |
| study-adjusted type-specific Poisson | study-adjusted type-specific Poisson | study-adjusted type-specific negative binomial | -33.23 |
| study-adjusted type-specific negative binomial | study-adjusted type-specific negative binomial | study-adjusted null negative binomial | -26.87 |

**Table S4** Comparison of model fits to simulated data using bridge sampling. Each row corresponds to data simulated from the specified model. All eight models were fitted to each set of simulated data. The model fits were compared used bridge sampling. The table shows the models adjudged to be the first and second best-fitting to each dataset, using bridge sampling. The final column shows the logarithm of the Bayes factor by which the best-fitting model was favoured over the second best-fitting model.

| Model | Log(Bayes factor) relative to most likely model |
|---|---|
| study-adjusted type-specific negative binomial | 0.00 |
| study-adjusted type-specific Poisson | -30.92 |
| type-specific negative binomial | -96.18 |
| study-adjusted null negative binomial | -119.47 |
| null negative binomial | -142.10 |
| study-adjusted null Poisson | -793.66 |
| type-specific Poisson | -840.98 |
| null Poisson | -1838.49 |

**Table S5** Comparison of model fits to child carriage and disease serotype data using Bayes factors calculated with bridge sampling. Models are ranked by their logarithmic marginal likelihoods. The logarithmic Bayes factors are calculated for each model relative to the most likely given the data; hence the value is zero for the first row.

| Model | Log(Bayes factor) relative to most likely model |
|---|---|
| study-adjusted type-specific negative binomial | 0.00 |
| study-adjusted type-specific Poisson | -17.03 |
| study-adjusted null negative binomial | -17.54 |
| null negative binomial | -28.38 |
| type-specific negative binomial | -36.37 |
| type-specific Poisson | -470.82 |
| study-adjusted null Poisson | -945.88 |
| null Poisson | -1814.07 |

**Table S6** Comparison of model fits to child carriage and adult disease serotype data using Bayes factors calculated with bridge sampling. The table is displayed as described for Table S5.

| Population | Vaccine period | Carriage study time interval | Carriage isolate source | Disease isolate source | No. of swabs | Population of children | No. disease isolates from children | Population of adults | No. disease isolates from adults | References |
|---|---|---|---|---|---|---|---|---|---|---|
| Finland | Pre-PCV | 1994 - 1996 | Unvaccinated children <2 years old in Tampere | Unvaccinated children <2 years across Finland | 329 | 120,238 | 143 | - | - | [74,136] |
| Oxford | Pre-PCV | 1994 - 2001 | Unvaccinated healthy children <5 years old in Oxford | Unvaccinated children <5 years old with IPD in Oxford | 639 | 37,467 | 84 | - | - | [31,137] |
| Portugal | Pre-PCV | January 2001 - December 2003 | Unvaccinated healthy children <7 years old in Lisbon and Oeiras | Unvaccinated children (<18 years) and adults (>18 years) across Portugal | 1,170 | 2,071,223 | - | 8,284,894 | 152 | [54,128] |
| Stockholm | Pre-PCV | 1997 - 2004 | Unvaccinated children <7 years old attending day care centres in Stockholm County | Unvaccinated children <18 years old from the Stockholm area | 1,330 | 397,289 | 65 | - | - | [78,110,111, 129] |
| South Africa | Post-PCV7 | 2009 - 2010 | HIV negative children <13 years old in Soweto and Agincourt | HIV negative children <7 years old in South Africa | 2,674 | 7,187,314 | 515 | - | - | [37,101,103, 104] |
| USA | Post-PCV7 | 2006 - 2009 | Children <7 years of age in Massachusetts | Children <7 years old in the USA ABCS regions | 1,983 | 1,931,331 | 405 | - | - | [37,93,106,1 38,139] |

| | | | | | | | | | | |
|---|---|---|---|---|---|---|---|---|---|---|
| South Africa | Post-PCV13 | 2011 - 2013 | HIV negative children <13 years old in Soweto and Agincourt | HIV negative children <7 years old in South Africa | 2,023 | 7,329,006 | 340 | - | - | [37,101,103, 104] |

**Table S7**: Study populations used for the joint analysis of strain and serotype invasiveness. The disease isolates from Portugal came from a mixture of infants and adults, but are tabulated based on them primarily arising from the latter age category.

| Model | ELPD difference | ELPD difference standard error |
|---|---|---|
| type-specific strain-modified Poisson | 0.00 | 0.00 |
| strain- and serotype-specific Poisson | -1.56 | 3.57 |
| strain-specific type-modified Poisson | -4.77 | 3.50 |
| type-specific strain-modified negative binomial | -26.97 | 3.34 |
| strain-specific type-modified negative binomial | -37.88 | 4.89 |
| type-specific negative binomial | -41.08 | 5.91 |
| strain-specific Poisson | -43.61 | 18.76 |
| type-specific Poisson | -45.21 | 14.48 |
| strain- and type-specific negative binomial | -48.23 | 6.02 |
| strain-specific negative binomial | -61.00 | 6.81 |

**Table S8** Model fit comparison with LOO-CV using the child strain and serotype data on carriage and disease. These values were generated using the logarithm of the likelihoods calculated for the observations of isolates from carriage and disease. The table is displayed as described for Table S2.

| Model | ELPD difference | ELPD difference standard error |
|---|---|---|
| type-specific strain-modified Poisson | 0.00 | 0.00 |
| strain- and type-specific Poisson | -0.57 | 2.66 |
| strain-specific type-modified Poisson | -3.29 | 2.34 |
| type-specific strain-modified negative binomial | -21.14 | 2.48 |
| type-specific negative binomial | -32.08 | 5.14 |
| strain-specific type-modified negative binomial | -32.94 | 4.17 |
| strain-specific Poisson | -36.85 | 15.83 |
| strain- and type-specific negative binomial | -41.17 | 5.35 |
| type-specific Poisson | -44.04 | 13.05 |
| strain-specific negative binomial | -51.28 | 5.63 |

**Table S9** Model fit comparison with LOO-CV using the child strain and serotype data on carriage and disease. These values were generated using the logarithm of the likelihoods calculated for the observations of isolates from disease only. The table is displayed as described for Table S2.

| Model | Log(Bayes factor) relative to best-fitting model |
| --- | --- |
| type-specific strain-modified Poisson | 0.00 |
| type-specific negative binomial | -5.59 |
| type-specific strain-modified negative binomial | -5.99 |
| type-specific Poisson | -24.61 |
| strain-specific type-modified Poisson | -25.27 |
| strain-specific serotype-modified negative binomial | -29.98 |
| strain- and type-specific Poisson | -33.29 |
| strain-specific negative binomial | -33.94 |
| strain- and type-specific negative binomial | -37.33 |
| strain-specific Poisson | -49.46 |

**Table S10** Comparison of model fits to strain and serotype data from child carriage and disease using Bayes factors calculated with bridge sampling. The table is displayed as described for Table S5.

| Model | Log(Bayes factor) relative to best-fitting model |
|---|---|
| type-specific strain-modified Poisson | 0.00 |
| type-specific negative binomial | -5.63 |
| type-specific strain-modified negative binomial | -5.94 |
| type-specific Poisson | -24.70 |
| strain-specific type-modified Poisson | -25.28 |
| strain-specific serotype-modified negative binomial | -29.67 |
| strain- and type-specific Poisson | -32.86 |
| strain-specific negative binomial | -33.49 |
| strain- and type-specific negative binomial | -36.48 |
| strain-specific Poisson | -49.40 |

**Table S11** Comparison of model fits to strain and serotype data from child carriage and disease using Bayes factors calculated with bridge sampling. In this analysis, the carriage sample size for three studies (Finland pre-PCV, Oxford pre-PCV and Stockholm pre-PCV) was increased 100-fold, to test for the sensitivity of the model comparisons to the uncertainty in this parameter (Text S2). The table is displayed as described for Table S5.

| Previous publication | Serotype | Low invasiveness genotype | Low invasiveness strain | High invasiveness genotype | High invasiveness strain | Results from this study |
|---|---|---|---|---|---|---|
| Sá-Leão *et al* 2011 | 3 | ST180 | GPSC12 | ST156 | GPSC6 | GPSC6 was estimated to have higher invasiveness, albeit with overlapping credibility intervals |
| Gladstone *et al* 2019 | 6A | CC172 | GPSC5 | CC1094 | GPSC41 | GPSC41 invasiveness was found to be significantly higher than GPSC5 |
| Sá-Leão *et al* 2011 | 6A | ST460, ST315, ST1879 | GPSC47, GPSC64, ST1879 | - | - | Other GPSCs of this serotype were not tested in adults |
| Sá-Leão *et al* 2011 | 6B | ST176, ST315 | GPSC24, GPSC47 | - | - | Other GPSCs of this serotype were not tested in adults |
| Hanage *et al* 2005 | 6B | - | - | ST138 | GPSC24 | GPSC24 was found to be more invasive than other 6B GPSCs |
| Hanage *et al* 2005 | 7F | - | - | ST191 | GPSC15 | Not tested |
| Sá-Leão *et al* 2011 | 11A | ST62 | GPSC3 | - | - | Not tested in adults |
| Hanage *et al* 2005 | 11A | ST62 | GPSC3 | - | - | Estimated to have similar invasiveness as GPSC22 |
| Gladstone *et al* 2019 | 14 | CC63 | GPSC9 | CC15 | GPSC18 | GPSC18 invasiveness was estimated to be higher than that of GPSC9, albeit with slightly overlapping credibility intervals |
| Sá-Leão *et al* 2011 | 14 | - | - | ST156 | GPSC6 | GPSC6 was found to be more invasive than GPSC9, albeit with overlapping credibility intervals, in adults |
| Hanage *et al* 2005 | 14 | - | - | ST156 | GPSC6 | GPSC6 was found to be more invasive than GPSC9, albeit with overlapping credibility intervals |
| Brueggemann *et al* 2003 | 14 | - | - | ST9 | GPSC18 | GPSC18 was found to be more invasive than GPSC9, albeit with overlapping credibility intervals |
| Brueggemann *et al* 2003 | 14 | - | - | ST124 | GPSC39 | GPSC39 was estimated to be more invasive than GPSC9, albeit with overlapping credibility intervals |

| | | | | | | |
|---|---|---|---|---|---|---|
| Hanage *et al* 2005 | 14 | - | - | ST124 | GPSC39 | GPSC39 was estimated to be more invasive than GPSC9, albeit with overlapping credibility intervals |
| Gladstone *et al* 2019 | 16F | CC4088 | GPSC33 | CC30 | GPSC46 | Not tested |
| Brueggemann *et al* 2003 | 18C | - | - | ST113 | GPSC50 | Estimated invasiveness is similar to that of GPSC67 and GPSC68 |
| Gladstone *et al* 2019 | 19F | CC347 | GPSC21 | CC320 | GPSC1 | GPSC1 invasiveness was estimated to be significantly higher than that of GPSC21 |
| Sá-Leão *et al* 2011 | 19F | ST177 | GPSC44 | - | - | Not tested in adults |
| Hanage *et al* 2005 | 19F | ST485 | GPSC175 | - | - | Low invasiveness, with overlapping credibility intervals with other GPSCs |
| Sá-Leão *et al* 2011 | 19A | ST81 | GPSC16 | ST193, ST230 | GPSC11, GPSC10 | Not tested in adults |
| Hanage *et al* 2005 | 19A | - | - | ST482 | ST482 | Higher invasiveness than GPSC17, albeit with overlapping credibility intervals |
| Sá-Leão *et al* 2011 | 22F | - | - | ST443 | ST443 | Not tested in adults |
| Gladstone *et al* 2019 | 23B | CC439 | GPSC7 | CC172 | GPSC5 | Not tested, although GPSC5 was found to be more invasive that GPSC7 in serotype 23A |
| Sá-Leão *et al* 2011 | 23F | ST176, ST439 | GPSC24, GPSC7 | - | - | GPSC7 was estimated to have lower invasiveness than GPSC5 in adults, albeit with overlapping credibility intervals |
| Sá-Leão *et al* 2011 | 34 | ST1439 | GPSC45 | - | - | Not tested in adults |

**Table S12** Comparison of previously-identified within-serotype differences in strain invasiveness with the results of this analysis. The exclusion of strain and serotype combinations represented by fewer than ten isolates, summed across carriage and disease, meant that some within-serotype tests between strains were not undertaken in this analysis.

**Supplementary Datasets**

**Dataset S1** Invasiveness estimates for serotypes in children. The median estimates, and lower and upper bounds of the 95% credibility interval, are listed for each serotype.

**Dataset S2** Invasiveness estimates for serotypes in adults. The median estimates, and lower and upper bounds of the 95% credibility interval, are listed for each serotype.

**Dataset S3** Invasiveness estimates for serotypes and strains, estimated from genotyped isolates from child disease and carriage. Each row corresponds to a serotype-strain combination. Both the invasiveness associated with the serotype, and the invasiveness coefficient associated with the strain, are listed, along with the corresponding lower and upper bounds of their 95% credibility intervals.
